# Cell identities along the proximal-distal and micropylar-chalazal axes in the Arabidopsis heart-stage seed

**DOI:** 10.64898/2026.07.27.741042

**Authors:** Khadija L. B. Rombi, Thomas Hartwig, Nora R. Zöllner, Tin Yau Pang, Martin J. Lercher, Michael M. Wudick, Wolf B. Frommer, Ji Yun Kim

## Abstract

• Seeds are complex reproductive organs consisting of diverse maternal and filial tissues. During development, the embryo and specialized tissues for nutrient storage required for seed germination and early seedling establishment emerge.

• To explore the cellular diversity and differentiation of seeds, we performed single cell RNA-sequencing on heart stage Arabidopsis seeds and identified 20,097 cells that were grouped into ≥21 distinct cell clusters. 20 of the 21 clusters were spatially assigned by combining bioinformatic analysis, imaging reporter fusion marker lines, and spatial transcriptomics.

• Our analysis revealed a high degree of differentiation of epidermal cell and inner cell layers along the rotational and axial seed axes, highlighting the importance of cell position and ontogenesis. We identified unexpected spatial domains, including a cluster marked by abscission zone-specific transcripts, and a nucellar cluster shaped by developmentally programmed cell death. Surprisingly, embryo and endosperm showed similarities in transcript profiles despite distinct and complementary functions.

• In summary, our findings establish seeds as a transcriptionally complex organ with high cell type heterogeneity and provide a basis for investigating the differentiation of diverse cell layers and spatial transcript profiles.

## Introduction

Classically, the cell type composition of tissues and organs had been defined based on light microcopy. More recently, *in situ* hybridization and promoter-reporter fusions provided insights into the transcriptional outfit of individual cell types and enabled the identification of novel cell types. Single cell RNA-sequencing (scRNA-seq) and spatial transcriptomics have revolutionized the field, providing new perspectives on the role of specific cell types and their developmental trajectories. In plants, the first scRNA-seq studies focused on root systems, characterizing cell type identities and trajectories in roots (Denyer et al. 2019; Ryu et al. 2019; Shahan et al. 2022; Zhi et al. 2023). This technology has then been applied to leaves enabling the characterization of cell types lacking established markers, such as the phloem parenchyma cells in leaves, and distinct cellular states including pathogen-responsive founder cells (Kim et al. 2021; Nobori et al. 2025). Furthermore, scRNA-seq combined with spatial validation revealed differentiation of maize leaf bundle sheath cells and identified stage-specific activation of sequence-conserved ancient genes during embryogenesis (Bezrutczyk et al. 2021; Wu et al. 2025). While roots and leaves have been shown to contain a fundamental set of 8 -10 cell types plus cells in different developmental stages (Denyer et al. 2019), we hypothesize that seeds, exhibit a higher cell type complexity due to their developmental programs involving coordinated differentiation of the maternal, embryonic, and endosperm tissues. Seeds are key reproductive organs of plants, deriving from the double fertilization of the maternal ovule. During fertilization, one sperm nucleus fuses together with the central cell of the ovule to form the triploid endosperm of the seed, while the other sperm nucleus fuses with the egg cell to form the diploid zygote. Coordinately, upon fertilization, the maternal ovule integuments differentiate into the seed coat integuments (Haughn and Chaudhury 2005). The seed coat has traditionally been regarded as a simple unit that encloses the developing embryo and endosperm. Anatomically, the seed coat is composed of two distinct integuments: an outer and an inner integument. The outer integument consists of two cell layers, termed outer integument 1 (oi1) and outer integument 2 (oi2) (Figure 1a). The inner integument consists of two to four cell layers, including the inner integument 1 (ii1), also known as endothelium, the inner integument 1’ (ii1’), which develops from periclinal cell divisions of the ii1 and the inner integument 2 (ii2). While the cell layers at the distal seed coat (distal to the funiculus) are clearly identified, the proximal region (proximal to the funiculus) is collectively referred to as the chalazal seed coat, and its cellular heterogeneity remains largely unexplored (Schneitz et al. 1995; Debeaujon et al. 2003). The products of double fertilization, the endosperm and the embryo, form the central tissues in the seed and are characterized by a complex developmental program defined by morphological and transcriptional changes.

**Figure 1:**
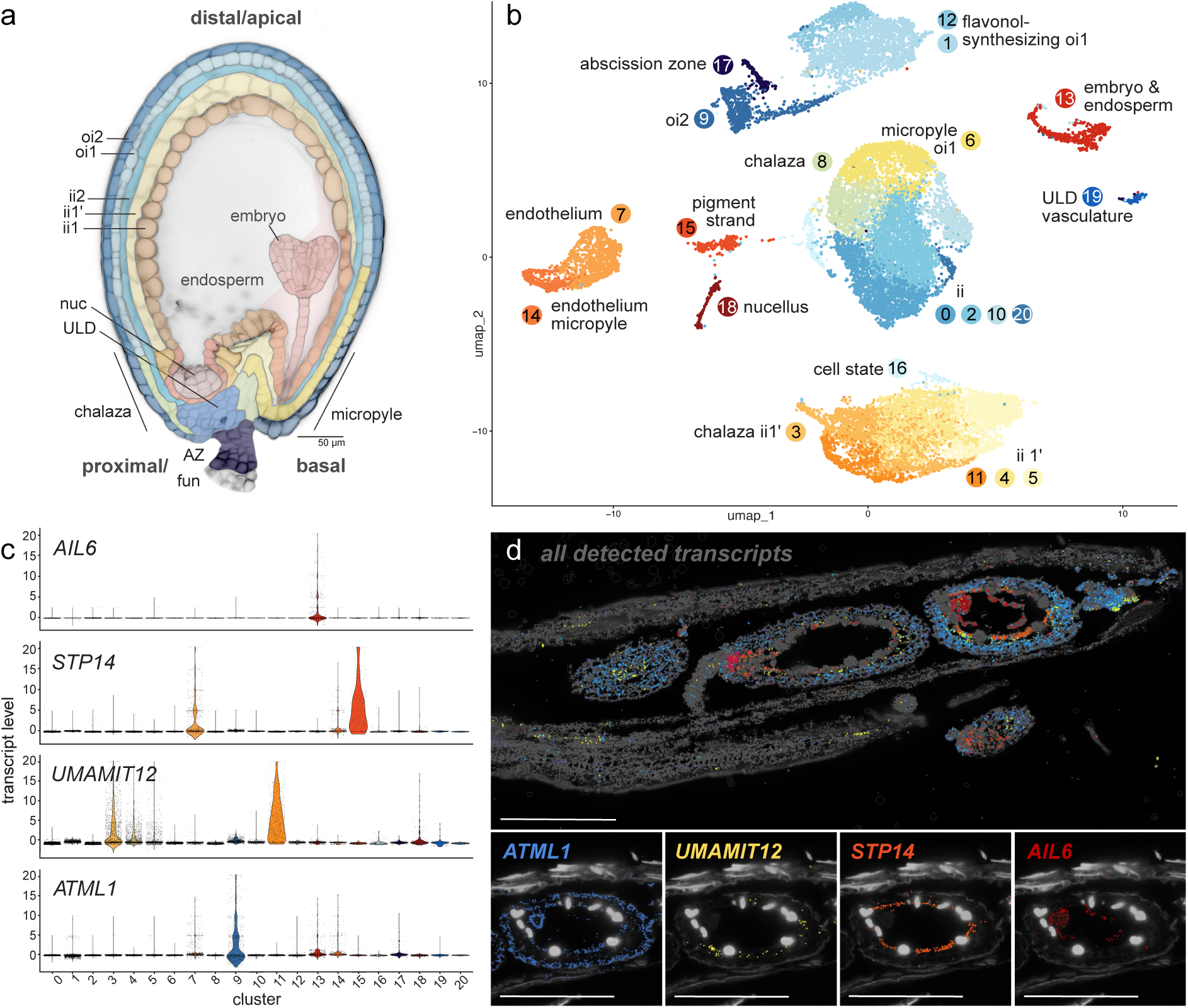
Spatially validated scRNA-seq atlas revealing transcriptional complexity of the developing seed. (a) A representative Arabidopsis seed at heart stage of embryo development with cell layers and spatial domains along the proximal (to funiculus)/basal, distal (to funiculus)/apical and micropylar-chalazal axes, false-colored corresponding to scRNA-seq cluster annotation. oi2, outer integument 2/adaxial epidermis; oi1, outer integument 1/abaxial epidermis; ii2, inner integument 2; ii1’, inner integument 1’; ii1, inner integument 1/endothelium; nuc, nucellus; ULD, unloading domain; AZ, abscission zone; fun, funiculus. (b) UMAP plot of integrated scRNA-seq datasets with cluster annotations. (c) Violin plots showing enrichment for select marker transcripts detected in the spatial transcriptomics dataset. (d) MERFISH spatial transcriptomics showing a silique enclosing three developing seeds sectioned at different seed plane, with all detected transcripts visualized in grey, select transcripts color coded (see Supporting Information Fig. S3, Supporting Information Table S1). Zoom on seed with select transcript visualizations corresponding to marker violin plots in (c). Scale bars 250 µm.

Recent single nucleus RNA-seq (snRNA-seq) analysis profiled early Arabidopsis seed development with focus on the endosperm and imprinting complexity (Picard et al. 2021; Martin et al. 2026). An independent single nucleus transcriptomic atlas spanning the Arabidopsis life cycle, including multiple organs including seeds, revealed eight distinct clusters, half of which could be assigned to seed tissues, two of which were assigned to outer and inner integuments (Lee et al. 2025). Beyond Arabidopsis, a spatially resolved multi-omics single nucleus atlas has been developed to resolve transcriptome profiles and chromatin accessibility during soybean seed development (Zhang et al. 2025), as well as a four-dimensional spatial transcriptome atlas resolving barley grain development (Peirats-Llobet et al. 2026)

In this study, we aimed to capture a near complete set of cellular transcriptomes of Arabidopsis seeds at the heart stage of embryo development and spatially map the transcript profiles of at least 20 cell types and states. Our analysis revealed surprising findings, including similarities between transcript profiles of embryo and endosperm despite distinct functions and clear differentiation of the epidermal cell layers along both apical-basal and radial axes. We also identified a cluster representing the seed abscission zone. Notably, the integuments fall into 16 distinct clusters. Our seed scRNA-seq atlas provides a basis for exploring differentiation of the diverse cell layers, early specification of the seed abscission zone, characterization of the cells involved in specialized metabolic pathways and the spatial domains along the proximal-distal seed axes.

## Material and Methods in brief

Detailed materials and methods are available in Supporting Information Note 1.

### Plant growth and transgenic plant material

Arabidopsis plants were grown in soil in long day conditions (8 h dark, 16 h light) at a PAR of 100 µmol m^−2^ s^−1^, 23-24 °C (day), 20-21 °C (night) and 70% humidity. Plant material for scRNA-seq and spatial transcriptomics was harvested 4-6 h after light exposure. Transgenic material used in this study can be found in Supporting Information Note 1.

### Enrichment of seed protoplasts

For scRNA-seq experiments, Five-weeks-old Arabidopsis Col-0 plants were used to isolate protoplasts from seeds corresponding to the heart stage of embryo development. Detailed methods are described in Supporting Information Note 1.

### ScRNA-seq and data analysis

For single-cell RNA sequencing, an approximate count of 20.000 viable protoplasts (620 µL of protoplast suspension) was loaded into the BD Rhapsody cartridge and processed according to the BD Rhapsody user guide (BD Rhapsody 2019). Raw fastq files were mapped to the Arabidopsis Col-0 genome from TAIR10, Araport11 using the BD Rhapsody pipeline version 2.2.1 with default parameters. Quality control and preprocessing were performed according to a single cell best practices guide (Heumos et al. 2023). Visualization, marker analysis, subclustering analysis, gene set enrichment, GO term analysis were performed using an R studio Seurat environment. R code is available in Supporting Information Note 2. Pathway activity scores, depending on transcript levels of respective constitutive genes, were calculated following Xiao et al. 2019 and Kim et al. 2021 using the AraCyc pathways 17.2.0. as described in Supporting Information Note 1.

### Spatial transcriptomics

For use in MERFISH spatial transcriptomics, Arabidopsis Col-0 siliques with seeds in embryonal heart stage were processed according to MERSCOPE® User Guide, 91600112 Rev C. After tissue processing, microtome sections containing longitudinal silique sections were microscopically selected prior to mounting on MERFISH slides. Slides were stored at –20 °C prior to internal processing at Vizgen headquarters (Cambridge, USA). Spatial transcriptomics data was visualized in the MERSCOPE Visualizer software.

### Confocal imaging

To enable the detection of fluorescence with cellular resolution in deeper cell layers of developing Arabidopsis seeds, seeds were subjected to optical clearing, adapted from Kurihara et al. 2021; Attuluri et al. 2022. Whole seeds were stained with SR2200; protoplast suspensions were either imaged directly or stained. Arabidopsis seeds, siliques, and protoplasts were mounted onto slide glasses and analyzed by CLSM. Detailed methods for seed clearing, staining and microscope settings can be found in Supporting Information Note 1.

## Data availability

The raw and processed sequencing data are available at Gene Expression Omnibus (GEO) (www.ncbi.nlm.nih.gov/geo/) under accession number X.

## Results

### Marker-assisted optimization of single-cell isolation from developing seeds

scRNA-seq has been considered to have distinct advantages compared to snRNA-seq, such as higher sensitivity and capture of cytoplasmic RNA (Guillotin et al. 2023). Despite these advantages, a major drawback limiting the application of scRNA-seq in plants is the necessity of generating a single-cell population that reflects all cell types of the tissue of interest (Denyer and Timmermans 2022). While standard protocols have been adapted to efficiently isolate protoplasts from various tissues including root, leaf, and hypocotyls (Denyer et al. 2019; Kim et al. 2021; Zhang et al. 2021), marker-based optimization of the protocols was required to obtain protoplasts representing all cell types of a given organ. To develop a methodology that ensures representation of all tissues of developing Arabidopsis seeds, we searched for markers that reflect the broad spectrum of cellular characteristics within the cell. Diverging cellular characteristics in the seed include cell size (ranging in diameter from 3-10 µm in the unloading domain (ULD) to 30-60 µm in the endothelium), cell wall composition (from pectin-rich oil cells to tannin-containing endothelial cells), starch content, as well as depth within the seed (Beeckman et al. 2000; Debeaujon et al. 2001). Morphological features, such as pigment bodies and starch granules were used to identify the endothelium and the outer integument cell layers, respectively, in protoplast preparations (Figure 1a, Supporting Information Fig. S1). In addition, we used transcriptional or translational GFP-reporter fusions marking distinct cell populations, such as *pbZIP9:GFP-GUS* for the unloading domain and *pSWEET12:SWEET12-eGFP* marking the micropylar integuments. To avoid contamination with silique and funiculus tissues, we manually collected individual seeds from opened siliques. To cover a developmental stage in which both tissue differentiation and tissue elimination are balancing, seeds at the early to late heart stage of embryo development were used. Individual isolation of seeds enabled precise control over the developmental stage of the embryo based on morphological features such as a partial greening of the micropyle, seed size and shape (Supporting Information Fig. S1). The developmental stage of seeds was validated by visually assessing the incompletely digested embryos in the protoplasting suspension prior to washing steps. To facilitate penetration of cell wall–digesting enzymes through the multiple waxy cuticles between seed cell layers, we ruptured the seed coats using forceps during isolation of individual seeds. Seed coats were further mechanically disrupted by shearing the seed suspension through a pipette tip. Pectolyase was included in the protoplasting solution to digest pectin-rich seed coat cell walls. Since the protoplasts of major and diverse cell types could be released successfully (Supporting Information Fig. S1), the optimized marker-assisted protoplast isolation protocol was used to generate protoplasts for scRNA-seq.

### Spatial assignment of distinct cell-type clusters in developing Arabidopsis seeds

scRNA-seq libraries were generated from two independent replicates and sequenced to approximately 53.000 reads per cell. After removing low-quality cells and doublets using CellBender and Seurat, respectively, 20.097 cells with a mean of 2.4 k genes and 10.9 k transcripts per cell were obtained. To ensure reproducibility, a second independent repeat was performed. Comparison of the independent replicates revealed a similar distribution of cell identities (Supporting Information Fig. S2). Replicates were integrated using the linear Harmony algorithm to remove batch effects. Plotting the single-cell transcriptomes in two dimensions using UMAP revealed at least 21 distinct clusters (Figure 1b). Developing Arabidopsis seeds comprise three genetically distinct domains; the diploid embryo and the triploid endosperm as the products of double fertilization, as well as five maternal seed coat cell layers, the nucellus and the chalazal seed coat. To assign seed tissue identities to the clusters, we analyzed the enrichment of transcripts of genes known to exhibit tissue or cell type-specific enrichment, as well as fluorescent reporter lines, and used spatial transcriptomics as a complementary approach (Figure 1c-e, Supporting Information Table S1, S3, Fig. S1, S3). From these analyses we were able to identify all nine major canonical “cell types/tissues” of the seed. Embryonal and endosperm cell types made up 2.7%, nucellar 1%, vascular cell types 0.6% of single cell transcriptomes. Despite the seven canonically described seed coat tissues, we surprisingly identified 16 seed coat clusters, which accounted for 93.5% of single cell transcriptomes. We therefore explored whether the unexpected transcriptional complexity of the seed coat clusters reflected spatial domains, distinct cell states, or cell types. We first analyzed the transcriptional signatures of single-cell clusters from the conventionally well-characterized five cell layers of the distal seed coat (Figure 2a-b), beginning with the epidermis.

**Figure 2:**
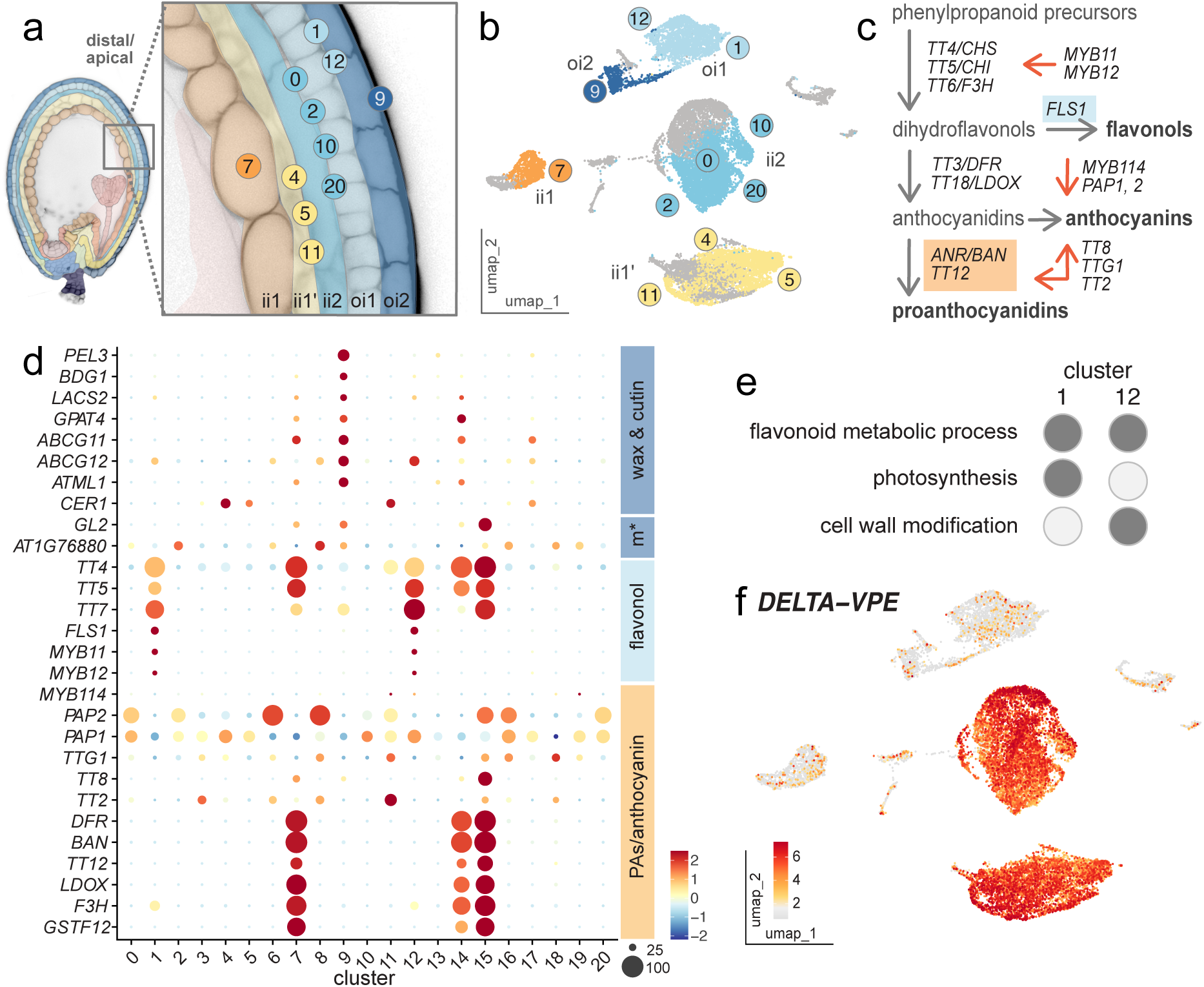
Defining distal seed coat cell layers by specialized metabolites and developmental progression. (a) Close-up of a representative Arabidopsis distal/apical seed coat with five cell layers false-colored corresponding to scRNA-seq cluster annotation, marked on a cell layer level in (b) UMAP plot of integrated scRNA-seq datasets. (c) Flavonoid biosynthesis pathway (adapted from Nesi et al. 2001; Li 2014) transcripts are detected in (d) clusters corresponding to distinct cell layers, together with wax, cutin- and mucilage-associated (m*) transcripts. Dot size indicates percent of cells in which transcript was detected, color indicates log-fold enrichment, legend on the right, (e) The two oi1 clusters, 1 and 12, are distinguished by photosynthesis and cell wall modification while sharing flavonoid metabolism in a comparative GO term enrichment analysis. Representative GO terms were selected from the three highest adjusted p-value GO terms (Supporting Information Table S2, code in Supporting Information Note 2). (f) The ii2 and ii1’ are marked by the programmed cell death associated vacuolar processing enzyme transcripts *DELTA-VPE*, illustrated in a UMAP feature plot. Color indicates log-fold enrichment, legend on the left. Cell types and transcripts derived from them are color-coded consistently throughout the figure as established in (a).

### Distinct specialized metabolites and cell states of the epidermal clusters

The seed epidermis is morphologically composed of two distinct cell layers: the outer integument 2 (oi2), also termed adaxial epidermis, and the outer integument 1 (oi1), also termed abaxial epidermis (Figure 1b, 2a-b). Each of the one-cell-thick layers is characterized by distinct functions with unique metabolite profiles (Figure 2c). In the oi2, a mucilage layer is deposited between the heart and torpedo stage of embryo development, likely involved in seed dispersal and germination (Western et al. 2000; Windsor et al. 2000). Additionally, a cuticle layer is formed, implicated in prevention of postgenital organ fusion (Watanabe et al. 2004; Panikashvili et al. 2009). Transcripts related with cutin, such as *DEFECTIVE IN CUTICULAR RIDGES (DCR/PEL3)*, *BODYGUARD1 (BDG1)*, *LONG-CHAIN ACYL-COA SYNTHETASE 2* (*LACS2*), *GLYCEROL-3-PHOSPHATE sn-2-ACYLTRANSFERASE 4* (*GPAT4*), and transcripts of genes involved in apoplasmic transport of cutin and wax precursors (*ARABIDOPSIS THALIANA WHITE-BROWN COMPLEX HOMOLOG PROTEIN 11* and *12* (*ABCG11*/*12))* were enriched in the 839 cells-containing cluster 9 (Figure 2d), indicating that cluster 9 represents cells with epidermal identity (Schnurr et al. 2004; Li et al. 2007; Ukitsu et al. 2007; Coen et al. 2019). The enrichment of the epidermis-specific transcription factor *ARABIDOPSIS THALIANA MERISTEM LAYER 1* (*ATML1*) and its detection by spatial transcriptomics support the identity of cluster 9 as adaxial epidermis/oi 2 (Takada et al. 2013) (Figure 1c-d, 2d). Pathway activity analysis, which quantifies the average transcript detection of defined pathways in the clusters, showed that only a small number of pathways exhibited high pathway activity scores in cluster 9, implicating functional specialization (Xiao et al. 2019). Enriched pathways included wax and cutin formation, consistent with cuticle layer formation in the epidermis (Supporting Information Fig. S4). Notably, the seed coat epidermis also undergoes a complex differentiation, including cytoplasmic rearrangement and secondary cell wall synthesis, ultimately leading to the secretion of pectinaceous mucilage (Beeckman et al. 2000). Between heart and torpedo stage of embryo development, mucilage is synthesized in oi2, and starch grains group to form columellae (Beeckman et al. 2000; Western et al. 2000). To test whether the mucilage secretory cell fate is already primed at the heart stage before becoming morphologically apparent, we mined our data for mucilage biosynthesis and secretion associated transcription factors. We identified *GLABRA2* (*GL2*), which is known to target *MUCILAGE MODIFIED 4* (*MUM4*), a rhamnose synthase required for mucilage biosynthesis, as well as *DE1 BINDING FACTOR 1* (*DF1*), a transcription factor implicated in modulating mucilage biosynthesis (Shi et al. 2012; Xu et al. 2022) (Supporting Information Fig. S5). However, mucilage biosynthesis itself was not yet detectable in our pathway activity analysis at the heart stage. Instead, the pathway analysis highlighted pathways related to growth, cell wall and membrane remodeling (Supporting Information Fig. S4), implicating that the cytoplasmic and cell wall remodeling associated processes precede mucilage biosynthesis.

The abaxial epidermal cell layer, i.e., the outer integument (oi1), is characterized by a higher number of smaller cells relative to the adaxial epidermis. Ontogenetically, oi1 and oi2 arise from an epidermal layer that divides and grows around the inner integument (ii), nucellus and ultimately encloses the female gametophyte. As they originate from the same precursor cell population, a transcriptional kinship between the oi1 and oi2 might be expected. Indeed, a trajectory of cells from the oi2 adaxial epidermal cluster 9 bridges to two joint clusters 1 and 12. Clusters 1 and 12 were assigned as oi1, as they were specifically enriched for *EUI-LIKE P450 A1* (*ELA1/CYP714A1*), which had previously been shown to localize in oi1 and ii1’, as well as *GAMMA-INTERFERON-RESPONSIVE LYSOSOMAL THIOL REDUCTASE* (*GILT*), an oi1 marker in *Brassica napus* (Wu et al. 2011; Creff et al. 2015) (Figure 2b and Supporting Information Fig. S5). Interestingly, clusters 1 and 12 were both enriched with transcripts of several *TRANSPARENT TESTA* genes associated with early steps of flavonoid biosynthesis, e.g., *TT4*, *TT5*, and *TT7* (Figure 2d). From the early biosynthesis products, dihydroflavonols, multiple compounds are synthesized, such as proanthocyanins and flavonols which exhibit cell-layer specificity in the developing seed. While proanthocyanidins accumulate in the endothelium, UV-protective flavonols are known to accumulate in the outer integument, specifically in oi1 (Pourcel et al. 2005) (Figure 2c). Consistent with the cell-layer specificity, late flavonoid biosynthetic genes such as *DFR* or *BAN* could not be detected in clusters 1 and 12 (Li 2014) (Figure 2c-d). By contrast, *FLS1*, encoding a flavonol synthase enzyme responsible for catalyzing the final step in flavonol biosynthesis, was specifically enriched in clusters 1 and 12 ((Schilbert et al. 2024) Figure 2d). Thus, clusters 1 and 12 could be assigned to flavonoid-synthesizing oi1 cells. We hypothesized that the two clusters may separate due to: (1) a developmental progression between the clusters, or (2) belong to different spatial domains within oi1. To test the hypothesis, we investigated oi1 specific markers in our spatial transcriptomic dataset, yet no apparent spatial domains were detectable. Interestingly, GO term enrichment comparison between clusters 1 and 12 revealed diverging programs. While shared GO terms were related to *flavonoid metabolism*, cluster 1 was enriched for terms related to *translation*, *respiration*, and *photosynthesis*, and cluster 12 was enriched for *cell wall modeling* and *organization* (Figure 2e, Supporting Information Table S2). During seed development, the abaxial epidermal oi1 cells undergo complex structural differentiation entailing starch granule enlargement and vacuolar fragmentation followed by shrinkage of both vacuoles and starch granules. While starch is being degraded at the torpedo stage of embryo development, the abaxial cell walls are reinforced with lignin and suberin before appearing as a thick cell wall in oi2 at maturity (Beeckman et al. 2000; Windsor et al. 2000; Hyvärinen et al. 2025). Because this process has only been described anatomically, we evaluated putative players in the developmentally programmed cell wall remodeling by examining transcript enrichment of gene families associated with cell wall biogenesis. Members of *XTH*, *CESA*, *PME*, *BGAL* families were found to be enriched in the oi1 cluster 12, as well as cluster 1 (Supporting Information Fig. S6). In line with secondary cell wall thickening in the oi1, *BXL1* transcripts, encoding a bifunctional β-D-xylosidase/⍺-L-arabinofuranosidase involved in xylan remodeling, were enriched in all epidermis clusters (Goujon et al. 2003). We further hypothesized that cell wall carbohydrates produced by cell wall degradation and recycling might be transported to the apoplasm and/or neighboring cells. Analysis of sugar transporter gene families in clusters 1 and 12 identified *SWEET7,* a uniporter for glucose and xylose, as well as *STP7,* a D-xylose and L-arabinose H^+^-symporter (Rottmann et al. 2018; Kuanyshev et al. 2021) (Supporting Information Fig. S5). Because the comparative GO term analysis showed enrichment for terms related to *respiration* and *photosynthesis* for cluster 1, we asked whether the difference between the clusters 1 and 12 might be the result of a developmental progression that coincided with a change in cell state. By mapping the percentage of plastidial transcripts, we found that cluster 1 is characterized by the highest enrichments of mitochondrial and chloroplast transcripts, with a reduced number of genes and transcripts detected per cell compared to the other clusters (Supporting Information Fig. S2). As elevated organellar transcript proportions are associated with cell death in scRNA-seq studies, cluster 1 may represent an oi1 sub-population undergoing cell death, rather than solely reflecting photosynthetic and metabolically active cells.

### Early onset of developmentally programmed cell death in the inner integument 1’ and 2

Dihydroflavonols are not only precursors for flavonols, but also proanthocyanidins (Figure 2c). While flavonoid compounds can be found in oi1, proanthocyanidins that oxidize during seed development and form visible, brown procyanidins (condensed tannins) are specific to the endothelium and the chalazamicropylar pole (Nesi et al. 2001; Pourcel et al. 2005). The biosynthesis of proanthocyanidins is well characterized and involves *TT3*, coding for a dihydroflavonol-4-reductase (DFR), a leucoanthocyanidin reductase encoded by *BANYULS* (*BAN*), and the vacuolar transporter *TT12* (Debeaujon et al. 2001; Nesi et al. 2001). *DFR*, *BAN* and *TT12* are highly enriched in several clusters, including cluster 7 (Figure 2d). Transcripts of the proanthocyanidin metabolism associated genes *LEUCOANTHOCYANIDIN DIOXYGENASE* (*LDOX*), *TT4*, *TT5*, *FLAVANONE 3-HYDROXYLASE* (*F3H*), and *GLUTATHIONE-S-TRANSFERASE* (*GSTF12*) are also enriched in cluster 7, supporting a proanthocyanidin synthesizing role (reviewed in Saito et al. 2013) (Figure 2d). Spatial transcriptomics of the highly enriched marker *STP14* confirmed endothelial identity of cluster 7 (Figure 1c-d, Supporting Information Fig. S3). The endothelium, as the innermost part of the inner integuments, interfaces the endosperm. At the abaxial interface, a cuticle layer is deposited during development (Giorgi et al. 2015; Demonsais et al. 2020). Interestingly, besides GO term and PAS features related to well-characterized endothelium properties, such as *flavonoids, cutin, fatty acids*, cluster 7 also exhibited a high enrichment for *GDP-L-fucose biosynthesis I from GDP-D-*mannose (Supporting Information Table S2). As fucose is known to play roles in cell wall composition and plasma membrane-cell wall adhesion through modification of xyloglucans and pectin, its enrichment in cluster 7 may reflect a role in reinforcing the endothelium–endosperm interface.

In contrast to the proanthocyanidin synthesizing endothelium, the cutin and mucilage synthesizing outer integument 2, and the flavonol synthesizing oi1, the two cell layers between the ii1 and oi1, the ii2 and ii1’, marked in turquoise and yellow on the UMAP, and spatially validated by *UMAMIT12* transcript detection, did not appear to be distinctly enriched for transcripts associated with specialized metabolites (Figure 1c-d, Figure 2a-b). Both cell layers were enriched for *delta-VPE*, coding for a vacuolar processing enzyme with a role in programmed cell death in the ii1’ and ii2 during early seed development (Nakaune et al. 2005) (Figure 2f). Together with the absence of a high specialized metabolism transcript enrichment and a low gene and transcript per cell count (Supporting Information Fig. S2), these findings are consistent with a developmental progression towards cell death for the ii1’ and ii2. The putative state change is supported by enrichment for GO terms such as *cellular catabolic process*, *xylan catabolic process*, *hemicellulose catabolic process*, *cell wall polysaccharide catabolic process*, *abscisic acid-activated signaling pathway* and *plant organ senescence* for ii1’ and ii2 clusters (Supporting Information Table S2). Indeed, ii1’ and ii2 had been described as lacking specialized features and undergoing programmed cell death of the seed coat layers the earliest. Nuclei of ii2 and ii1’ had also been shown to degrade at the heart stage (Haughn and Chaudhury 2005; Nakaune et al. 2005). Conversely, *MUCILAGE-MODIFIED 2* (*MUM2*), coding for a glycosyl hydrolase involved in mucilage formation, was enriched in the ii1’ (Western et al. 2000; Voiniciuc et al. 2015). Additionally, *CER1*, involved in cuticle formation, as well as *TT2*, *TTG1*, *PAP1*, *MYB114*, transcription factors involved in anthocyanin and proanthocyanidin synthesis, were detected in ii1’ clusters (Figure 2c,d) (Aarts et al. 1995; Nesi et al. 2001; Li 2014). These observations propel the question whether the ii1’ might metabolically support the surrounding layers, whether it provides a cuticle layer at some positions, or whether the ii1’ could modulate specialized metabolite production in neighboring cell layers.

Going from the outer cell layers towards the inner seed tissues, the cells of oi 2 are distinctly shaped by cutin and mucilage biosynthetic transcripts, while oi1 cells were highly enriched for flavonol synthesis. The inner integuments did not appear to be defined by specialized metabolites, but rather by a developmental progression, while the innermost integument layer, the endothelium, was again characterized by specialized metabolites, the proanthocyanidins. The endothelium was additionally characterized by a cutin layer, separating it from the products of double fertilization, the endosperm and embryo.

### Mirroring of embryo and endosperm distal/apical – proximal/basal patterns of seed coat integuments

Spatial transcriptomics of the AP2/ERF-family transcription factor *AINTEGUMENTA-LIKE 6* (*AIL6*), indicated that cluster 13 encompasses both endospermic and embryonal cells (Figure 1d). Assuming very different fates throughout development, we were surprised by the lack of distinction between endospermic and embryonal cells in the global UMAP. Subclustering analysis revealed transcriptionally distinct subpopulations, comprising 7 subclusters organized into two major groupings. At a subcluster resolution, spatial distribution of *MONOPTEROS* (*MP*) and *METHYLESTERASE 10* (*MES10*) indicated separation of embryonal (MP) and endospermic (MES10) domains, subcluster 2 being the embryo and subclusters 0, 1, 3-6, representing the endosperm (Figure 3a-b) (Cole et al. 2009).

**Figure 3:**
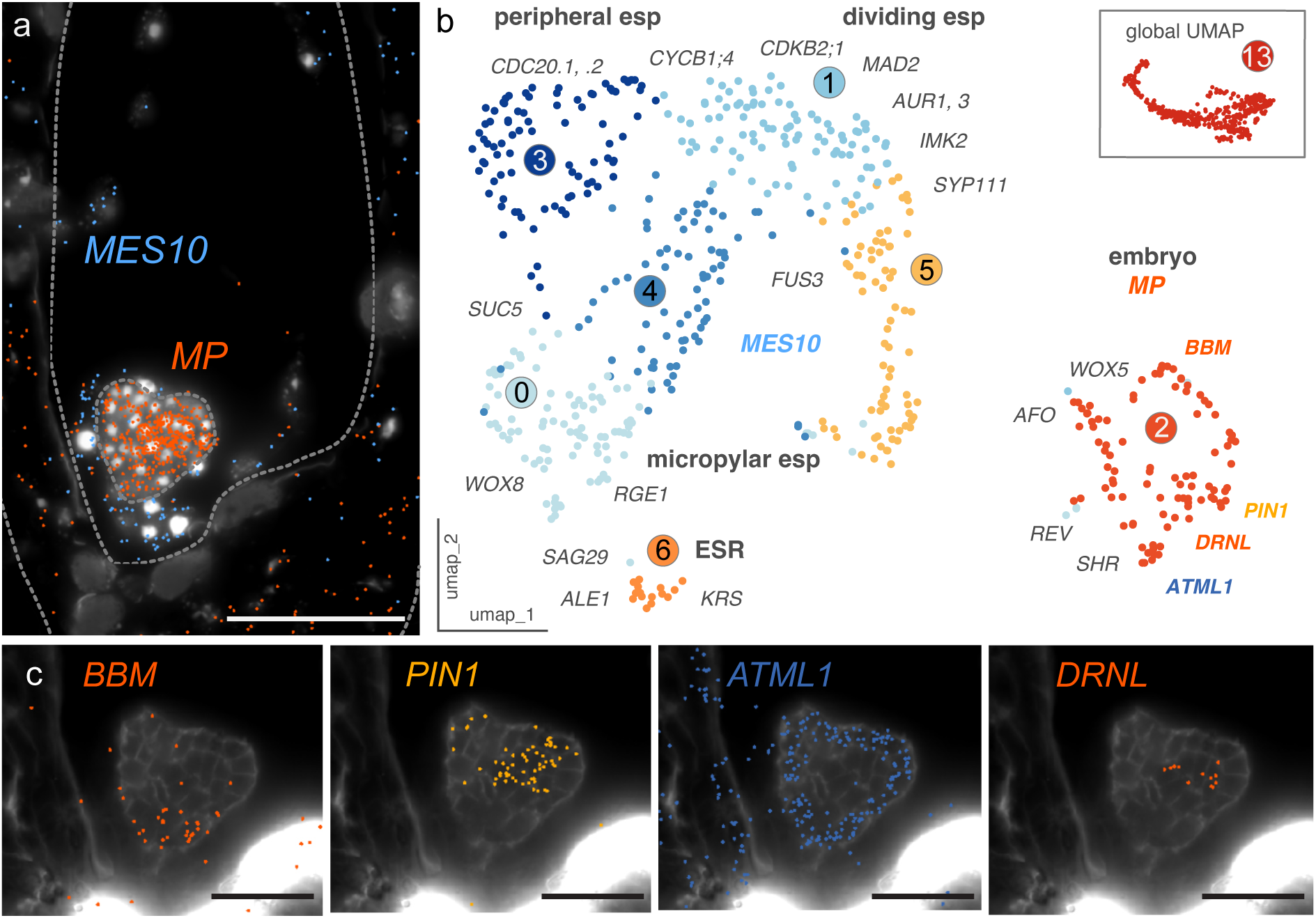
**Differentiation of endosperm and embryo along the distal/apical-proximal/basal axis.**(a) MERFISH spatial transcriptomics assigns embryo (*MP*, orange dots) and endosperm (*MES10*, blue dots) spatial domains. Nuclei are stained by DAPI, endosperm, embryo and seed outlines indicated by dashed lines. (b) Subclustering of the global UMAP cluster 13 results in seven subclusters representing spatial domains of the embryo and endosperm, as indicated by marker transcripts. esp, endosperm; ESR, embryo surrounding endosperm. The endosperm subclusters show a trajectory of transcriptional similarities from the basal/proximal embryo surrounding endosperm to the apical/distal peripheral endosperm. (c) MERFISH spatial transcriptomics maps (sub)epidermal, apical and basal embryo domains. Scale bars 50 µm (a), 25 µm (c).

The endosperm is a highly dynamic tissue. At the heart stage of Arabidopsis embryo development, the micropylar endosperm is cellularized, while the peripheral endosperm is actively undergoing cellularization. Consistent with active cellularization, subcluster 3 is marked by cell cycle–related genes such as *CYCB1;*3 and cytokinesis markers such as *SYP111*/*KNOLLE* (Lauber et al. 1997; Romeiro Motta et al. 2022). GO term enrichment for *microtubule organization* and *spindle assembly*, and enrichment of cytokinesis modulators including *MAD2* and *IMK2* further supports a role in endosperm cytokinesis (Caillaud et al. 2009; Smertenko et al. 2026). While subcluster 1 was enriched for cyclins (*CYCB1;4*), the enrichment of cell-cycle modulators (*CDC20.1*/*2*, *AUR1*, *AUR3*, *CDKB2*) in subcluster 3 reflected high proliferative activity (Supporting Information Fig. S7). Thus, subclusters 1 and 3 together map the proliferative and cellularizing states of peripheral endosperm, respectively (Menges et al. 2005; Van Damme et al. 2011; Yang et al. 2021).

The peripheral endosperm occupies the most distal region in the endosperm cavity, while the chalazal and micropylar endosperm are located in the proximal seed region. The chalazal endosperm is composed of the chalazal nodule and the chalazal cyst, which remains in an acellularized state throughout development, hence was not captured in the single-cell data, but was thoroughly assessed by Martin et al. 2026. The micropylar endosperm is characterized by an early onset of cellularization and its interaction with the embryo, e.g., in forming the embryonal cuticle from both the embryo and the embryo-surrounding endosperm (ESR). Multiple genes, such as *RETARDED GROWTH OF EMBRYO 1* (*RGE1/ZOU*), *ABNORMAL LEAF-SHAPE 1* (*ALE1*) and *KRS* are involved in the development of an embryonal-endosperm cuticle layer and known to be strictly confined to the endosperm, concentrated in the ESR (Tanaka et al. 2001; Kondou et al. 2008; Moussu et al. 2017). Their transcript enrichment patterns, together with expression of the micropylar endosperm markers sucrose transporter *SUC5*, and spatial transcriptomics data of *WOX8*, which is absent from the embryo but expressed in the surrounding endosperm, indicated a transcriptional trajectory from the peripheral endosperm identity (subclusters 3 and 1) to the micropylar endosperm (subclusters 4 and 0), culminating in an ESR identity (subcluster 6) (Haecker et al. 2004; Baud et al. 2005) (Figure 3b, Supporting Information Fig. S7-S10). The spatial enrichment of the amino acid transporter *UMAMIT25* further supported an ESR identity of cluster 6 (Supporting Information Fig. S8). Interestingly, the overall pathway activity score was enriched in the micropylar endosperm and ESR, while depleted in the peripheral endosperm (Supporting Information Fig. S9-10). Specially, hormone metabolism, amino acid metabolism and photosynthesis associated pathways showed consistently high PAS values in the ESR/micropylar endosperm subclusters 6 and 0 (Supporting Information Fig. S9-10). These features might indicate a high degree of functionalization in the developmentally older micropylar endosperm. Subcluster 6 was transcriptionally distinct from the peripheral endosperm, positioning the ESR at the interface between embryo and endosperm identities: ontogenetically related with the endosperm, but spatially and potentially functionally close to the embryo – presumably giving an indication as to why embryo and endosperm transcriptomes do not separate on the global UMAP. The embryo subcluster, consisting of 92 cells, displayed distinct embryonal domains. *WOX5* and *BABYBOOM* (*BBM*), expressed in the root meristem, were enriched in the upper half of subcluster 2 (Song et al. 2008; Horstman et al. 2015) (Figure 3 c). Conversely, the enrichment of, e.g., the auxin efflux transporter *PIN1* indicated the representation of the apical embryonal domain in the lower half of subcluster 2. Epidermal cells, marked by *ATML1*, as well as subepidermal cells of the apical portion of the embryo, marked by the transcription factor *DRNL*, were represented in subcluster 2 as well (Aida et al. 2002; Chandler et al. 2007) (Figure 3 c). The distinct subpopulations of both the endosperm as well as the embryo mirror the apical/distal – basal/proximal axis of the seed coat.

### Micropylar and chalazal domains of the proximal seed

In contrast to the distal seed coat, the proximal seed coat does not appear to be composed of distinctly layered cells (Figure 4a). To identify canonical cell types in the morphologically heterogeneous proximal seed coat, we analyzed transcript enrichment patterns of published marker genes and our spatial transcriptomics dataset and identified eight clusters belonging to the proximal seed coat. These included unique cell types belonging to the nucellus (cluster 18), ULD (cluster 19), abscission zone (cluster 17), as well as multiple clusters of integument origin, such as two endothelial (clusters 14, 15) and two oi1 (clusters 6, 8) clusters (Figure 4a-b). The presence of two distinct endothelium and two distinct oi1 clusters prompted us to ask whether there would be additional factors separating these two populations of the same origin. We therefore mapped the clusters using known markers with distinct spatial domains, including *RR22*, a cytokinin-related chalazal seedcoat marker, *PIN3*, an auxin transporter that localizes predominantly in the chalaza, the *SWEET12*, *UMAMIT15* transporters and *TT10*, as well as *TT1*, expressed in the general and micropylar endothelium, and *ATT1/CYP86A2,* enriched in the general and micropylar endothelium, but reduced in the chalaza (Pourcel et al. 2005; Horák et al. 2008; Chen et al. 2015; Coen et al. 2019, 2020; Liu et al. 2023) (Figure 4c). For both endothelial and oi1 proximal seed coat clusters, a micropylar and a chalazal cluster were identified. Notably, this points towards a micropylar-chalazal spatial axis, adding to the complexity of the proximal seed coat. To better understand the basis of the transcriptional distinction between the micropylar and chalazal cell populations of the same ontogeny, we compared GO term enrichment between chalazal (clusters 15, 8) and micropylar (clusters 14, 6) subgroups. Subgroupings of the same cell type (cluster 14, 15 and cluster 6, 8) shared fewer GO terms than the different cell types from the same spatial domain; the only shared GO terms between the endothelial clusters 14 and 15 were those related to *flavonoids*, consistent with the defining role of specialized metabolites in the endothelium (Figure 4d, Supporting Information Table S2). The micropylar subgroup was enriched for GO terms related to *fatty acid metabolism*, *glutathione metabolism* and *cell wall metabolism*, in agreement with the enriched pathway activities for very long fatty acid chains, wax, xylan and cutin-related pathways (Figure 4d, Supporting Information Fig. S4). The chalazal subgrouping was defined by GO terms associated with *di/oligosaccharide, trehalose, nucleobase metabolism*, and *fluid transport* (Figure 4d). The endothelial cells engulfing the nucellus are referred to as the pigment strand. In our data, the pigment strand cells were enriched for pathways related to hormone signaling and amino acid metabolism (Supporting Information Table S2). Surprisingly, only five GO terms were shared between the two oi1 populations. The top five GO terms for the micropylar oi1 were most significantly enriched for *cell wall polysaccharide metabolic process*, whereas the chalazal oi1 terms were related to *di/oligosaccharide metabolic process*, *water transport*, *trehalose metabolic process*, *nucleobase metabolic process* (Supporting Information Table S2). The distinct transcriptional separation of cell populations of the same ontogeny based on their spatial domain raises the question whether, in the seed coat, positional cues and the spatial context exert a greater influence on transcriptional identity than the developmental lineage.

**Figure 4:**
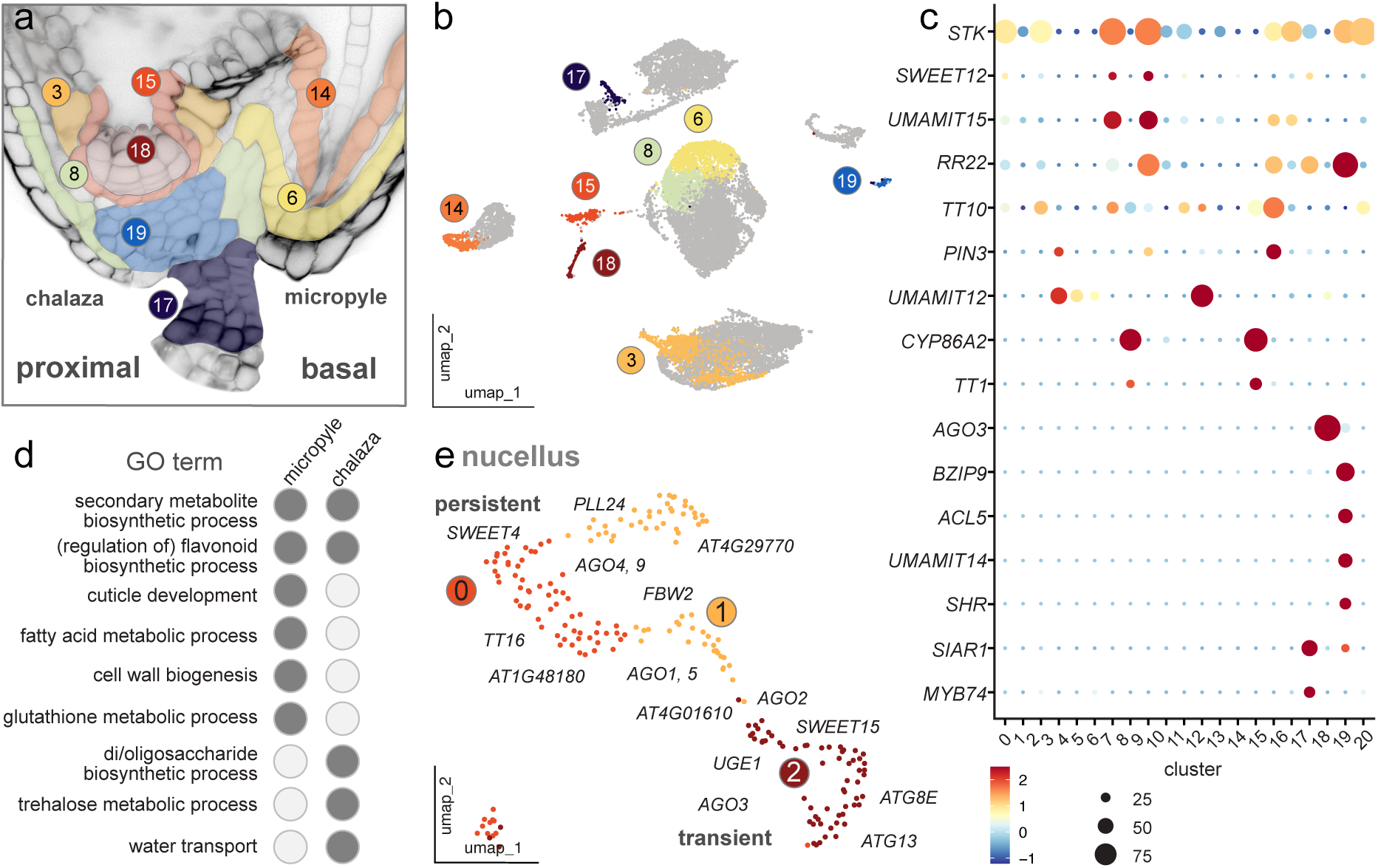
Cell-types within the proximal seed coat revealed by spatially resolved subdomains of micropyle and chalaza. (a) Close-up of a representative Arabidopsis proximal/basal seed coat with cell layers false-colored corresponding to scRNA-seq cluster annotation, marked on a cell layer level in (b) UMAP plot of integrated scRNA-seq datasets. (c) Transcripts identifying different proximal seed coat populations. Dot size indicates percent of cell in which transcript was detected, color indicates log fold enrichment, legend on the bottom. (d) The two proximal seed coat spatial domains, micropyle and chalaza, are distinguished by unique GO terms. Representative GO terms were selected from the three highest adjusted p-value GO terms in a comparative GO term enrichment analysis (Supporting Information Table S2, code in Supporting Information Note 2). (e) Subclustering the nucellus cluster results in three subclusters. Marker transcripts map the persistent and transient nucellus cell populations along a clear proximal-distal gradient.

Adding to the spatial context of apical-basal/distal-proximal, the basal seed coat is further patterned along the micropylar-chalazal axis. Positionally, several additional clusters were assigned as unique to the chalaza, including the abscission zone, the unloading domain and the nucellus. The Arabidopsis nucellus is a transient tissue which is eliminated from 2 DAP in a distal-to-proximal progression, thereby creating space for endosperm growth. However, a few nucellar layers persist to form the persistent nucellus, which has been implicated in nutrient transport from the chalaza to the endosperm (Lu and Magnani 2018). The nucellus marker *AGO3* as well as other *AGO* family members, including *AGO1* and their upstream regulators (*FBW2*) and targets (*tasiRNA*) were enriched in cluster 18 (Jullien et al. 2020) (Figure 4e, Supporting Information Fig. S11). Interestingly, the nucellus cluster seemed to display a clear trajectory. Subclustering of the nucellus population resulted in three subclusters (Figure 4e). Subcluster 0, annotated as the persistent nucellus, was enriched with *SWEET4*, *TT16/ABS*, *PLL24*, in accordance with its roles in nutrient transport (Lu et al. 2021; Banfi et al. 2025; Iannaccone et al. 2026) (Figure 4e, Supporting Information Fig. S11). GO terms related to *amino acid metabolism*, *glucan metabolism* and *polysaccharides* were enriched, but also *programmed cell death* (Supporting Information Table S2). Subcluster 2, was enriched with positive regulators of cell death (*AT4G01610,* a putative cysteine protease), as well as autophagy related transcripts, such as *ATG8E* and *ATG13*, and *UGE1*, an UDP-glucose and xylose-4-epimerase involved in cell wall remodeling (Barber et al. 2006), representing cell states of the transient nucellus (Ge et al. 2016) (Figure 4e, Supporting Information Fig. S11). GO term enrichment of subcluster 2 highlighted terms related to *nucleoside catabolism*, *autophagy*, *mitophagy*, *vacuole organization*, supporting the transient identity (Supporting Information Table S2). Interestingly, *SWEET15* was identified in 12 cells representing the transient nucellus, and transcriptional enrichment seemed to translate into protein level, underpinning the recently explored dynamics between cell death and nutrient transport within the nucellus (Supporting Information Fig. S11).

Nutrient transport is one of the defining features of the unloading domain, where the maternal vasculature terminates and nutrients are transferred to the developing seed (Liu et al. 2025). The unloading domain is characterized by various transcripts associated with vascular cell types, including xylem and phloem markers, such as *ACL5* and *bZIP9* (Muñiz et al. 2008; Kim et al. 2021) (Figure 4c, 5a-b, Supporting Information Fig. S1). Multiplexed *in situ* hybridization of *bZIP9*, *UMAMIT14* and *CALS8/GSL04* spatially mapped the unloading domain identity of cluster 19 (Figure 5a). Notably, numerous transport-associated transcripts are highly enriched in cluster 19, including members of the UMAMIT family (*UMAMIT10*, *23*, *14*), SWEET sugar transporters and *GTR*s, as well as callose synthases, such as *CALS8* (Figure 5c). While transport associated transcripts are enriched in the ULD, pathway activities related to several phytohormones were also enriched, supporting the notion that the ULD functions as the gate connecting the seed to the rest of the plant through the funiculus (Liu et al. 2025; Xu et al. 2025) (Supporting Information Fig. S4, S12).

**Figure 5:**
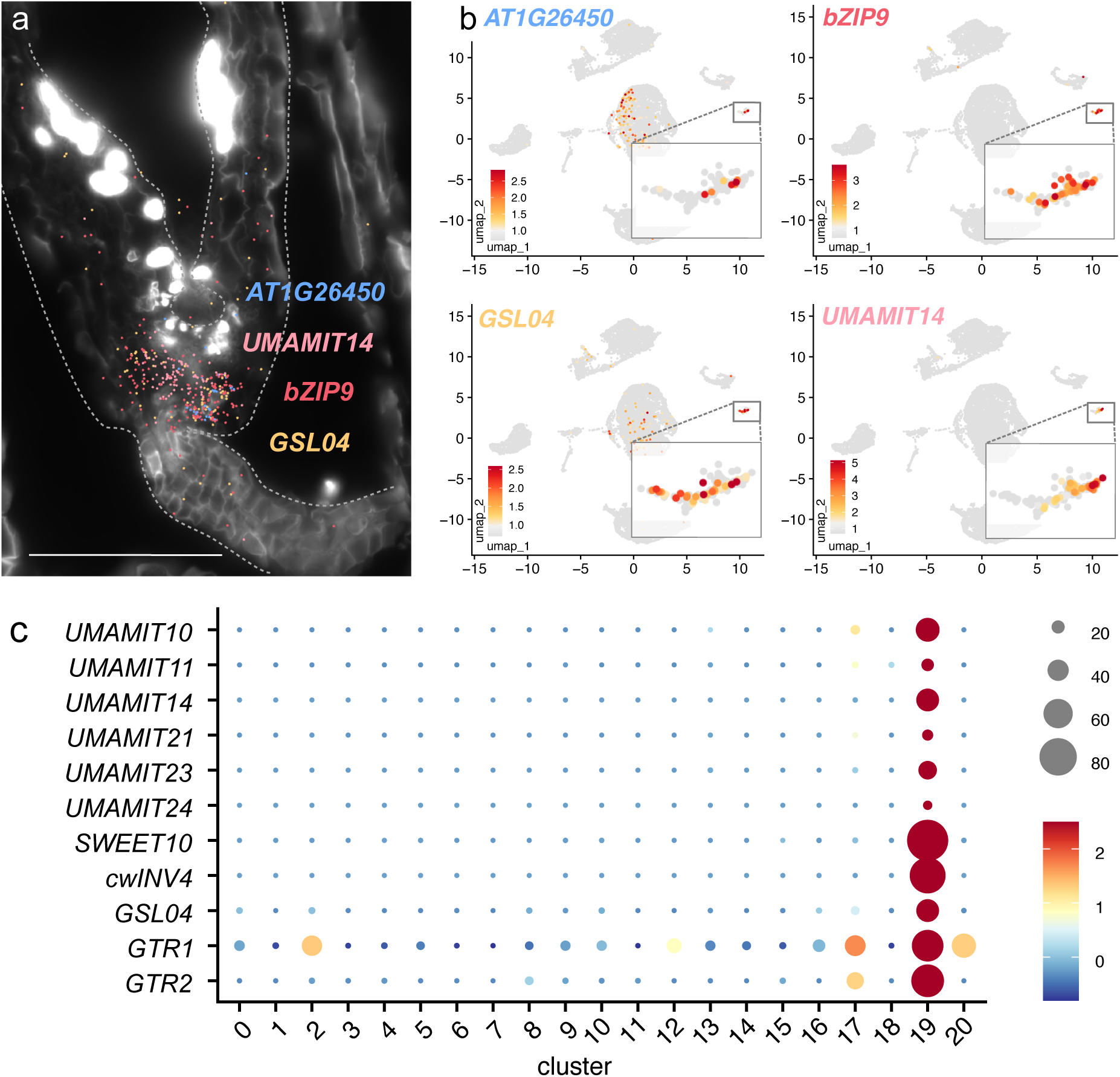
Transport associated transcript enrichment in the seed unloading domain. (a) Multiplexed *in situ* hybridization image showing spatial localization of *AT1G26450* (blue), *UMAMIT14* (pink), *bZIP9* (magenta), and *GSL04/CALS8* (yellow) transcripts within the seed ULD. (b) UMAP plots showing enrichment of *AT1G26450*, *bZIP9*, *GSL04/CALS8* and *UMAMIT14* in cluster 19. Insets show magnified view. (c) Dot plot of transport-associated transcript enrichments in cluster 19 across all clusters (clusters 0–20). The diameter of the dot indicates the percentage of cells within a class, while the color encodes average enrichment across all cells within a class. Scale bar 100 µm.

### Characterization of the most proximal seed tissue - the seed abscission zone

The funiculus connects ovules to the maternal plant. After fertilization and until the maturation phase of seed development, seeds remain connected to the funiculus. Upon maturation, seed plants shed/abscise their seeds to facilitate seed dispersal. While seed abscission itself is not much studied, abscission has been characterized extensively in the context of floral organ shedding. The floral abscission zone (AZ) is composed of two distinct cell layers, between which separation occurs: residuum and secession cells. Secession cells form a lignin brace enabling shedding, while residuum cells remain on the organ and differentiate into epidermal-like cutin containing cells, reclosing the open surface (Lee et al. 2018). Spatial transcriptomics of several marker genes supported a seed AZ identity of cluster 17 (Figure 6a).

**Figure 6:**
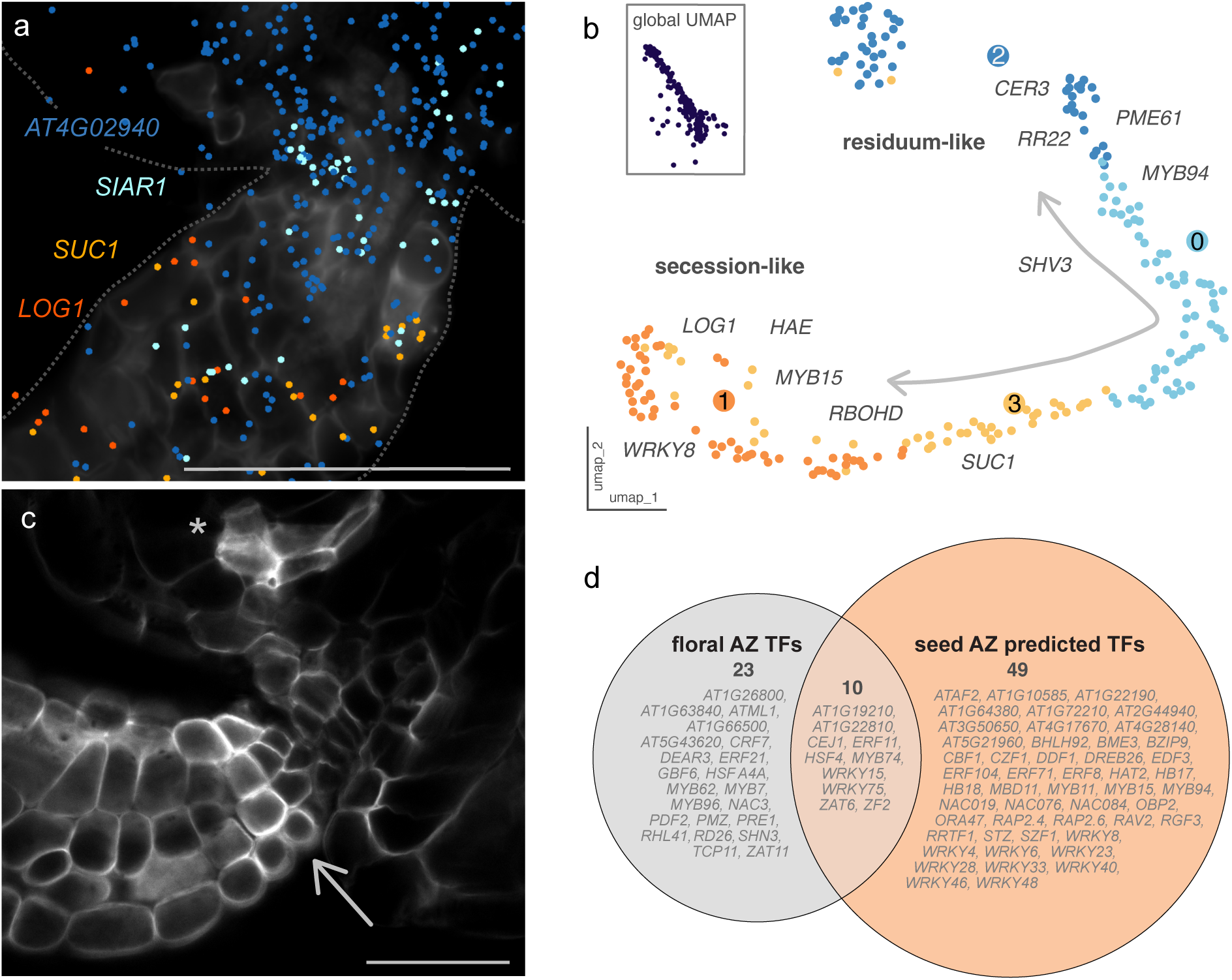
Mapping the seed abscission zone as the most proximal seed tissue. (a) MERFISH spatial transcriptomics assigns the spatial subdomains corresponding to (b). Subclustering of the global UMAP seed abscission zone cluster 17, resulting in four subclusters that show a trajectory overlapping with cellular identities of secession and residuum-like cells of floral abscission. (c) Confocal laser scanning microscopy reveals distinct cellular features of the seed AZ, including an intense cell wall staining, marked by an arrow (similar to intense sieve element cell wall stain, marked by an asterisk). (d) Prediction of putative and known transcription factors in the seed abscission zone compared with transcription factors in the floral abscission zone from Wen et al. 2025 highlights shared and seed abscission zone specific transcription factors. Scale bars 50 µm (a), 25 µm (c).

To explore whether distinct cell types of the floral AZ can be found in the seed AZ cluster, we examined canonical markers of secession and residuum cells from Lee et al. 2018. Subclustering of the putative seed AZ revealed 4 subclusters, with two being spatially distinct, connected through two additional clusters on the UMAP (Figure 6b). We mapped and summarized transcript enrichment of the top 500 floral secession and residuum cell DEGs from Lee et al. 2018 by p-value onto the subclusters. 34.2% of floral secession and 15% of floral residuum cell canonical markers were identified in cluster 17 (Supporting Information Note 2, Supporting Information Fig. S13). The enrichment of floral AZ secession cell markers in cluster 1 and residuum cell markers in cluster 2 indicated that the seed AZ showed transcriptional profiles of both major floral AZ cell types, secession and residuum. The enrichment of the chalazal seedcoat marker *RR22* in cluster 2 and 0 spatially mapped the “residuum-like” cells on the seed side of the AZ, supported by spatial transcriptomics markers (Figure 6a-b, Supporting Information Fig. S14). Between these transcriptomic signatures, we observed a trajectory, as may be expected ongoing differentiation. As the full identity of residuum cells manifests only after abscission, we were surprised to identify secession and residuum cell profiles already at the heart stage of embryo development, days prior to abscission. Nonetheless, the β-1,4/1,3-glucan specific dye SR2200 strongly stained the AZ already at this stage (Musielak et al. 2015) (Figure 6c). Together with the enrichment of transcripts such as the cellulose synthase-like *CSLD3* (Figure 6a, Supporting Information Fig. S14), these findings provide a basis to dissect the cellular differentiation of the seed AZ.

To evaluate a potential developmental progression of AZ cells at the heart stage of embryo development, we examined AZ differentiation state markers of floral residuum cells, as the seed AZ clusters bore similarities to floral AZ cell identities (Wen et al. 2025). We found that 27% of AZ differentiation state markers of floral residuum cells were detected in cluster 17 (Supporting Information Fig. S13). While for state 1, which has been characterized by GO terms associated with *photosynthesis* and *translation* in floral AZ, only 13% of marker transcripts were detected in the seed AZ, while for floral AZ state 3, which has been associated with *lipid, cell wall, wax*, already 33% of marker transcripts were detected in the seed AZ cluster. Floral AZ state 2, which has been associated with GO terms associated with *response to stress*, 47% of marker transcripts were detected, suggesting that the seed AZ at heart stage of embryo development shows most similarity to the state 2 of floral AZ (Supporting Information Fig. S13). Consistent with this, GO terms described for the floral AZ state 2 have been detected in our GO terms analysis of the seed abscission zone (Wen et al. 2025) (Supporting Information Table S2).

State 2 has been described as a transition state where the mesophyll identity of the residuum cells transdifferentiates towards an epidermal identity. Since MYB TFs have been implicated in differentiation processes in general, we searched in our predicted TFs for MYB family members. Indeed, we identified *MYB15, MYB94* and *MYB74* as markers of the seed AZ (Supporting Information Fig. S14). Interestingly, MYB74 had been implicated in the trans-differentiation of residuum cells in floral AZ and its expression peaked before abscission, indicating further transcriptional similarities between floral and seed AZ (Wen et al. 2025). By comparing the TFs listed in (Wen et al. 2025) with marker genes of cluster 17 classified/predicted as TFs, we found an overlap of 10 of 33 TFs in the seed AZ cluster, including known AZ modulators such as *MYB74* (Figure 6d). Two of the predicted TFs of the MYB family, *MYB15* and *MYB94*, especially attracted our attention since it had recently been shown that MYB15 plays a role in lignin production, and was enriched in the secession-like cells in our data, while MYB94 was previously shown to be involved in cuticular wax formation while being enriched in the residuum-like cells in our data (Lee et al. 2016; Kim et al. 2020). We then asked whether other TFs specific to the seed AZ, and distinct from the floral AZ regulatory program, could be identified. Notably, we identified 49 putative seed AZ cluster specific TFs (Figure 6d). Among these, 18% belonged to the WRKY family which coordinate transcriptional reprogramming in response to diverse developmental and environmental cues (Bakshi and Oelmüller 2014). In the floral AZ, genes associated with responses to biotic and abiotic cues shown to be enriched in the residuum cells prior to abscission (Wen et al. 2025). This has led to the hypothesis that the absence of a protective cuticle layer during abscission exposes the residual tissue to a/biotic risks, thus prompting residuum cells to transition to an epidermal identity before abscission to prepare the plant for potential (a)biotic cues (Wen et al. 2025). Previous work demonstrated that several WRKY-binding sites are required for the transcriptional competence of the secreted peptide IDA, a major trigger of floral AZ formation (Galindo-Trigo et al. 2024). We identified enrichment of transcripts known to be directly involved in cell differentiation and abscission, such as *HAE, IDL7* and *MLO2*, in cluster 17 as putative targets of WRKYs (Butenko et al. 2003; Vie et al. 2015) (Supporting Information Fig. S14). The comparative analysis of floral and seed AZ single-cell transcriptomes revealed three major aspects: (1) the transcriptional program described for floral residuum cells showed parallels in the seed AZ, and the seed AZ transcriptomes might also capture elements of secession cell differentiation, (2) the enrichment of floral AZ state 2 markers indicated that already at the heart stage of embryo development, the seed AZ is transcriptionally preparing for abscission and that (3) WRKY TFs might play an important role in facilitating seed abscission. Together, these findings provide a transcriptional framework for investigating the seed AZ and future studies could profit from probing the seed AZ specific WRKY and MYB TF mutants.

## Discussion

Here, we established a protoplast isolation strategy that, combined with single cell transcriptomics, bioinformatics, and spatial transcriptomics enabled the capture of a comprehensive cellular transcriptional profiles of seed cells. Our analyses reveal that the developing seed is more transcriptionally heterogeneous than previously appreciated and identified trajectories and different cell states. While canonical seed tissues had been anatomically well defined, we here identified extensive transcriptional diversification within the seed coat, including developmentally distinct epidermal states, the embryo and endosperm and spatially distinct micropylar and chalazal domains. We provide a first cellularly resolved transcriptional characterization of the seed abscission zone and explore nutrient transport and cell death dynamics in the persistent and transient nucellus. Our findings reveal a high degree of differentiation of cell layers along the rotational and axial seed axes, highlighting the heterogeneity in importance of cell position and ontogenesis on forming cell identities. In the seed, cell identity is, next to spatial position and ontogenesis, additionally shaped by a cascade of developmentally programmed cell death. The developmentally programmed cell death starts as early as fertilization with the elimination of ovule tissues and ultimately results in a 2-cell layered seed coat and one cell layer of endosperm left surrounding the mature embryo (Haughn and Chaudhury 2005). We provide evidence for programmed cell death in multiple cell types at the heart stage of embryo development. Because cellular structures persist for a period after nuclear degradation, single-cell transcriptomics provides unique access to transcriptional states that are inaccessible to single-nucleus approaches, enabling the characterization of cells undergoing the transition toward elimination. Profiling seed single-cell transcriptomes across additional developmental stages and integrating scRNA-seq with high resolution spatial transcriptomics through imputation will further enhance our understanding of the dynamic cellular programs during seed development (Demesa-Arevalo et al. 2026).

## Supporting information

Supplementary Table and Script

## Acknowledgments

We sincerely apologize to the authors whose work could not be cited due to space limitations. We thank Tobias Lautwein from the sequencing facility of the Genomics & Transcriptomics Laboratory (HHU) for running the BD Rhapsody system and sequencing services, Rüdiger Simon and Svenja Augustin for advice on spatial transcriptomics. We also thank Diana Weidauer for assistance in seed harvesting for protoplast isolation, and Inkyung Cho for her help in assessing the seed phenotype. We thank Rosanna Petrella for critical reading of the manuscript. This work was supported by grants from the Alexander von Humboldt Professorship (WBF), Deutsche Forschungsgemeinschaft (DFG, German Research Foundation) under Germanýs Excellence Strategy – EXC-2048/1 – project ID 390686111 (CEPLAS). Work of JYK’s lab was supported by Basic Science Research Program through the National Research Foundation of Korea (NRF) funded by the Ministry of Education (No. RS-2019-NR040081) and by the National Research Foundation of Korea (NRF) grant funded by the Korea government (MSIT), No. RS-2025-00560593 and No. RS-2026-25511199.

## Conflict of interest

The authors declare no conflict of interest. Access to transgenic lines is available under MTA from HHU.

## Author contributions

KLBR, JYK and WBF developed the concept. KLBR performed the experiments, NRZ and MMW contributed to spatial transcriptomics. KLBR, WBF and JYK analyzed the data. TH advised bioinformatics, TYP performed pathway analysis, advised by MJL. KLBR, WBF and JYK wrote the manuscript. All authors have approved the final version of the manuscript.

## Abbreviations

Single cell RNA-sequencing (scRNA-seq)

## Notes

### Competing Interest Statement

The authors have declared no competing interest.

