## Supplementary material for "Cell identities along the proximal-distal and micropylar-chalazal axes in the Arabidopsis heart-stage seed": Suppl.

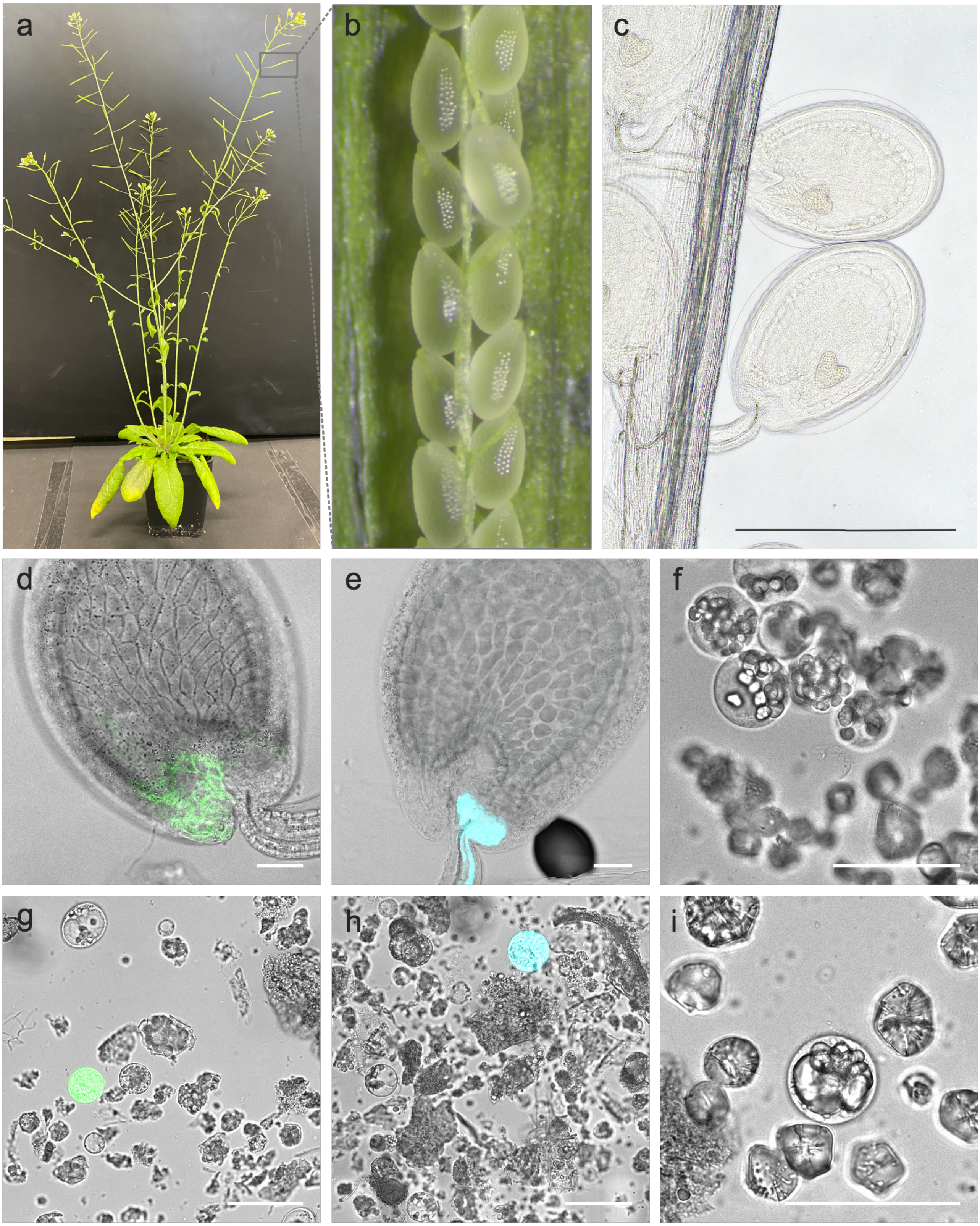


**Supporting Information Figure S1: Selection of seeds for scRNA-seq experiments and representation of diverse cell types in protoplast populations for scRNA-seq.** (a) Representative image of one of the five week-old *Arabidopsis thaliana* plants used for seed isolation for scRNA-seq experiments. Box indicates an example of a sampled silique. (b) After selection, siliques were dissected, exposing the developing seeds. Seeds were visually assessed by their size, shape and micropylar greening. Individual seeds were dissected to probe the developmental stage during seed selection and embryo developmental stage was again assessed in the protoplasting suspension prior to washing steps. (c) Exemplary image of siliques selected in (b) subjected to Hoyers' clearing to confirm the developmental stage. To develop a methodology for seed protoplasting yielding protoplasts from diverse tissues, fluorescent marker reporter lines (d-e, g-h) and morphological features such as starch granules and specialized vacuoles were employed (f, i). (d) Confocal microscopy image of a chemically fixed and cleared *Arabidopsis thaliana* seed expressing *pSWEET12:SWEET12-eGFP* showed fluorescence (green) predominantly in the micropylar integuments. The seed is shown using a transmitted light detector.

(e) Confocal microscopy image of a chemically fixed and cleared *Arabidopsis thaliana* seed expressing *pbZIP9:GFP-GUS* showed fluorescence (cyan) in the phloem of the funiculus and in the seed unloading domain. The seed is shown using a transmitted light detector. Confocal and transmitted light microscopy images of protoplasts (prior to washing steps) containing starch granules (f), SWEET12-GFP positive protoplasts (g), ULD protoplasts marked by *pbZIP9:GFP* (h) and protoplasts with specialized vacuoles (i) indicated a representation of diverse cell types in protoplast populations generated from our seed-optimized protocol. Crystal-like structures in (f, i) likely represent unfiltered enzyme crystals. Scale bars (c) 500 µm, (d-i) 50 µm.

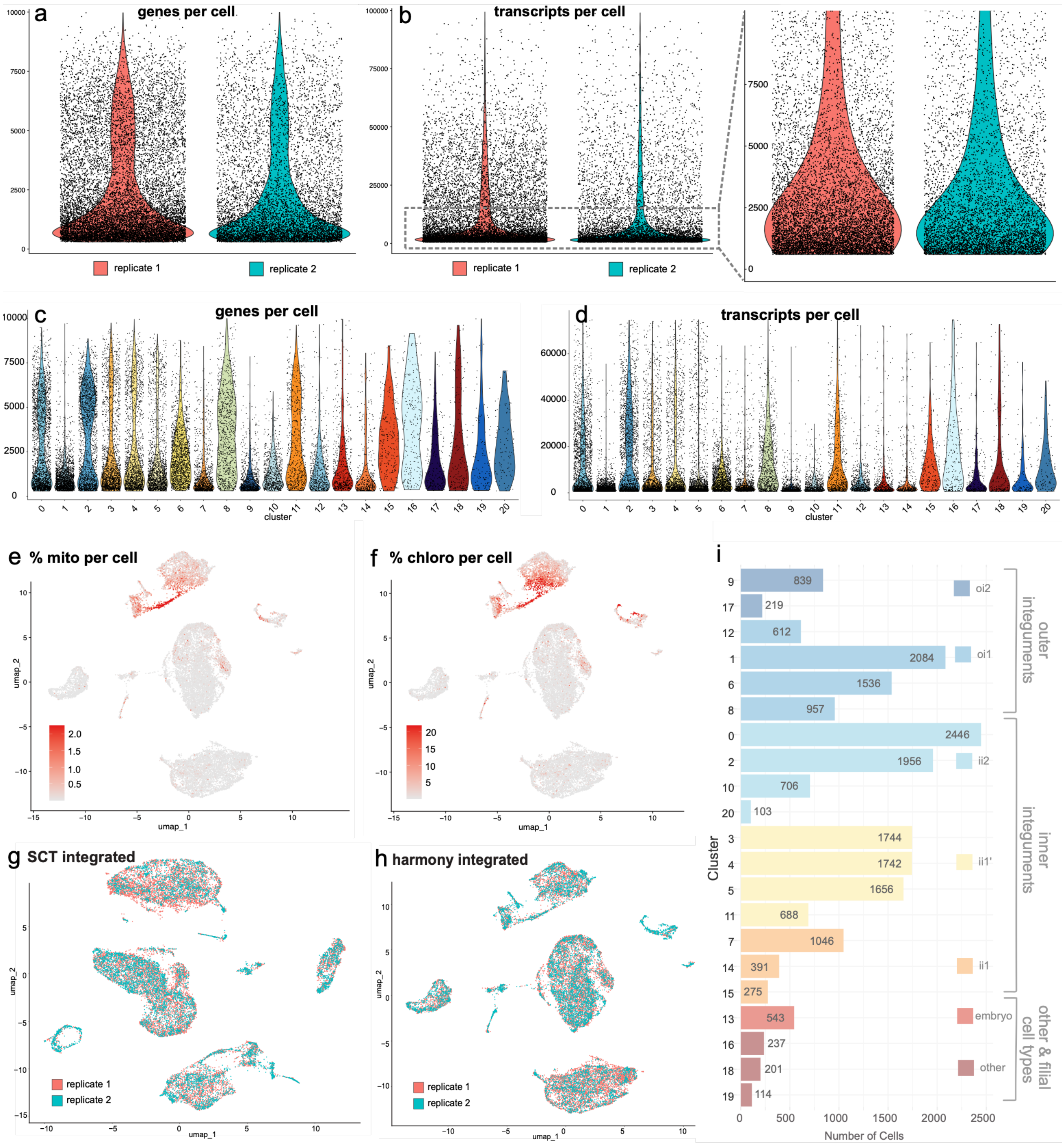


**Supporting Information Figure S2: Quality metrics of two independent scRNA-seq datasets.** (a,b) Violin plots showing the number of genes (a) and transcripts (b) detected per cell for replicate 1 (red) and replicate 2 (teal). Inset shows a magnified view of the transcript distribution for both replicates. (c,d) Violin plots showing the number of genes (c) and transcripts (d) detected per cell, grouped by clusters. Y-axis of (d) was limited to 75,000 transcripts. (e,f) UMAP projections colored by the percentage of mitochondria (e) and chloroplast (f) transcripts per cell. (g,h) UMAP projections comparing dataset integration methods, SCT transform (g), and harmony (h) with replicate 1 colored red and replicate 2 colored teal. (g) Bar chart of cell numbers per cluster (cluster 0-20), grouped and color-coded by tissue identity.

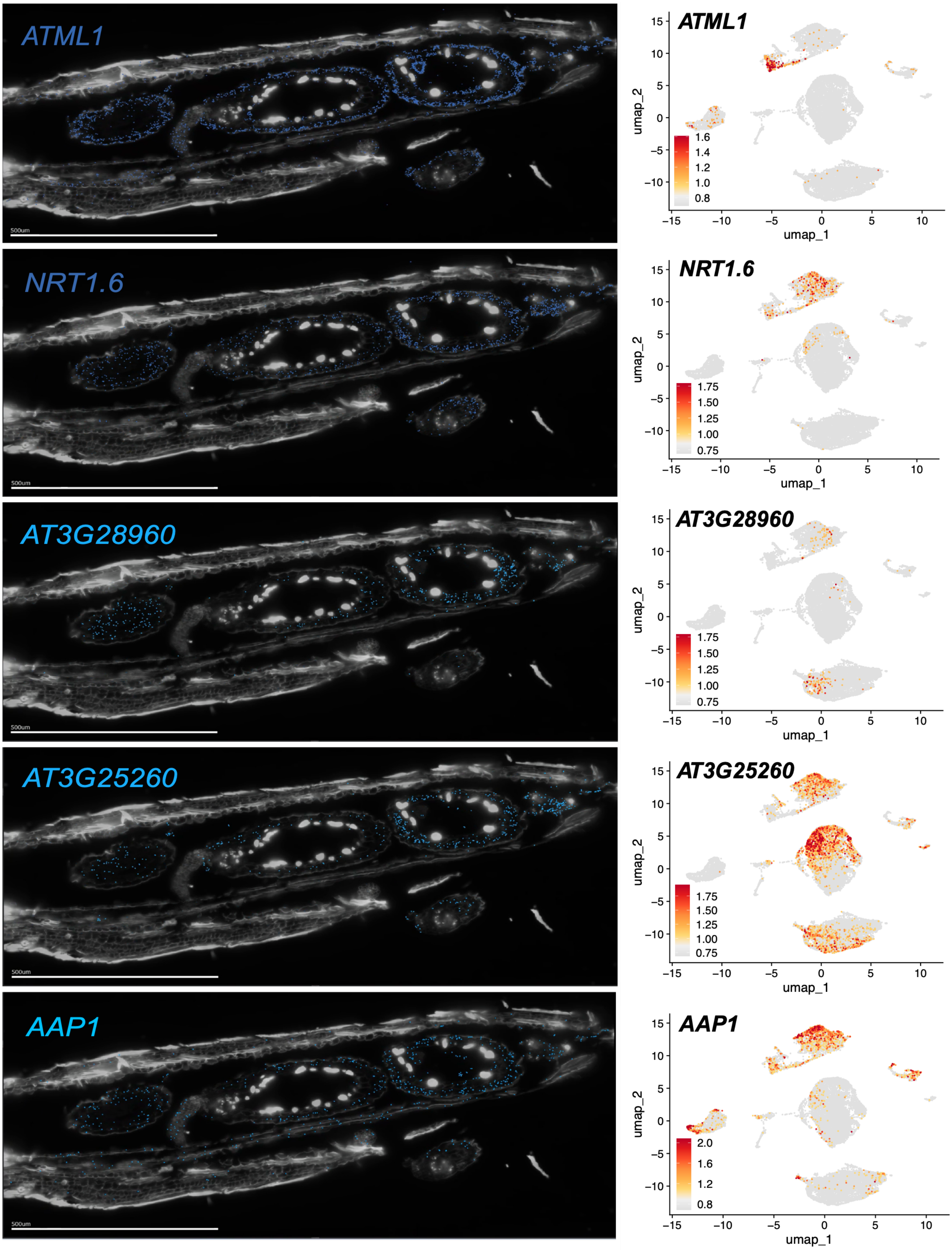

ATML1
500um
NRT1.6
500um
AT3G28960
500um
AT3G25260
500um
AAP1
500um
ATML1
NRT1.6
AT3G28960
AT3G25260
AAP1
umap_1
umap_2

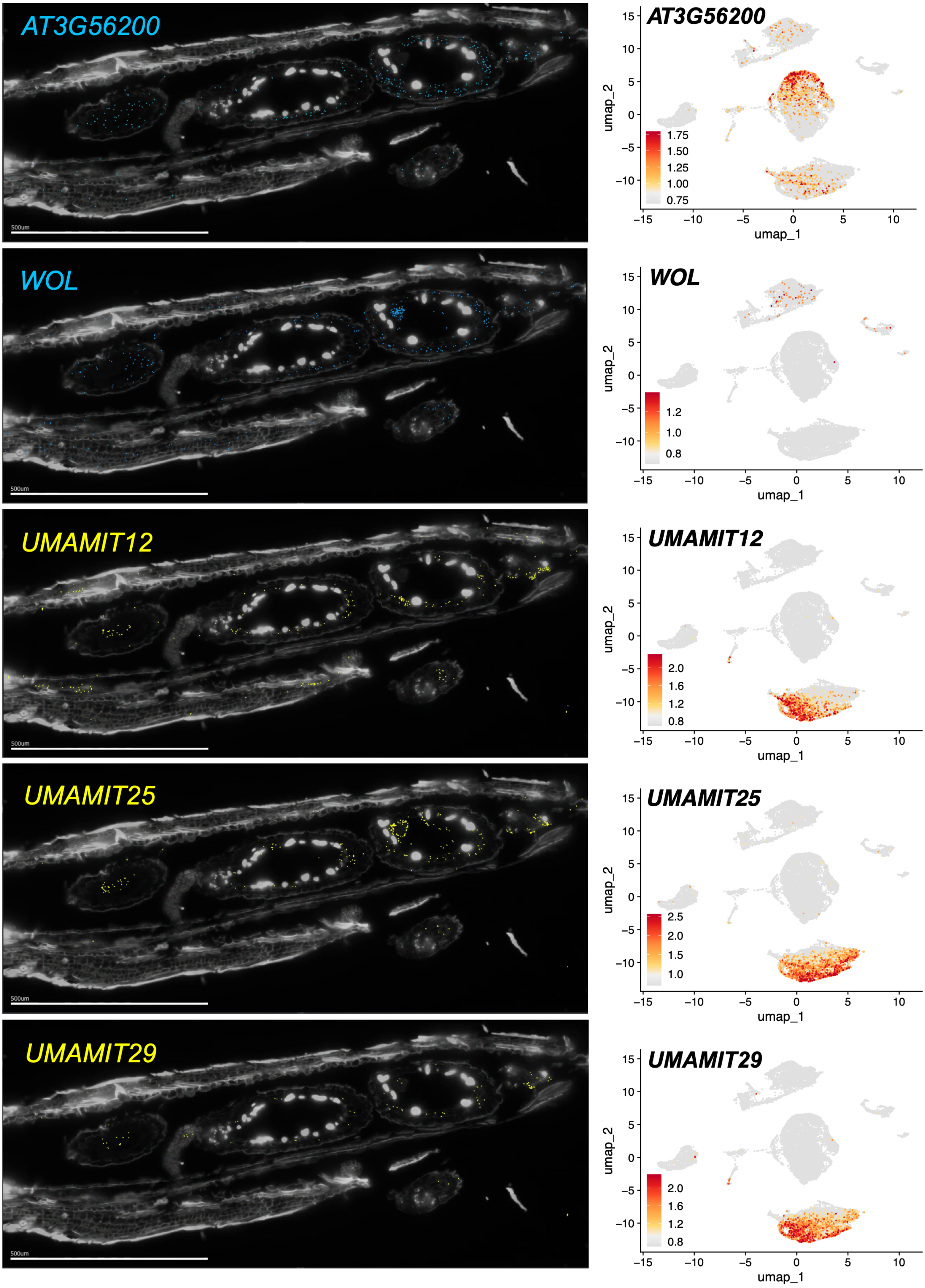

AT3G56200
WOL
UMAMIT12
UMAMIT25
UMAMIT29
500um
umap_1
umap_2
1.75
1.50
1.25
1.00
0.75
1.2
1.0
0.8
2.0
1.6
1.2
0.8
2.5
2.0
1.5
1.0
−15
−10
−5
0
5
10
15

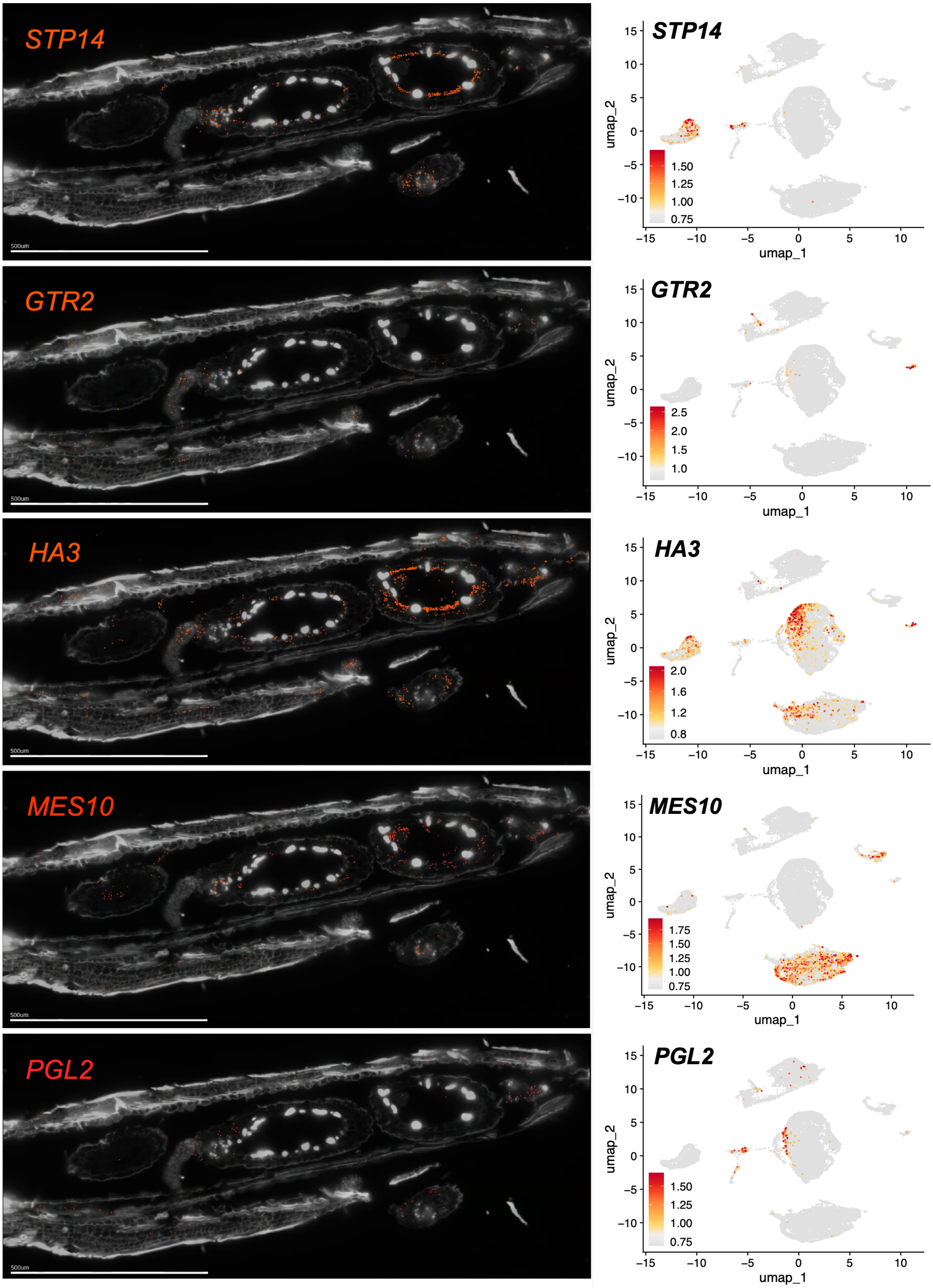

STP14
500um
STP14
umap_2
umap_1
1.50
1.25
1.00
0.75
GTR2
500um
GTR2
2.5
2.0
1.5
1.0
HA3
500um
HA3
2.0
1.6
1.2
0.8
MES10
500um
MES10
1.75
1.50
1.25
1.00
0.75
PGL2
500um
PGL2
1.50
1.25
1.00
0.75

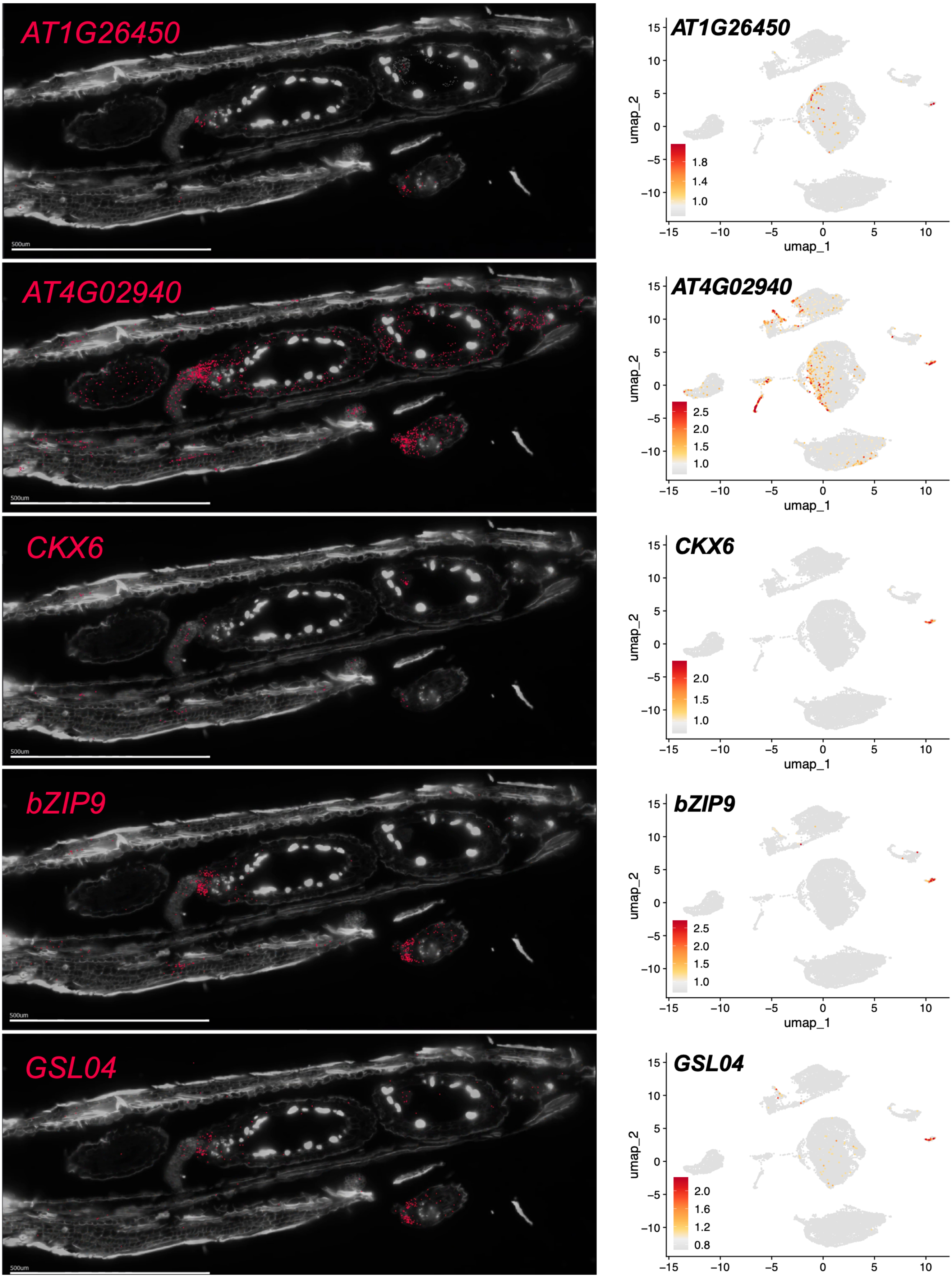
AT1G26450
500um
AT1G26450
umap_2
umap_1
1.8
1.4
1.0
AT4G02940
500um
AT4G02940
umap_2
umap_1
2.5
2.0
1.5
1.0
CKX6
500um
CKX6
umap_2
umap_1
2.0
1.5
1.0
bZIP9
500um
bZIP9
umap_2
umap_1
2.5
2.0
1.5
1.0
GSL04
500um
GSL04
umap_2
2.0
1.6
1.2
0.8

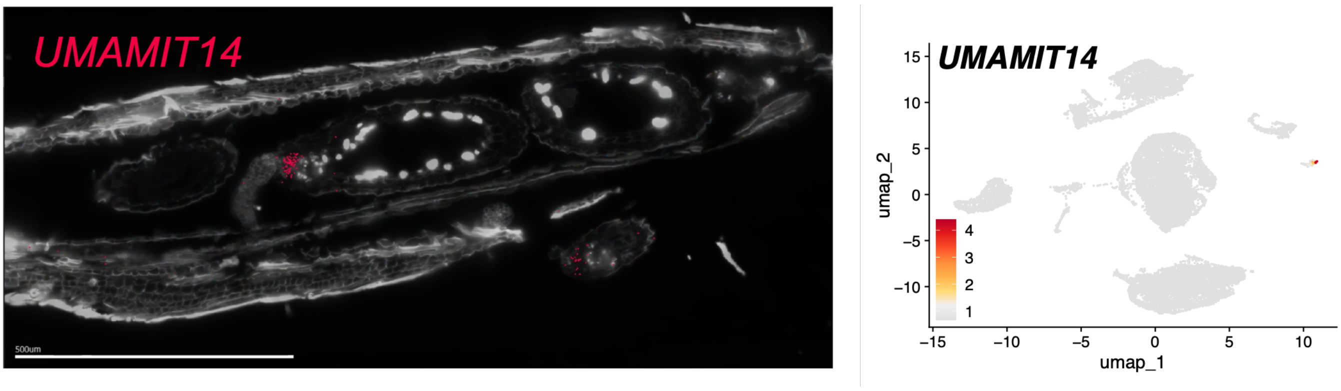


**Supporting Information Figure S3: MERFISH spatial transcriptomics data complements scRNA-seq cluster annotation.** Multiplexed *in situ* hybridization images of a silique enclosing three developing seeds sectioned at different seed planes showing spatial localization of marker transcripts in the adaxial epidermis (dark blue), abaxial epidermis (light blue), inner integument 1' (yellow), endothelium, chalaza/pigment strand (orange), chalaza/unloading domain (pink/red) (left) and corresponding transcript enrichment in scRNA-seq global UMAP plots (right). Scale bars 500 µm.

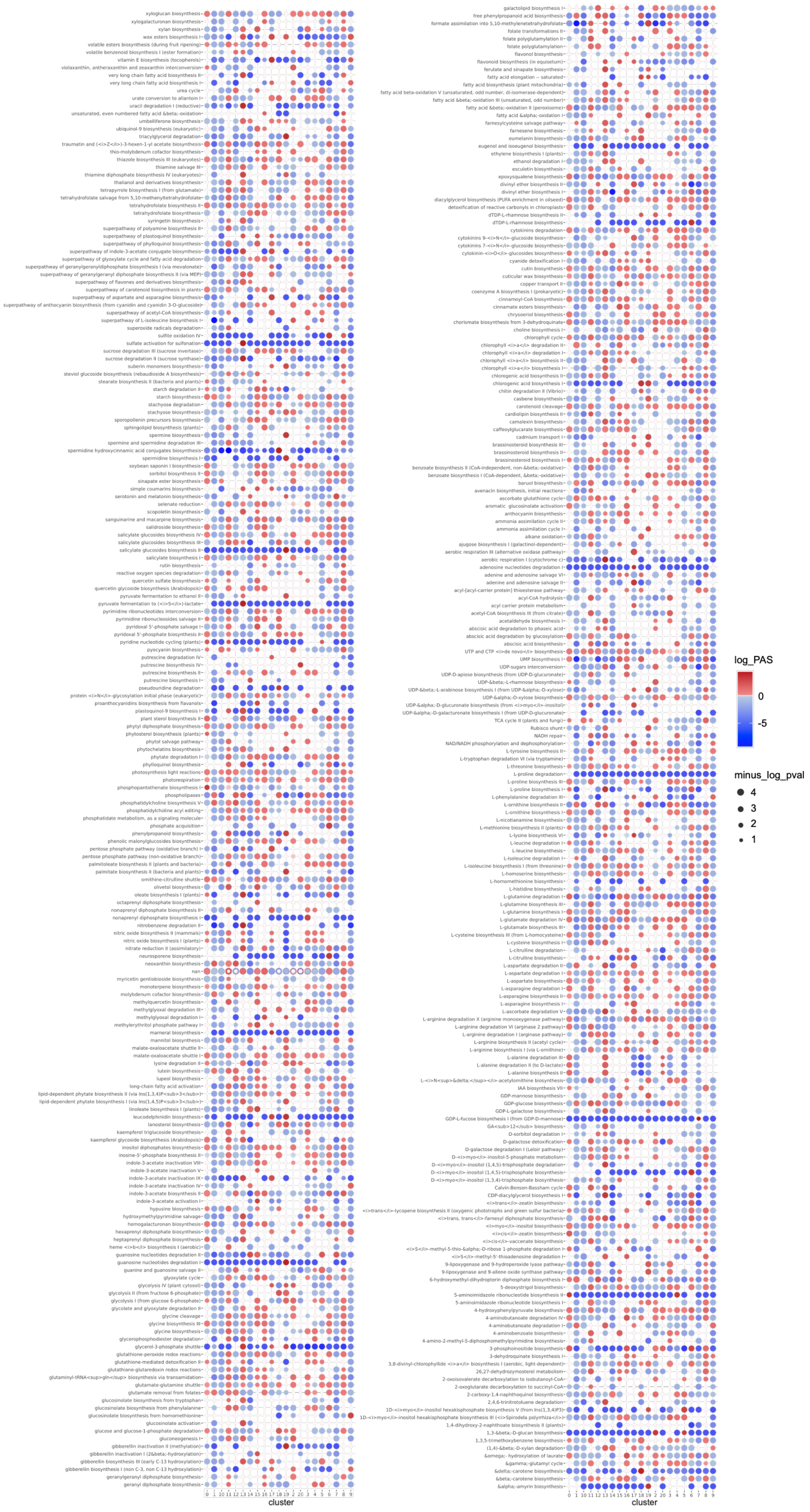


**Supporting Information Figure S4: Metabolic pathway activity across all seed cell types.** Colors represent pathway activity score (PAS). A score <1 (blue) reflects a lower than average activity of the pathway in the given cell type, a score >1 (red) indicates a higher activity. Statistical significance is presented as differences in dot size.

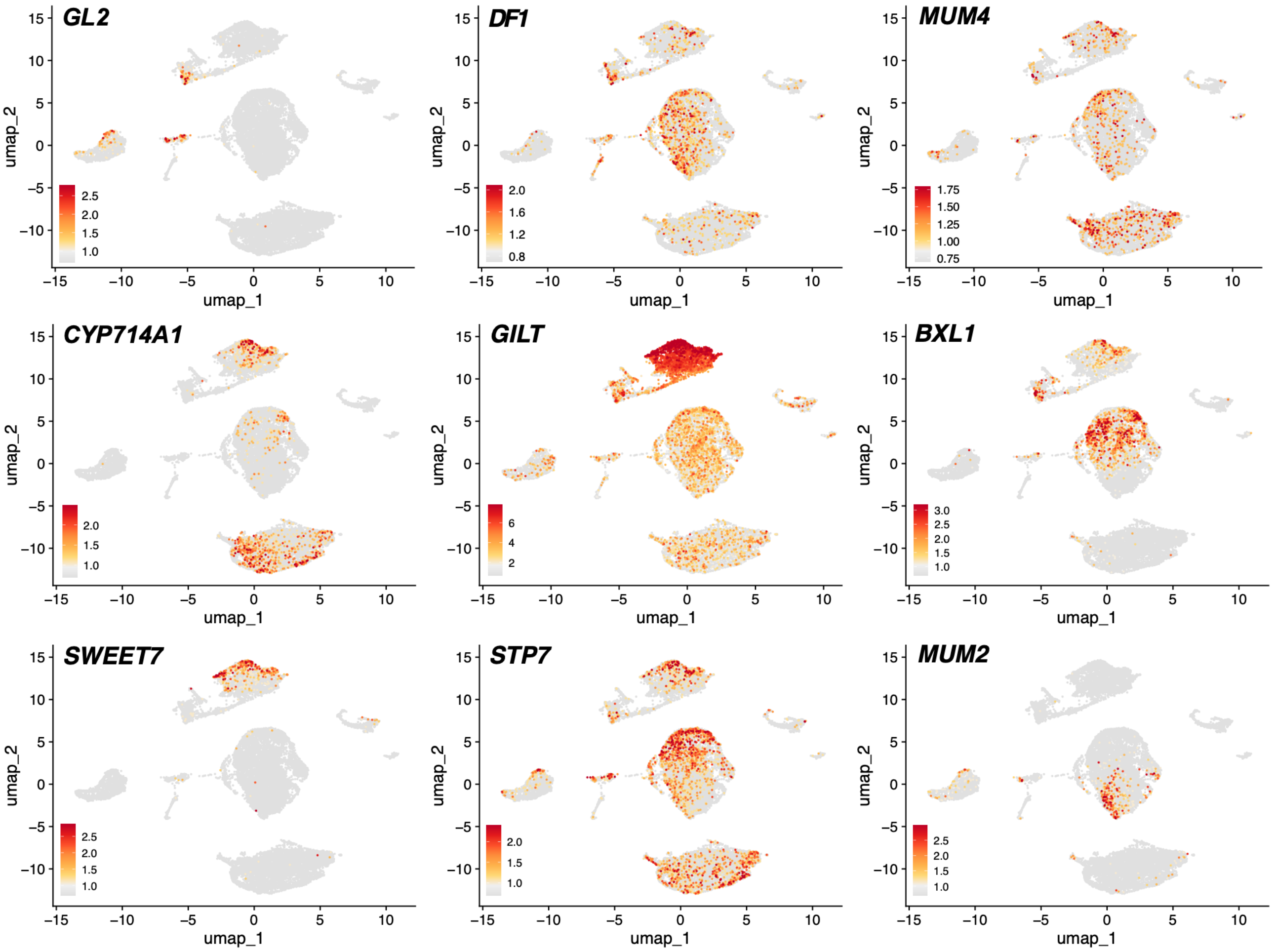


**Supporting Information Figure S5: Enrichment of transcripts in the epidermal clusters (supporting Figure 2).** UMAP plots showing enrichment of *GL2*, *MUM4*, *DF1* and *CYP714A1*, *GILT*, *BXL1*, *SWEET7*, *STP7,* and *MUM2* in the oi2 and oi1, respectively.

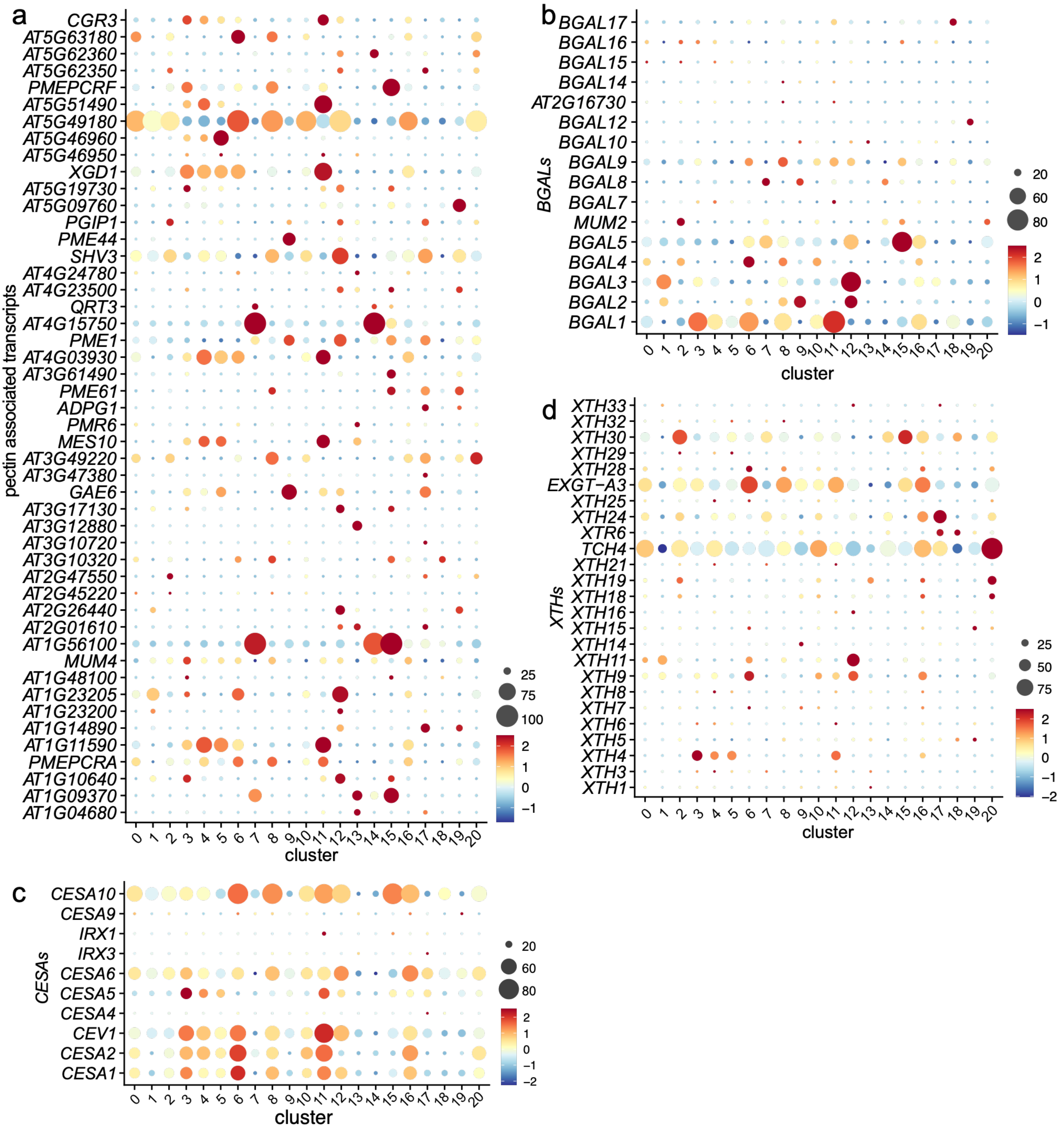


**Supporting Information Figure S6: Gene family enrichment analysis of transcripts associated with cell wall biogenesis (supporting Figure 2).** Dot plots showing transcript enrichments of pectin associated transcripts (a), members of *BGAL* (b), *CESA* (c) and *XTH* (d) gene families. The diameter of the dot indicates the percentage of cells within a class, while the color encodes average enrichment across all cells within a class.

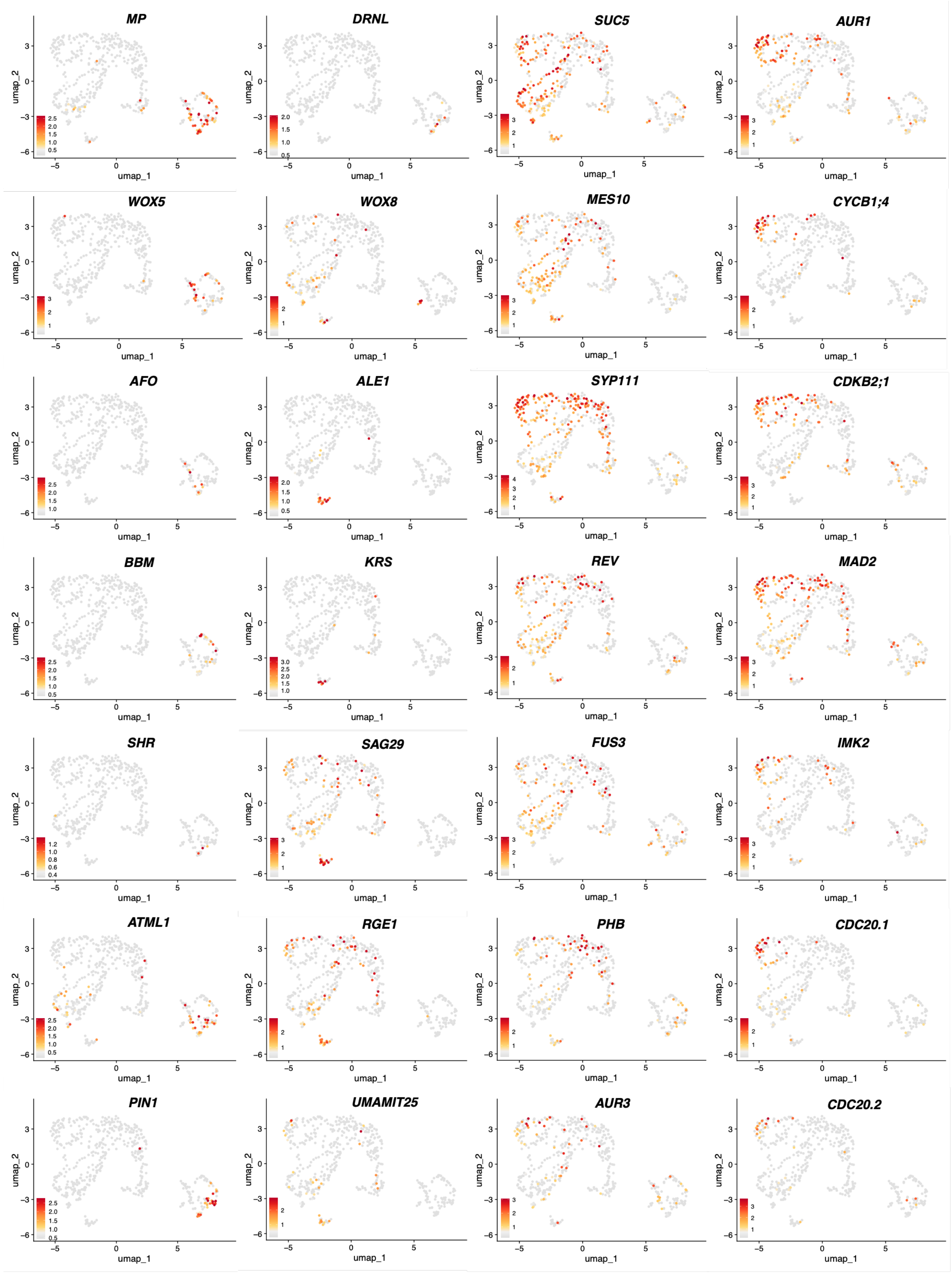


**Supporting Information Figure S7: Enrichment of transcripts in the embryo and endosperm subclusters (supporting Figure 3).** UMAP plots of AZ subcluster transcripts described in main Figure 3.

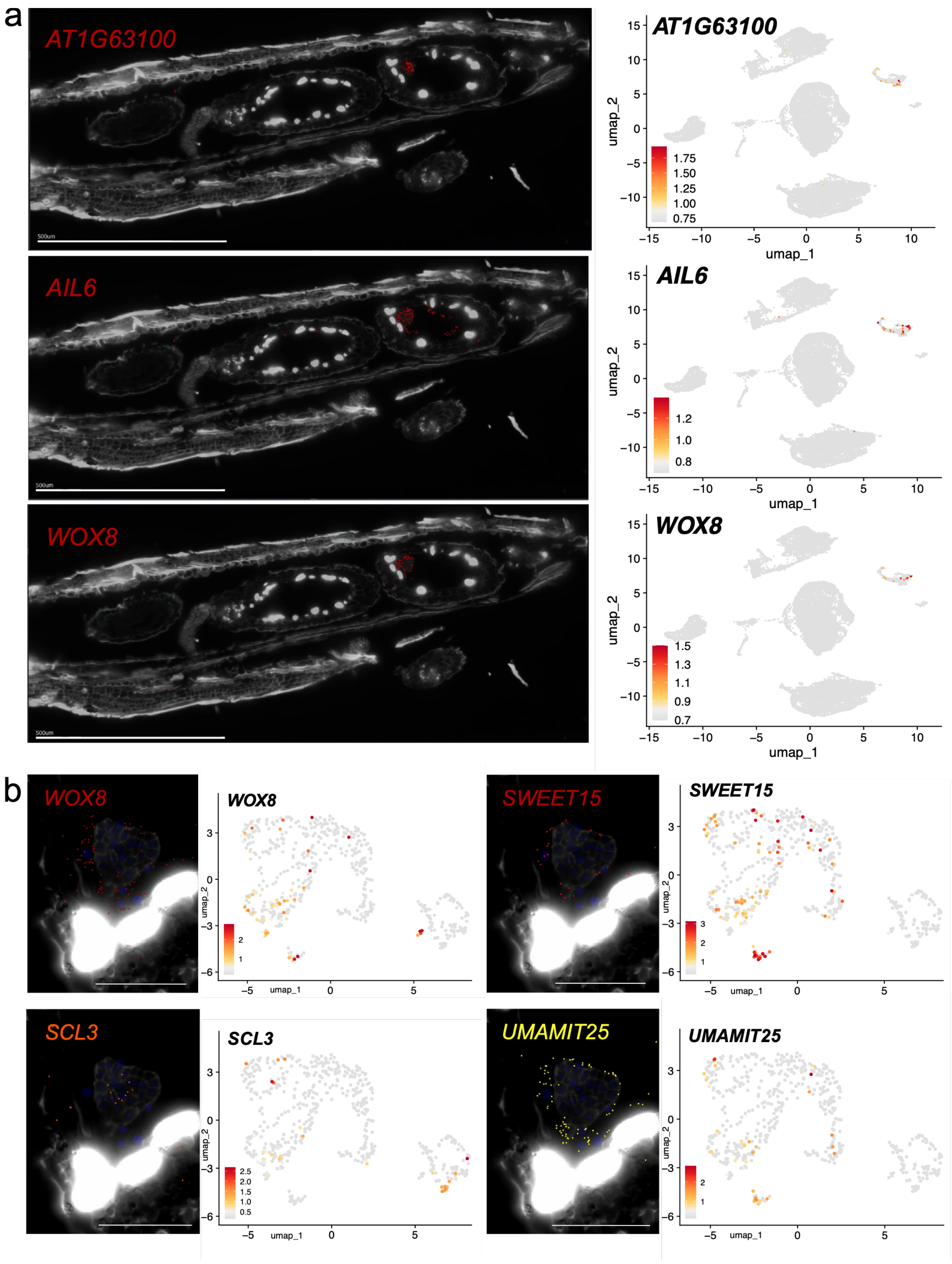


**Supporting Information Figure S8: Embryo and endosperm specific transcripts in spatial transcriptomics and scRNA-seq.** (a) Multiplexed *in situ* hybridization images showing spatial localization of marker transcripts in the embryo and endosperm (left) and corresponding transcript enrichment in scRNA-seq global UMAP plot (right). (b) Two collumns of multiplexed *in situ* hybridization images close-ups showing spatial localization of marker transcripts in the embryo and endosperm (left) and corresponding transcript enrichment in scRNA-seq cluster 13 subcluster UMAP plot (right). Scale bars (a) 500 µm, (b) 50 µm.

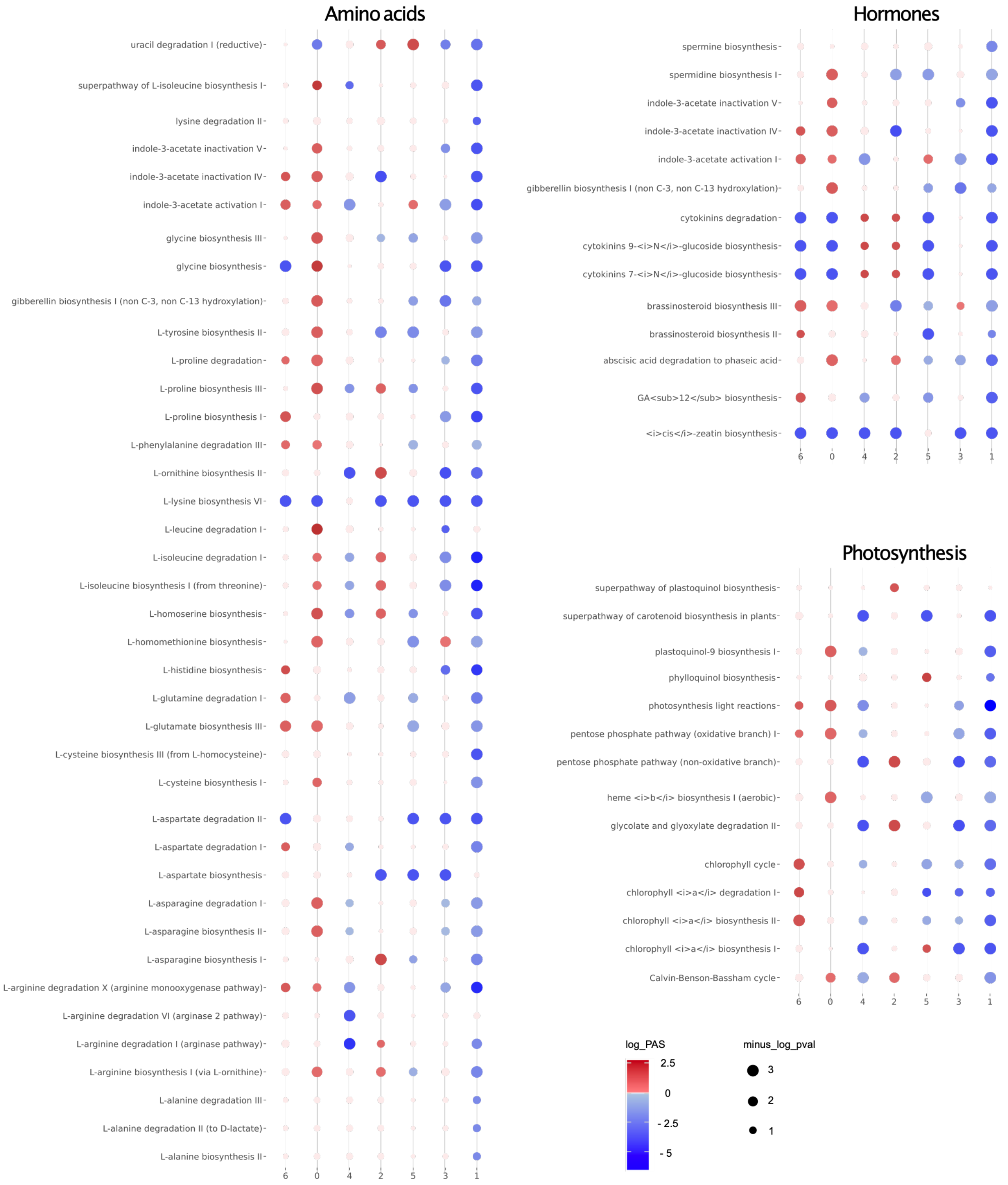


**Supporting Information Figure S9: Hormone and amino acid pathway activity across all embryo and endosperm subclusters.** Colors represent pathway activity score (PAS). A score <1 (blue) reflects a lower than average activity of the pathway in the given cell type, a score >1 (red) indicates a higher activity. Statistical significance is presented as differences in dot size.

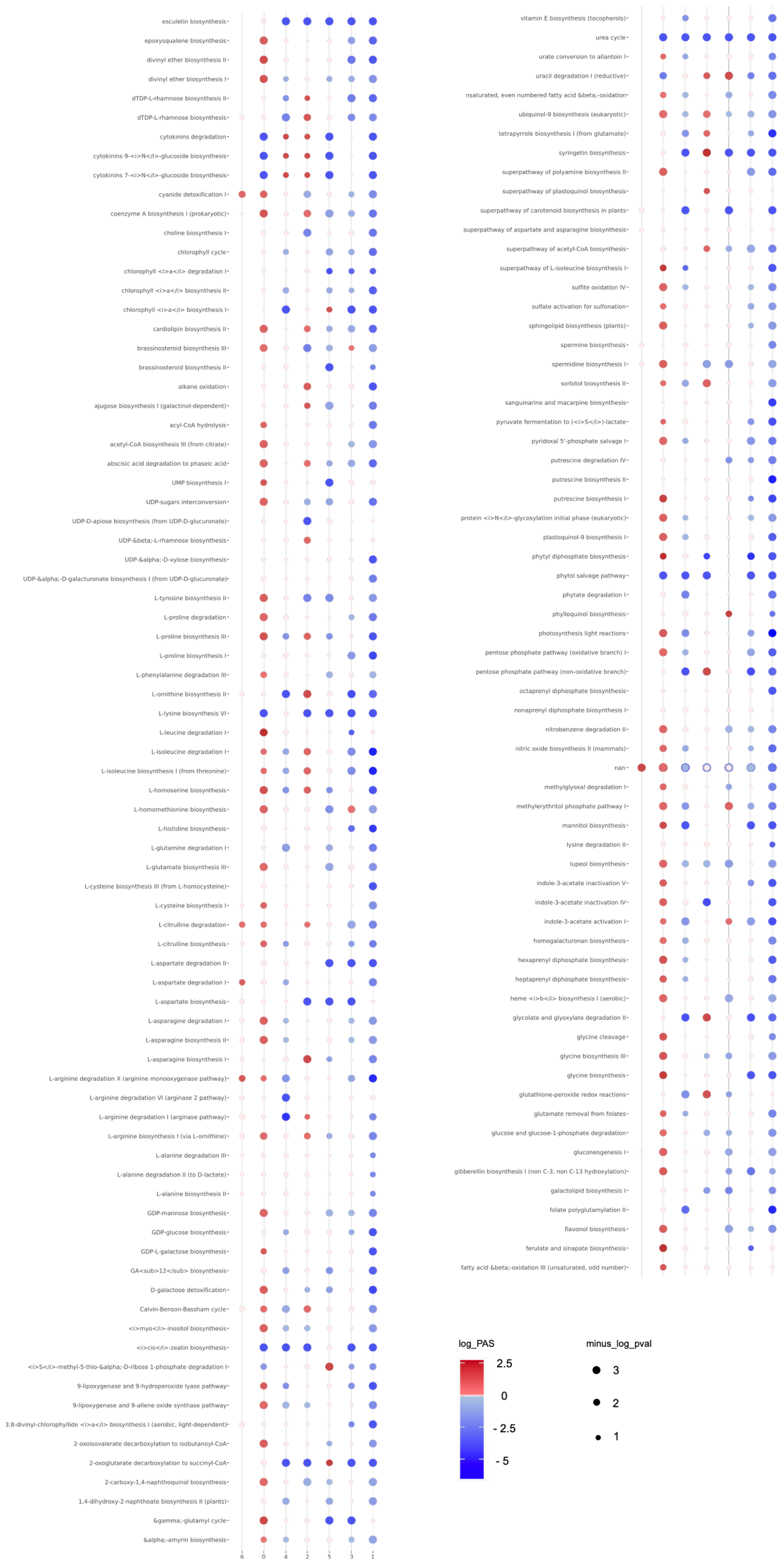


**Supporting Information Figure S10: Metabolic pathway activity across all embryo and endosperm subclusters.** Colors represent pathway activity score (PAS). A score <1 (blue) reflects a lower than average activity of the pathway in the given cell type, a score >1 (red) indicates a higher activity. Statistical significance is presented as differences in dot size.

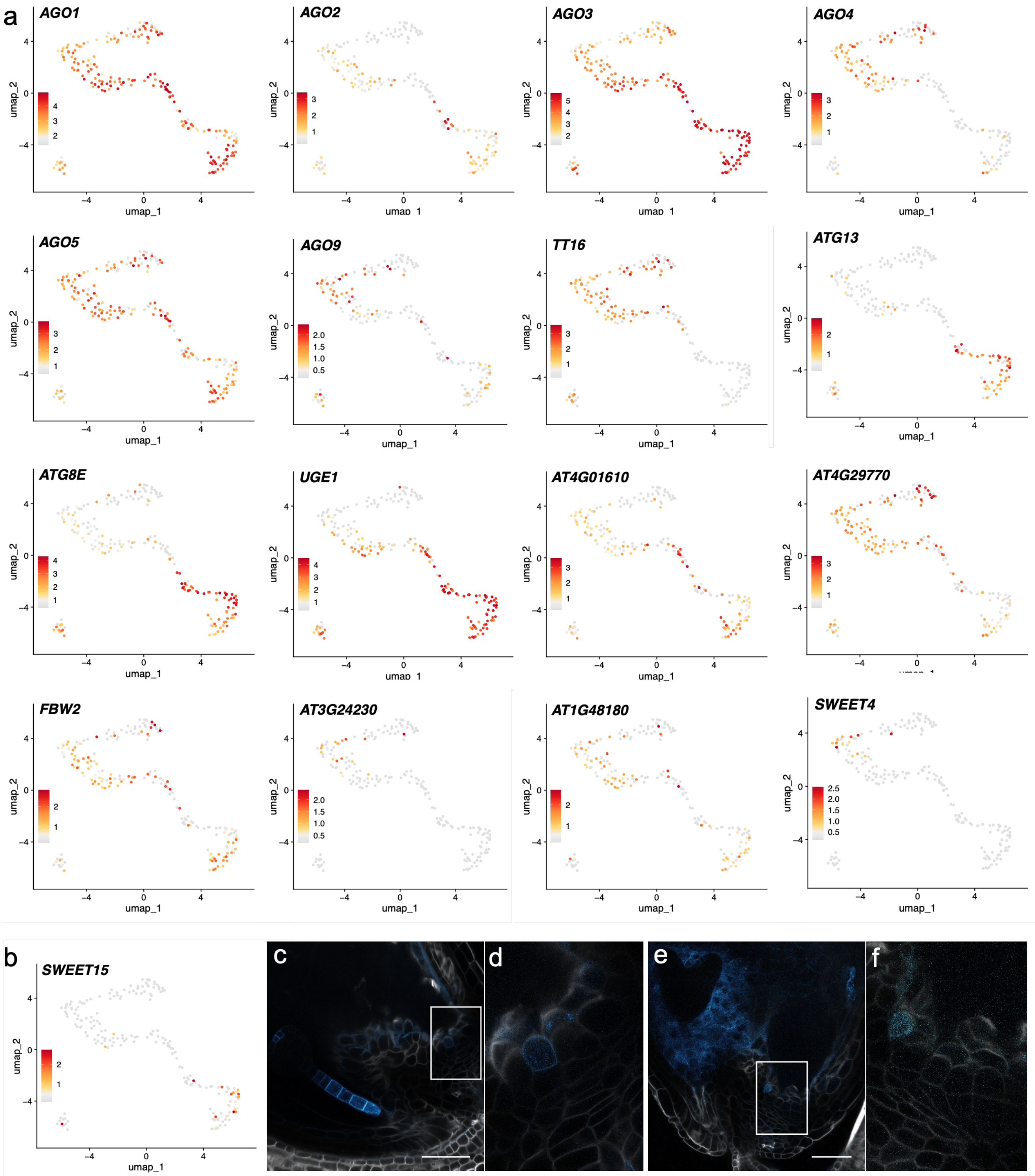


**Supporting Information Figure S11: Enrichment of transcripts in the nucellus (supporting figure 4) and single-cell transcript enrichment predicts undescribed protein domain of SWEET15 in the transient nucellus.** (a) UMAPs of nucellus subcluster transcripts described in main figure 4. (b) *SWEET15* transcripts are enriched in a subpopulation of the transient nucellus cluster. (c) Confocal microscopy image of a chemically fixed and cleared *Arabidopsis thaliana* seed of early heart developmental stage expressing *pSWEET15:SWEET15-eGFP* (blue). Hemicellulose in cell walls is stained with SR2200 (white) (d) inset of (c) showing fluorescence in the transient nucellus (e) Confocal microscopy image of a chemically fixed and cleared *Arabidopsis thaliana* seed of late heart developmental stage expressing *pSWEET15:SWEET15-eGFP* (blue). Hemicellulose in cell walls is stained with SR2200 (white) (f) inset of (e) showing fluorescence in the transient nucellus. Scale bars 50 µm.

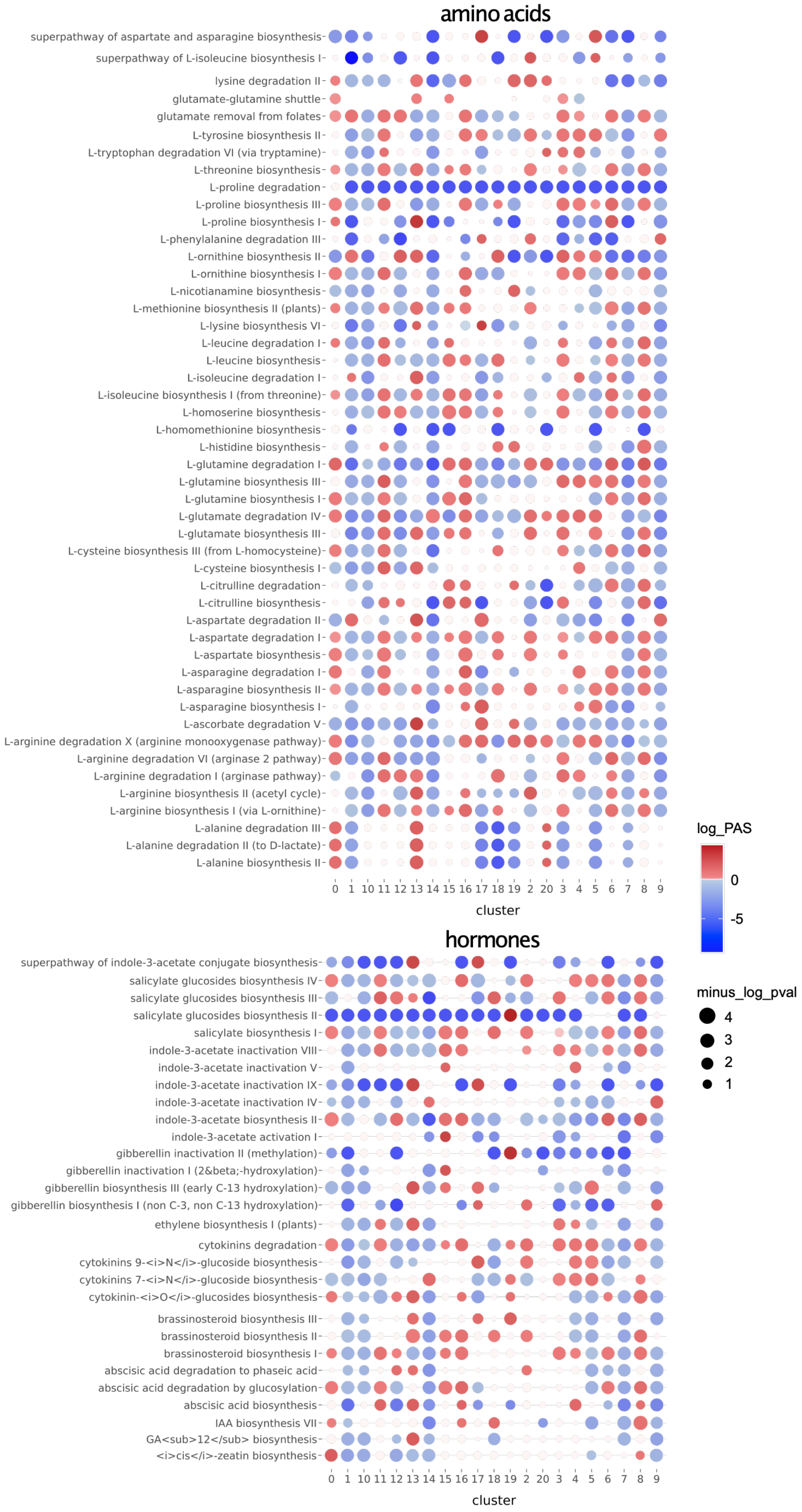


**Supporting Information Figure S12: Hormone and amino acid pathway activity across all seed cell types.** Colors represent pathway activity score (PAS). A score <1 (blue) reflects a lower than average activity of the pathway in the given cell type, a score >1 (red) indicates a higher activity. Statistical significance is presented as differences in dot size.

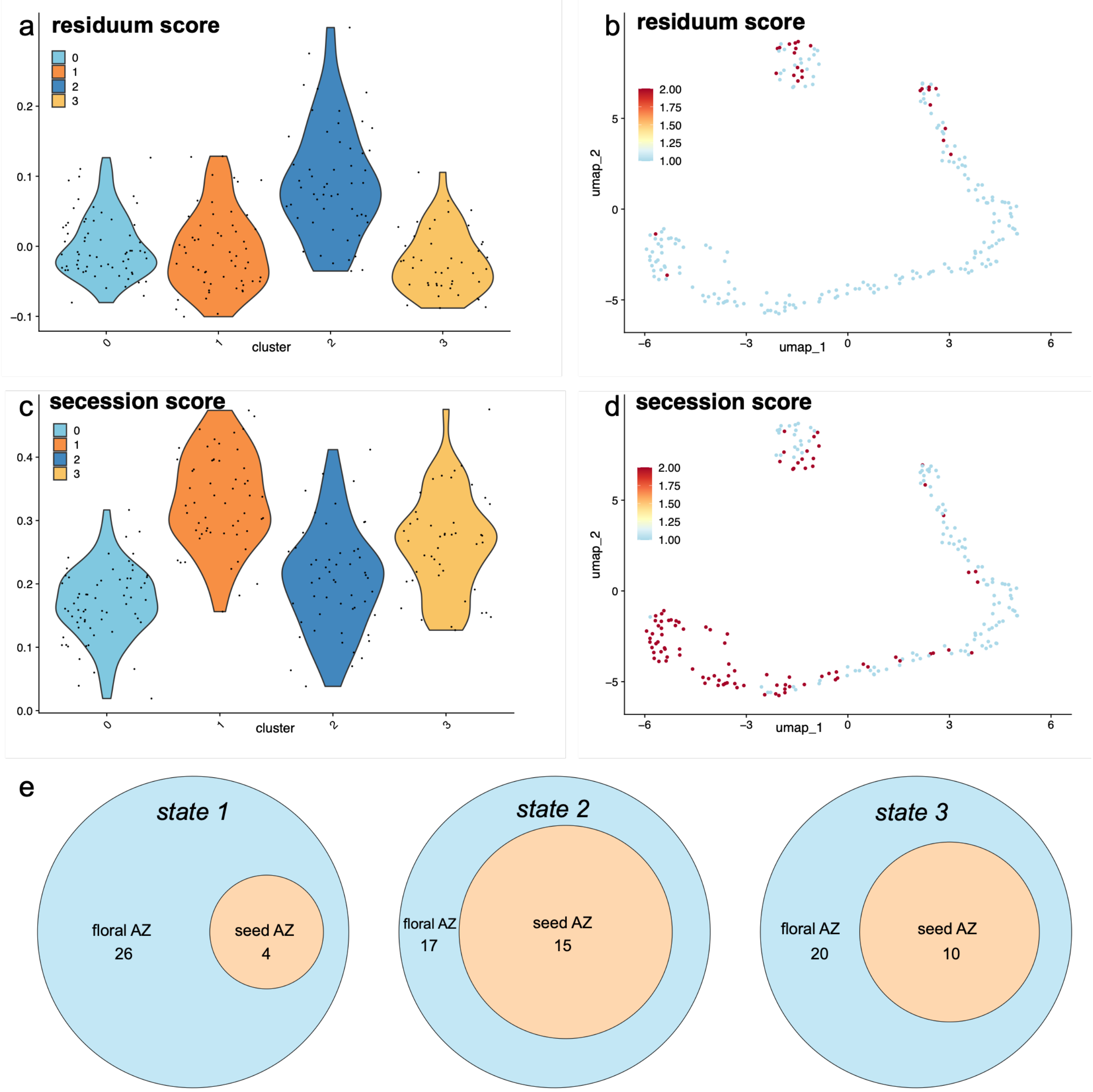


**Supporting Information Figure S13: Comparing seed and floral abscission zone transcriptomes.** (a-d) Module score mapping of the top 500 (sorted by p-value) residuum (a, b) and secession (c, d) cell markers from Lee et al. 2018 onto the seed AZ subclusters, as violin plots (a, c) and UMAP plots (b, d). (e) Venn diagrams showing the detection of transcripts in the seed AZ (orange, detected in at least 10% of AZ cluster 17 cells, at least 2 logfold enrichment) described to have a differentiation state specificity from 1 to 3 in the floral AZ residuum cells according to Wen et al. 2025 (blue).

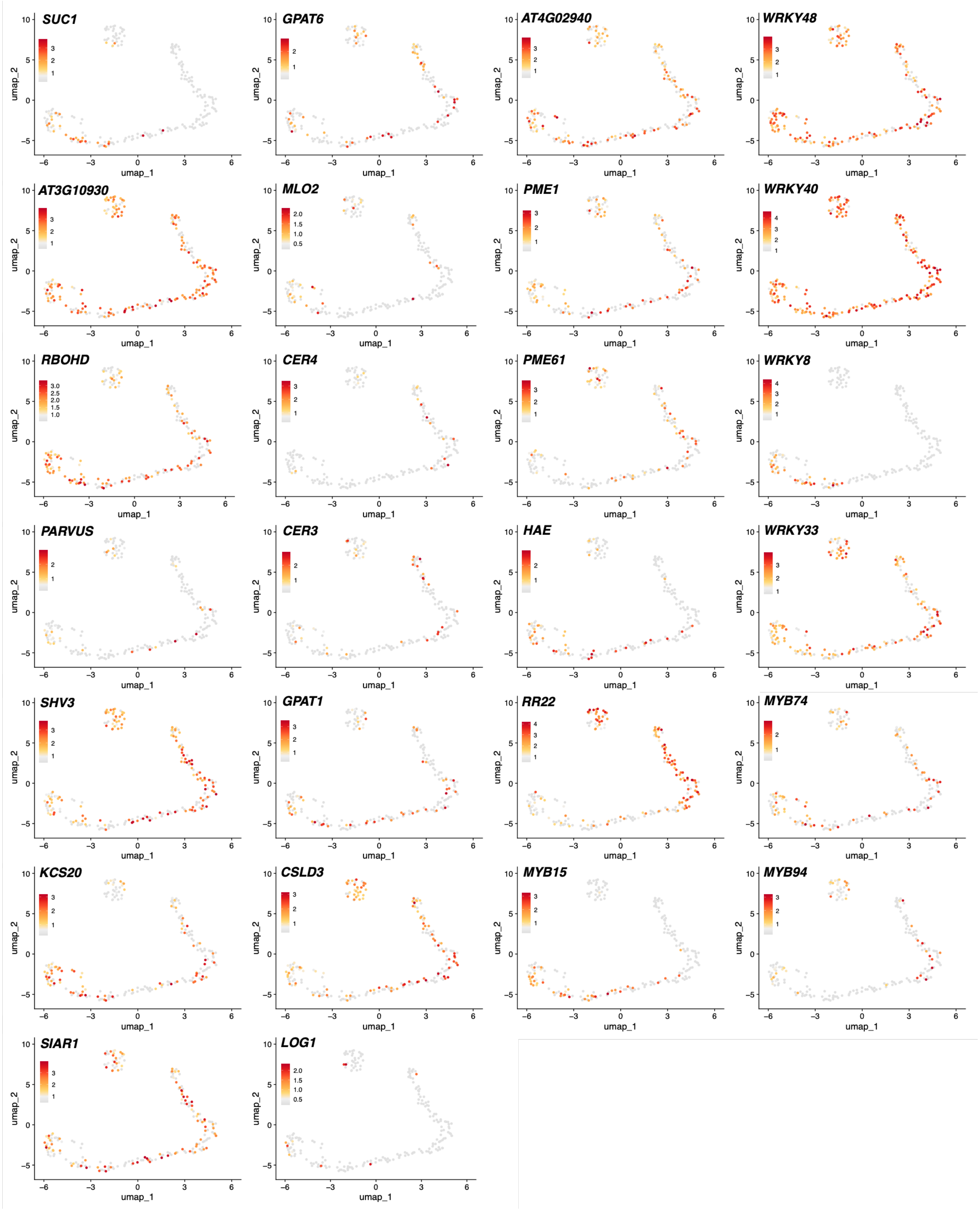


**Supporting Information Figure S14: Enrichment of transcripts in the seed abscission zone (supporting Figure 5).** UMAP plots of AZ subcluster transcripts described in the main Figure 5.
