## Supplementary Note for "Cell identities along the proximal-distal and micropylar-chalazal axes in the Arabidopsis heart-stage seed"

### Supplemental information

#### Extended Material and Methods

##### Plant growth and transgenic plant material

Arabidopsis plants were grown in soil in long day conditions (8 h dark, 16 h light) at a PAR of 100 µmol m−2 s−1, 23-24 °C (day), 20-21 °C (night) and 70% humidity. The promoter reporter line *pbZIP9:GUS-GFP* was employed to identify ULD cells in the protoplast population and validate the ULD cluster (Kim et al. 2021). To identify micropylar cells in the protoplast population and validate the micropylar oi1 cluster, *pSWEET12:SWEET12-GFP* was used (Chen et al. 2015). The reporter line *pSWEET15:SWEET15-GFP* was microscopically characterized following the detection of transcript enrichment in the nucellus (Chen et al. 2015).

##### Enrichment of seed protoplasts

Protoplasts were isolated from 5-week-old Arabidopsis Col-0 plants grown in long day conditions (see plant growth and transgenic plant material). For protoplast isolation, siliques were opened along the replum and developing Arabidopsis seeds were individually detached from their respective funiculi using forceps. For replicate 1, 90 siliques from 16 individual plants were dissected, for replicate 2, 76 siliques from 11 individual plants were dissected, totaling 166 siliques isolated from 27 individual plants. Seeds were placed immediately into buffer (0.7 M mannitol, 20 mM MES pH 5.7) prior to adding freshly prepared digestion solution (10 mg/mL Cellulase (Worthington, Lakewood), 5 mg/mL Macerozyme R-10, (Duchefa, Haarlem), 15 mg/mL Driselase Basidiomycetes sp. (Sigma-Aldrich, München), 10 mg/mL Pectolyase Y-23 (Duchefa, Haarlem), 0.7 M mannitol, 20 mM MES pH 5.7, 5 mM CaCl2, 0.035% ß-mercaptoethanol, 0.1% bovine serum albumin, (BSA)). To enhance tissue accessibility of cell wall digestion enzymes, the seed suspension was sheared by pipetting and subjected to three times of 5 min vacuum with vacuum release in between (vacuum pump Fluke 6899, SpeedVac Concentrator SPD111V Savant, Thermo Fisher). We mechanically disrupted the seed coats by shearing the seed suspension by passing repeatedly (5x) through a 1 mL pipette tip prior to incubation on an orbital shaker at 50 rpm for 35 min. Samples were passed through a 1 mL pipette tip again 5x to increase protoplast detachment and incubated for an additional 35 min. Using a light microscope, protoplast suspensions containing intact embryos in the heart stage of development were selected for subsequent wash steps. We used two types of wash buffers to remove undesired debris and ruptured protoplasts in the protoplast solution, wash buffer 1 (WS1; 0.7 M mannitol, 20 mM MES pH 5.7, 0.1% BSA, 5 mM $CaCl_2$) were added and samples were filtered through a 70 µm mesh filter (Avantor, Darmstadt) prior to adding a 1:5 mix of WS1:WS2 (WS2; 0.7 M mannitol, 2 mM MES pH 5.7, 0.1% BSA). Samples were centrifuged (Hermle Z 446 K, Gosheim) at 250 g for 2:30 min, soft de/acceleration (setting 1) and the supernatant was discarded. Washing was repeated with WS1:WS2, followed by WS2, with 2:50 min of centrifugation. The volume of washed protoplast suspension was adjusted to 620 µL with WS2.

##### ScRNA-seq and analysis

For single-cell RNA sequencing, an approximate manual count of 20.000 viable protoplasts (620 µL of protoplast suspension) was submitted to the sequencing facility of the Genomics & Transcriptomics Laboratory (HHU) and loaded into the BD Rhapsody cartridge and processed according to the BD Rhapsody user guide (BD Rhapsody 2019). cDNA amplification and sequencing on the illumina NextSeq 1.000 was performed by the Genomics & Transcriptomics Laboratory (HHU). Raw fastq files were mapped to the Arabidopsis Col-0 genome from TAIR10,

Araport11 using the BD Rhapsody pipeline version 2.2.1 with default parameters. Mitochondrial and chloroplast genes were renamed to ATMG and ATCG annotations, except for genes with a genomic copy. Recursive Substitution Error Corrected (RESC) corrected and unfiltered BD Rhapsody output files (matrix.mtx, barcode.tsv, and features.tsv) were used as input for CellBender version 0.3.2 using default parameters with 150 epochs. Transcript output (.h5 format) of cells with a higher than 50% posterior probability of containing cells were retained and converted in a Seurat compatible .h5 format using pytables with default parameters except –complevel 5. The resulting filtered h5 were analyzed in R version 4.4.1-2 and RStudio version 02/09/2025 using the Seurat package version 5. For downstream analysis, only cells with more than 300 expressed genes, counting genes detected in at least 3 cells, more than 600 transcripts per cell, and fewer than 5% mitochondrial gene transcripts were retained. Raw cell transcript counts were normalized with SCT transformation. Potential doublet cells were removed using DoubletFinder version 04/10/2024 with default parameters except parameters k = 0.01 and a 10% estimated doublet rate. Estimated doublet rate was based on manufacturers' recommendations. Two experiments were integrated using Harmony integration. UMAP was generated retaining 50 dimensions with a resolution parameter of 0.8 and the Louvain algorithm with multilevel refinement using the igraph method. To identify differentially expressed genes, FindMarkers was applied with default settings, and markers were sorted after the adjusted p-value. Cluster marker genes were specified to contain only positive markers, filtered by an average transcript detection difference of at least logfc.threshold ≥ 0.25, transcript detection in at least 10% of cells within a cluster, while not being detected in more than 10% of cells in other clusters, and sorted after the adjusted p-value, ranging up to 0.01. Differential gene expression between clusters was visualized with dotplots, violinplots, featureplots or heatmaps using the packages Seurat and ggplot2. OpenAI GPT OSS 120B was used to correct code.

##### Subclustering analysis

Subclustering analysis was performed in the same R environment by subsetting the respective cluster. For the embryo/endosperm cluster, the functions NormalizeData, FindVariableFeatures, ScaleData, RunPCA were applied with default settings, followed by neighbor search and clustering with FindNeighbors with 10 dimensions and FindClusters with a resolution of 0.8. Subclusters were visualized with a UMAP, retaining 10 dimensions. For subclustering analysis of the seed AZ cluster 17, the same workflow was applied with 3 dimensions for FindNeighbors and a resolution of 0.5 for FindClusters. Subclusters were visualized with a UMAP, retaining 3 dimensions. For subclustering analysis of the nucellus cluster 18, the same workflow was applied with 20 dimensions for FindNeighbors and a resolution of 0.5 for FindClusters. Subclusters were visualized with a UMAP, retaining 4 dimensions.

##### GO term enrichment analysis

To explore cluster identities beyond cell type, we used the normalized RNA assay and employed the FindVariableFeatures, FindAllMarkers and enrichGO functions from the Seurat package. Only positive markers with an adjusted p-value above $p < 0.05$, calculated with Benjamini-Hochberg correction, min.pct of 0.1 and a logfc.treshold over 1.5 were selected. Gene symbols of the cluster specific markers were converted to TAIR identifiers using bitr and subsequently used as an input for the enrichGO function of the clusterProfiler package v4.14.6 after (Xu et al. 2024) using default parameters. Additionally, set-based comparisons of GO terms between selected clusters were performed. OpenAI GPT OSS 120B was used to correct code.

##### Pathway activities in different cell types

Pathway activity scores, depending on transcript levels of respective constitutive genes, were calculated following (Xiao et al. 2019) and (Kim et al. 2021) using the AraCyc pathways 17.2.0. Taking the Seurat normalized RNA read count as input, let as denote the count of gene *i* in cell *k* as $g_{i,k}$. The mean transcript level $E_{i,j}$ of gene *i* in cell cluster *j* is defined as $E_{i,j} = \frac{1}{n_j}\sum_{k=1}^{n_j} g_{i,k}$, where $n_j$ is the number of cells in cluster *j* and *k* is the index of cells. $E_{i,j}$ is further normalized to become the relative transcript levels $r_{i,j}$,

$$r_{i,j} = \frac{E_{i,j}}{\frac{1}{N}\sum_{\alpha=1}^{N} E_{i,\alpha}}$$

*N* is the total number of clusters and *α* is the index for clusters. The pathway activity score (PAS) of pathway *t* in cluster *j*, denoted as $p_{t,j}$, with

$$p_{t,j} = \frac{\sum_{i=1}^{m_t} w_i\, r_{i,j}}{\sum_{i=1}^{m_t} w_i}$$

where $m_t$ is the number of genes in pathway *t*, and $w_i$ is the weight of gene *i* and defined as the number of pathways that it is associated with.

The inclusion of $w_i$ in the calculation of the PAS ensures that genes associated with fewer pathways will have stronger influence on the PAS calculation. The average values of $p_{t,j}$ is 1, and $p_{t,j} > 1$ ($p_{t,j} < 1$) indicates an up-regulation (down-regulation) of the pathway *t* in cluster *j*. To assess the significance of the deviation of $p_{t,j}$ from 1, we calculated a null model, in which we calculate $p_{t,j}$ in each run with randomly permuted cell labels. Using 1000 runs, we calculated the distribution of $p_{t,j}$ in this null model and thereby the associated p-values.

**Spatial transcriptomics**

For the MERFISH spatial transcriptomics experiment, Arabidopsis Col-0 siliques with seeds in embryonal heart stage were processed according to MERSCOPE® User Guide (91600112 Rev C). Sample preparation was performed in RNase-reduced conditions at 4 °C. Silique pedicel were adhered to double sided tape and valves were opened and trimmed to 1 cm prior to immersion in freshly prepared fixing solution (4% paraformaldehyde in PBS, 0.1% Triton-X, 0.1% Tween-20). Vacuum infiltration was applied five times for 5 min each (vacuum pump Fluke 6899, SpeedVac Concentrator SPD111V Savant, Thermo Fisher) with vacuum release between cycles, or until siliques remained fully submerged. Following fixation, the solution was exchanged, and samples were incubated at 4 °C, with gentle shaking at 15 rpm overnight. Samples were subsequently washed twice in PBS for 1 h each and dehydrated through an ethanol series in 1 h increments (15%, 30%, 50%, 70% ethanol), followed by 1.5 h increments (80%, 90%, 100%, 100%) and an additional overnight incubation in 100% ethanol. Prior to embedding in molten Paraplast (Carl Roth), siliques were cleared in ROTIHistol (Carl Roth) at 20-22 °C, automized by a biosystems embedding machine (Leica), program 1. Individual siliques were placed into plastic molds before covering with molten Paraplast (Carl Roth). Prior to storage at 4 °C, samples were dried overnight in a fume hood. For sectioning, sample blocks were pre-chilled on ice, and microtome sectioning was performed at 20-22 °C. Siliques were carefully oriented parallel to the microtome blade to obtain longitudinal sections. After trimming the blocks, ribbons of 7 µM thickness were cut and collected. Sections containing longitudinal silique sections were microscopically identified and mounted onto MERFISH slides prepared according to the manufacturer's instructions (MERSCOPE® User Guide, 91600112 Rev C). Residual mounting liquid was removed and evaporated at 37 °C, followed by baking the sections to the slide at 55 °C for 15 min, according to manufacturer's instructions (MERSCOPE® User Guide, 91600112 Rev C). Slides were stored at – 20 °C prior to processing at Vizgen headquarters (Cambridge, USA). From deparaffinization on, samples were processed internally at Vizgen Headquarters according to the manufacturer's instructions. Spatial transcriptomics data was visualized in the MERSCOPE Visualizer software.

**Confocal imaging**

To enable the detection of fluorescence with cellular resolution in deeper cell layers of developing Arabidopsis seeds, samples were subjected to optical clearing prior to imaging. Developing seeds were excised from siliques and immediately submerged in fixation solution (4% Paraformaldehyde solution in PBS (Thermo Scientific Chemicals), 0.1% Tween 20 (Sigma-Aldrich), 0.1% Triton X-100 (Sigma-Aldrich)) on ice. Vacuum infiltration was performed through five cycles of vacuum application and release, each lasting 5 min, after which the fixation solution was replaced. Seeds were incubated in fixation solution for 10-16 h at 4 °C in darkness with gentle shaking on an orbital shaker at 30-50 rpm. Following fixation, samples were washed three times in 1X PBS, shaking at 30-50 rpm for 1 h each at 4 °C. Washing was repeated twice. Optical clearing was performed using a modified ClearSeeAlpha protocol based on (Kurihara et al. 2021; Attuluri et al. 2022). After removal of PBS, seeds were incubated in ClearSeeAlpha at 20-22 °C, shaking on an orbital shaker at 30-50 rpm for a minimum of 7 days and up to 40 days. ClearSeeAlpha was exchanged every other day for the first two weeks of clearing, subsequently twice per week. One day prior to CLSM, seeds were incubated for a minimum of 5 h and up to 20 h in ClearSeeAlpha supplied with 0.1% (v/v) solution of Renaissance SR2200 (Renaissance Chemicals, UK). To decrease background fluorescence, seeds were washed with ClearSeeAlpha for at least 1 h prior to imaging.

Arabidopsis seeds, siliques, and protoplasts were mounted onto slide glasses in their respective mounting media. Double-sided tape was used as a spacer between the slide and cover glass to prevent sample compression. Whole seeds and protoplasts were imaged by confocal laser scanning microscopy (CLSM) using a Leica TCS SP8 microscope equipped with 20x and 40x water immersion objectives. Whole seeds were stained as described above and protoplast suspensions were either imaged directly or incubated for 5 min with 5 µg/mL FM4-64FX (Sigma-Aldrich) for staining the plasma membrane. GFP, FM4-64FX, SR2200 were detected using the following setting ranges: GFP excitation 388 nm, emission 498-525 nm using the HyD 2 detector; FM4-64FX excitation 561 nm, emission 575-640 nm using the HyD 4 detector; SR2200 excitation 405 nm, emission 436-475 nm using the HyD 2 detector. Time gating was used in several cases to reduce autofluorescence from seed tissues, gating from around 0.3-1 ns to 4.5-8.5 ns. Transmitted light images were captured using the TLD with the PMT detector. Data was processed in ImageJ2 version 2.16.0/1.54p.

**Data availability**

The raw and processed sequencing data are available at Gene Expression Omnibus (GEO) (www.ncbi.nlm.nih.gov/geo/) under accession number X.
