## Supplementary Table and Script for "Cell identities along the proximal-distal and micropylar-chalazal axes in the Arabidopsis heart-stage seed": Supporting Information Note 2 R code for scRNAseq analysis.R

```
# This file contains the R script used to process and analyze the scRNA-seq datasets in Rombi et al. 2026. 

# loading libraries prior to starting script-----
# The respective packages need to be installed once prior to library loading.
options(future.globals.maxSize = 1e11)
library(plotly)
library(Seurat)
library(hdf5r)
library(DoubletFinder)
library(SeuratData)
library(ggplot2)
library(patchwork)
library(dplyr)
library(sctransform)
library(glmGamPoi)
library(harmony) 
library(reticulate)
library(plotly)
library(ComplexHeatmap)
library(presto)
library(tictoc)
library(magick)
library(harmony)
library(Matrix)
library(Seurat)
library(dplyr)
library(clusterProfiler)
library(org.At.tair.db)
library(enrichplot)
library(ggplot2)
library(RColorBrewer) 


# Data processing and integration of 2 replicates---------
# seed replicate 1
setwd("c:/Work/Collabs/Khadi/CB_4th/")
seed.data <- Read10X_h5(filename = "default.output_filtered.seurat.h5", use.names = TRUE)
# the Seurat object was initialized with the raw (non-normalized) data.
seed1 <- CreateSeuratObject(counts = seed.data, project = "Seed1", min.cells = 3, min.features = 200)
seed1[["percent.mt"]] <- PercentageFeatureSet(seed1, pattern = "^ATMG")
seed1.1g_2t <- subset(seed1, subset = nFeature_RNA > 1000 & nFeature_RNA < 10000 & nCount_RNA > 2000 & nCount_RNA < 100000 & percent.mt < 5)

# QC metrics as violin plots
VlnPlot(seed1.1g_2t, features = c("nFeature_RNA", "nCount_RNA", "percent.mt"), ncol = 3)
plot1 <- FeatureScatter(seed1.1g_2t, feature1 = "nCount_RNA", feature2 = "percent.mt")
plot2 <- FeatureScatter(seed1.1g_2t, feature1 = "nCount_RNA", feature2 = "nFeature_RNA")
plot1 + plot2

rm(seed1)
rm(seed.data)
seed1.1g_2t

# seed replicate 2
setwd("c0:/Work/Collabs/Khadi/rep2/")
seed.data2 <- Read10X_h5(filename = "output.default.rep2_filtered_seurat.h5", use.names = TRUE)
seed2 <- CreateSeuratObject(counts = seed.data2, project = "Seed2", min.cells = 3, min.features = 200)
seed2[["percent.mt"]] <- PercentageFeatureSet(seed2, pattern = "^ATMG")
seed2.1g_2t <- subset(seed2, subset = nFeature_RNA > 1000 & nFeature_RNA < 10000 & nCount_RNA > 2000 & nCount_RNA < 100000 & percent.mt < 5)

# QC metrics as violin plots
VlnPlot(seed2.1g_2t, features = c("nFeature_RNA", "nCount_RNA", "percent.mt"), ncol = 3)
plot1 <- FeatureScatter(seed2.1g_2t, feature1 = "nCount_RNA", feature2 = "percent.mt")
plot2 <- FeatureScatter(seed2.1g_2t, feature1 = "nCount_RNA", feature2 = "nFeature_RNA")
plot1 + plot2

rm(seed2)
rm(seed.data2)
seed2.1g_2t

# DoubletFinder was used to predict and remove putative doublets.
# DoubletFinder was used for both reps separately before merging
seed1.1g_2t <- SCTransform(seed1.1g_2t, vars.to.regress = "percent.mt", verbose = F, method = "glmGamPoi")
seed1.1g_2t <- RunPCA(seed1.1g_2t)
ElbowPlot(seed1.1g_2t, ndims = 50)
seed1.1g_2t <- FindNeighbors(seed1.1g_2t, dims = 1:40)
seed1.1g_2t <- FindClusters(seed1.1g_2t, resolution = 0.8, algorithm = 2)
seed1.1g_2t <- RunUMAP(seed1.1g_2t, dims = 1:40)

# pK Identification (no ground-truth)
sweep.res.list_seed1 <- paramSweep(seed1.1g_2t, PCs = 1:40, sct = T)
sweep.stats_seed1 <- summarizeSweep(sweep.res.list_seed1, GT = FALSE)
bcmvn_seed1 <- find.pK(sweep.stats_seed1)

ggplot(bcmvn_seed1, aes(pK, BCmetric, group = 1))+
  geom_point() +
  geom_line()
bcmvn_seed1

# finding the optimal (=max) pK value
pK <- bcmvn_seed1 %>%
  filter(BCmetric == max(BCmetric)) %>%
  select(pK)
pK <- as.numeric(as.character(pK[[1]]))

# Homotypic Doublet Proportion Estimate
annotations <- seed1.1g_2t@meta.data$seurat_clusters
homotypic.prop <- modelHomotypic(annotations) 
nExp_poi <- round(0.075*nrow(seed1.1g_2t@meta.data))  # Assuming 7.5% doublet formation rate 
nExp_poi.adj <- round(nExp_poi*(1-homotypic.prop))

# DoubletFinder was run with varying classification stringencies
seed1.1g_2t <- doubletFinder(seed1.1g_2t, PCs = 1:40, pN = 0.25, pK = pK, nExp = nExp_poi.adj, reuse.pANN = FALSE, sct = T)

#Second Replicate DoubletFinder
seed2.1g_2t <- SCTransform(seed2.1g_2t, vars.to.regress = "percent.mt", verbose = F)
seed2.1g_2t <- RunPCA(seed2.1g_2t)
ElbowPlot(seed2.1g_2t, ndims = 50)
seed2.1g_2t <- FindNeighbors(seed2.1g_2t, dims = 1:40)
seed2.1g_2t <- FindClusters(seed2.1g_2t, resolution = 0.8, algorithm = 2)
seed2.1g_2t <- RunUMAP(seed2.1g_2t, dims = 1:40)

# pK Identification (no ground-truth)
sweep.res.list_seed2 <- paramSweep(seed2.1g_2t, PCs = 1:40, sct = T)
sweep.stats_seed2 <- summarizeSweep(sweep.res.list_seed2, GT = FALSE)
bcmvn_seed2 <- find.pK(sweep.stats_seed2)

ggplot(bcmvn_seed2, aes(pK, BCmetric, group = 1))+
  geom_point() +
  geom_line()
bcmvn_seed2

# finding the optimal (=max) pK value for rep2
pK2 <- bcmvn_seed2 %>%
  filter(BCmetric == max(BCmetric)) %>%
  select(pK)
pK2 <- as.numeric(as.character(pK2[[1]]))

# Homotypic Doublet Proportion Estimate
annotations2 <- seed2.1g_2t@meta.data$seurat_clusters
homotypic.prop2 <- modelHomotypic(annotations2) 
nExp_poi2 <- round(0.075*nrow(seed2.1g_2t@meta.data))  ## Assuming 7.5% doublet formation rate - tailor for your dataset
nExp_poi.adj2 <- round(nExp_poi2*(1-homotypic.prop2))

# DoubletFinder was run with varying classification stringencies
seed2.1g_2t <- doubletFinder(seed2.1g_2t, PCs = 1:40, pN = 0.25, pK = pK2, nExp = nExp_poi.adj2, reuse.pANN = FALSE, sct = T)

#Visualization of Doublets and sub-setting only singlets
DimPlot(seed1.1g_2t, pt.size = 1, label = T, label.size = 7, group.by = "DF.classifications_0.25_0.26_549")
DimPlot(seed2.1g_2t, pt.size = 1, label = T, label.size = 7, group.by = "DF.classifications_0.25_0.01_307")

seed1.1g_2t.wd <- subset(seed1.1g_2t, subset = DF.classifications_0.25_0.26_549 == "Singlet")
seed2.1g_2t.wd <- subset(seed2.1g_2t, subset = DF.classifications_0.25_0.01_307 == "Singlet")

# Redoing normalization for both replicates
seed1.1g_2t.wd[["percent.mt"]] <- PercentageFeatureSet(seed1.1g_2t.wd, pattern = "^ATMG")
seed1.1g_2t.wd[["percent.pt"]] <- PercentageFeatureSet(seed1.1g_2t.wd, pattern = "^ATCG")
seed2.1g_2t.wd[["percent.mt"]] <- PercentageFeatureSet(seed2.1g_2t.wd, pattern = "^ATMG")
seed2.1g_2t.wd[["percent.pt"]] <- PercentageFeatureSet(seed2.1g_2t.wd, pattern = "^ATCG")

seed1.1g_2t.wd <- SCTransform(seed1.1g_2t.wd, vars.to.regress = "percent.mt", verbose = F, , method = "glmGamPoi")
seed1.1g_2t.wd <- RunPCA(seed1.1g_2t.wd)
seed1.1g_2t.wd <- FindNeighbors(seed1.1g_2t.wd, dims = 1:40)
seed1.1g_2t.wd <- FindClusters(seed1.1g_2t.wd, resolution = 0.8, algorithm = 2, method = "igraph")
seed1.1g_2t.wd <- RunUMAP(seed1.1g_2t.wd, dims = 1:40)

seed2.1g_2t.wd <- SCTransform(seed2.1g_2t.wd, vars.to.regress = "percent.mt", verbose = F, method = "glmGamPoi")
seed2.1g_2t.wd <- RunPCA(seed2.1g_2t.wd)
seed2.1g_2t.wd <- FindNeighbors(seed2.1g_2t.wd, dims = 1:40)
seed2.1g_2t.wd <- FindClusters(seed2.1g_2t.wd, resolution = 0.8, algorithm = 2, method = "igraph")
seed2.1g_2t.wd <- RunUMAP(seed2.1g_2t.wd, dims = 1:40)

DimPlot(seed1.1g_2t.wd, reduction = "umap", pt.size = 1, label = T, label.size = 7)
DimPlot(seed2.1g_2t.wd, reduction = "umap", pt.size = 1, label = T, label.size = 7)

# Replicate integration
setwd("c:/Work/Collabs/Khadi/both_reps/")
seed1.1g_2t.wd$batch <- "replicate1" 
seed2.1g_2t.wd$batch <- "replicate2" 
rep.list <- list(seed1.1g_2t.wd, seed2.1g_2t.wd)

# Anchor gene identification, SCT integration
rep.list <- PrepSCTIntegration(rep.list, assay = "SCT", anchor.features = features)
anchors <- FindIntegrationAnchors(rep.list, normalization.method = "SCT", reduction = "cca", anchor.features = features)
seed.int <- IntegrateData(anchorset = anchors, normalization.method = "SCT")
DefaultAssay(seed.int) <- "integrated"
seed.int <- ScaleData(seed.int, verbose = F)
seed.int <- RunPCA(seed.int, verbose = F)
seed.int <- RunUMAP(seed.int, dims = 1:50, reduction = "pca")
seed.int <- FindNeighbors(seed.int, dims = 1:50)
seed.int <- FindClusters(seed.int, resolution = 0.8, algorithm = 2)
DimPlot(seed.int, reduction = "umap", pt.size = 1, label = T, label.size = 7)

#Harmony integration
table(seed.int$batch) # continue if this prints the correct number of rep1 and rep2 cells
harm.int <- RunHarmony(seed.int, group.by.vars = "batch")
harm.int <- RunUMAP(harm.int, dims = 1:50, reduction = "harmony")
harm.int <- FindNeighbors(harm.int, dims = 1:50, reduction = "harmony")
harm.int <- FindClusters(harm.int, resolution = 0.8, algorithm = 2, reduction = "harmony")

DimPlot(harm.int, reduction = "umap", pt.size = 1, label = T, label.size = 7)
DimPlot(harm.int, reduction = "umap", pt.size = 1, label = TRUE, label.size = 7, group.by = "orig.ident") 
table(Idents(harm.int))


# plots ----------

my_colors <- c("#80C8E0", "#B0DAEA", "#5AA7D0", "#FABB59", "#FBE78E", "#FDF7C3", "#F4E272", "#F9A34C", "#D2E2B2", "#346BA3", "#BDDFE7", 
               "#F98F24", "#8CC8E0", "#D22B1A", "#F47B3C", "#E94D26", "#D6F4FB", "#1C094E", "#8B1A1A", "#1560BD", "#3A78A8")
DimPlot(harm.int, reduction = "umap", pt.size = 1, label = T, label.size = 7, cols = my_colors)


# feature plot
DefaultAssay(harm.int) <- "SCT"

features <- c("GL2")
seurat_colors <- c("#E0E0E0","#f0f0f0","#FED976","#FEB24C","#FD8D3C","#FC4E2A","#E31A1C","#BD0026") 
p_feature <- FeaturePlot(harm.int, features = features, order = T, min.cutoff = 'q1', max.cutoff = 'q99') + scale_color_gradientn(colors = seurat_colors)
print(p_feature)

# Dotplot
features <- c("BAN")
DefaultAssay(harm.int) <- "SCT"
p <- DotPlot(harm.int, features = features) + RotatedAxis()
seurat_colors <- rev(brewer.pal(n = 11, name = "RdYlBu"))
p <- p + scale_color_gradientn(colors = seurat_colors) + coord_flip()
print(p)


# QC plots --------
# mitochondrial, ribosomal and chloroplastidial transcript enrichments 
layer_names <- Layers(harm.int[["SCT"]])
expr.mat <- GetAssayData(harm.int[["SCT"]], layer = layer_names[1])
cp.genes <- grep("^ATCG", rownames(expr.mat), value = TRUE)
mt.genes <- grep("^ATMG", rownames(expr.mat), value = TRUE)
ribo.genes <- grep("^(RPL|RPS|PRPL|PRPS|MRPL|MRPS)", rownames(expr.mat), value = TRUE)
length(ribo.genes)
harm.int$nCount.expr   <- Matrix::colSums(expr.mat)
harm.int$nFeature.expr <- Matrix::colSums(expr.mat > 0)

harm.int$percent.cp <- Matrix::colSums(expr.mat[cp.genes, , drop = FALSE]) /
  Matrix::colSums(expr.mat) * 100

harm.int$percent.mt <- Matrix::colSums(expr.mat[mt.genes, , drop = FALSE]) /
  Matrix::colSums(expr.mat) * 100

harm.int$percent.ribo <- Matrix::colSums(expr.mat[ribo.genes, , drop = FALSE]) /
  Matrix::colSums(expr.mat) * 100

FeaturePlot(harm.int, features = c("nFeature.expr", "nCount.expr", "percent.cp", "percent.mt", "percent.ribo"), reduction = "umap", min.cutoff = "q01", max.cutoff = "q99", 
            cols = c("grey90", "#E41A1C"), ncol = 3, order = TRUE)


# number of genes
VlnPlot(harm.int,  features = "nFeature_RNA", pt.size = 0.05, group.by = "orig.ident")

# number of transcripts
VlnPlot(harm.int,  features = "nCount_RNA", pt.size = 0.05, group.by = "orig.ident")

VlnPlot(
  harm.int,
  features = "nCount_RNA",
  pt.size = 0.05,
  group.by = "orig.ident"
) +
  ggplot2::coord_cartesian(ylim = c(0, 10000))


plot_features <- VlnPlot(
  harm.int,
  features = "nFeature_RNA",
  pt.size = 0.1
) +
  scale_fill_manual(values = my_colors) +
  ggtitle("Number of Features") +
  ylab("Number of Features")

print(plot_features)


plot_features <- VlnPlot(
  harm.int,
  features = "nCount_RNA",
  y.max = 75000,
  pt.size = 0.1
) +
  scale_fill_manual(values = my_colors) +
  ggtitle("Number of Counts") +
  ylab("Number of Counts")

print(plot_features)


# cluster marker genes------------------

DefaultAssay(harm.int) <- "SCT"
harm.int <- PrepSCTFindMarkers(object = harm.int, assay  = "SCT")

markers_raw <- FindMarkers(
  object          = harm.int,
  ident.1         = 3,
  only.pos        = TRUE,
  logfc.threshold = 0.25,
  min.pct         = 0.10) %>%
  rownames_to_column("gene")

cluster1_markers <- markers_raw %>%
  filter(p_val_adj < 0.01, pct.1 > 0.10) %>%
  mutate(pct_diff = pct.1 - pct.2) %>%
  arrange(desc(pct_diff), desc(avg_log2FC), p_val_adj)

write.csv(cluster1_markers, "cluster1_positive_markers_logFC0.25_q0.01_pctDiff.csv", row.names = FALSE)


# GO term analysis-------

DefaultAssay(harm.int) <- "RNA"
Idents(harm.int) <- "seurat_clusters"
harm.int <- JoinLayers(harm.int, assay = "RNA")
harm.int <- NormalizeData(harm.int)
harm.int <- FindVariableFeatures(harm.int)
cat("Layers:", Layers(harm.int[["RNA"]]), "\n")
cat("Data dims:", dim(GetAssayData(harm.int, layer = "data")), "\n")

markers <- FindAllMarkers(
  harm.int, only.pos = TRUE, min.pct = 0.1, logfc.threshold = 1.5)
cat("Total markers:", nrow(markers), "\n")

markers_filtered <- markers %>%
  filter(p_val_adj < 0.05, avg_log2FC > 1.5) %>%
  group_by(cluster) %>%
  slice_max(order_by = avg_log2FC, n = 10000) %>%   # limit per cluster
  ungroup()

cat("Filtered markers:", nrow(markers_filtered), "\n")

markers_filtered <- markers_filtered %>%
  mutate(gene_clean = gene %>%
           toupper() %>%
           gsub("-", "", .) %>%
           gsub("\\.", "", .))

gene_map <- bitr(
  markers_filtered$gene_clean,
  fromType = "SYMBOL",
  toType = "TAIR",
  OrgDb = org.At.tair.db)

gene_map_clean <- gene_map %>%
  group_by(SYMBOL) %>%
  filter(n() == 1) %>%
  ungroup()

markers_mapped <- markers_filtered %>%
  left_join(gene_map_clean, by = c("gene_clean" = "SYMBOL"))

markers_mapped <- markers_mapped %>%
  mutate(TAIR_final = case_when(
    grepl("^AT[1-5CM]G\\d+", gene) ~ gene,
    !is.na(TAIR) ~ TAIR,
    TRUE ~ NA_character_))

cat("Total genes:", nrow(markers_mapped), "\n")
cat("Mapped genes:", sum(!is.na(markers_mapped$TAIR_final)), "\n")

markers_clean <- markers_mapped %>%
  filter(!is.na(TAIR_final)) %>%
  distinct(cluster, TAIR_final, .keep_all = TRUE)

markers_clean %>% count(cluster)

all_genes <- rownames(harm.int)
all_genes_clean <- toupper(all_genes) %>%
  gsub("-", "", .) %>%
  gsub("\\.", "", .)

gene_map_bg <- bitr(
  all_genes_clean,
  fromType = "SYMBOL",
  toType = "TAIR",
  OrgDb = org.At.tair.db)

gene_map_bg_clean <- gene_map_bg %>%
  group_by(SYMBOL) %>%
  filter(n() == 1) %>%
  ungroup()

universe <- unique(c(
  gene_map_bg_clean$TAIR,
  all_genes[grepl("^AT[1-5CM]G\\d+", all_genes)]))

cat("Universe size:", length(universe), "\n")
ego_list <- markers_clean %>%
  split(.$cluster) %>%
  lapply(function(df) {
    genes <- unique(df$TAIR_final)
    cat("Cluster", unique(df$cluster), "- genes:", length(genes), "\n")
    if (length(genes) < 10) {
      message("Too few genes, skipping cluster ", unique(df$cluster))
      return(NULL)}
    
    enrichGO(
      gene = genes,
      universe = universe,
      OrgDb = org.At.tair.db,
      keyType = "TAIR",
      ont = "BP",
      pAdjustMethod = "BH",
      qvalueCutoff = 0.05,
      readable = TRUE)})

dotplot_list <- lapply(names(ego_list), function(clust) {
ego <- ego_list[[clust]]
  
  if (!is.null(ego) && nrow(as.data.frame(ego)) > 0) {
    dotplot(ego, showCategory = 1000) +
      ggtitle(paste("Cluster", clust))
  } else {
    message("No significant GO terms for cluster ", clust)
    NULL
  }
})

print(dotplot_list[[12]])

# cluster GO term comparison
ego2  <- as.data.frame(ego_list[["14"]])
ego13 <- as.data.frame(ego_list[["15"]])

cat("Cluster 2 terms:", nrow(ego2), "\n")
cat("Cluster 13 terms:", nrow(ego13), "\n")
terms2  <- ego2$ID
terms13 <- ego13$ID
shared_terms_ids <- intersect(terms2, terms13)
only_2_ids       <- setdiff(terms2, terms13)
only_13_ids      <- setdiff(terms13, terms2)

shared_terms <- ego2 %>%
  dplyr::filter(ID %in% shared_terms_ids) %>%
  dplyr::select(ID, Description, p.adjust)

only_2_terms <- ego2 %>%
  dplyr::filter(ID %in% only_2_ids) %>%
  dplyr::select(ID, Description, p.adjust)

only_13_terms <- ego13 %>%
  dplyr::filter(ID %in% only_13_ids) %>%
  dplyr::select(ID, Description, p.adjust)

cat("Shared terms:", nrow(shared_terms), "\n")
cat("Only cluster 1:", nrow(only_2_terms), "\n")
cat("Only cluster 12:", nrow(only_13_terms), "\n")

print(shared_terms)
print(only_2_terms)
print(only_13_terms)


# embryo and endosperm cluster 13-----
# cluster 13 subcluster analysis
DefaultAssay(harm.int) <- "RNA"
cluster13 <- subset(harm.int, idents = 13)
cluster13 <- NormalizeData(cluster13)
cluster13 <- FindVariableFeatures(cluster13)
cluster13 <- ScaleData(cluster13)
cluster13 <- RunPCA(cluster13)
cluster13 <- FindNeighbors(cluster13, dims = 1:10)
cluster13 <- FindClusters(cluster13, resolution = 0.8)
cluster13 <- RunUMAP(cluster13, dims = 1:10)
my_colors <- c ("#BDDFE7", "#8CC8E0", "#E94D26", "#0B3D91", "#4287BE", "#FABB59", "#FD8D3C")
DimPlot(cluster13, label = TRUE, cols = my_colors, reduction = "umap")

table(cluster13$seurat_clusters)

# subcluster markers
cluster13 <- JoinLayers(cluster13)
cluster13.markers <- FindAllMarkers(cluster13, only.pos = TRUE)
top150.markers <- cluster13.markers %>%
  group_by(cluster) %>%
  slice_max(order_by = p_val_adj, n = 1000)
head(top150.markers)
print(top150.markers, n = 100)

DefaultAssay(cluster13) <- "RNA"
features <- c("IMK2")
seurat_colors <- c("#E0E0E0","#f0f0f0","#FED976","#FEB24C","#FD8D3C","#FC4E2A","#E31A1C","#BD0026") 
p_feature <- FeaturePlot(cluster13, features = features, order = T, min.cutoff = 'q1', max.cutoff = 'q99') + scale_color_gradientn(colors = seurat_colors)
print(p_feature)


# GO term analysis
markers <- FindAllMarkers(cluster13, only.pos = TRUE)
markers_filtered <- markers %>%
  filter(p_val_adj < 0.05) %>%
  arrange(p_val_adj)

ego_list <- markers_filtered %>%
  split(.$cluster) %>%
  lapply(function(df) {enrichGO(
      gene          = df$gene,
      OrgDb         = org.At.tair.db,
      keyType       = "TAIR",   # Araport11 locus IDs (e.g., AT1G01010)
      ont           = "BP",
      pAdjustMethod = "BH",
      qvalueCutoff  = 0.05,
      readable      = TRUE)})

dotplot_list <- lapply(names(ego_list), function(clust) {
  ego <- ego_list[[clust]]
  if(!is.null(ego) && nrow(as.data.frame(ego)) > 0) {
    dotplot(ego, showCategory = 100) + ggtitle(paste("Cluster", clust))
  } else {
    message("No significant GO terms for cluster ", clust)
    NULL}})

print(dotplot_list[[2]])

# compare clusters 
groupA_clusters <- c("0")
groupB_clusters <- c("1")
get_terms <- function(cluster_ids) {
  dfs <- lapply(cluster_ids, function(cl) {
    ego <- ego_list[[cl]]
    
    if (!is.null(ego)) {
      df <- as.data.frame(ego)
      df$cluster <- cl
      return(df)
    } else {
      return(NULL)
    }}) dfs <- do.call(rbind, dfs)
  return(dfs)}

egoA <- get_terms(groupA_clusters)
egoB <- get_terms(groupB_clusters)
cat("Group A terms:", nrow(egoA), "\n")
cat("Group B terms:", nrow(egoB), "\n")
termsA <- unique(egoA$ID)
termsB <- unique(egoB$ID)

shared_ids <- intersect(termsA, termsB)
onlyA_ids  <- setdiff(termsA, termsB)
onlyB_ids  <- setdiff(termsB, termsA)

shared_terms <- egoA %>%
  dplyr::filter(ID %in% shared_ids) %>%
  dplyr::select(ID, Description, p.adjust) %>%
  distinct()

onlyA_terms <- egoA %>%
  dplyr::filter(ID %in% onlyA_ids) %>%
  dplyr::select(ID, Description, p.adjust) %>%
  distinct()

onlyB_terms <- egoB %>%
  dplyr::filter(ID %in% onlyB_ids) %>%
  dplyr::select(ID, Description, p.adjust) %>%
  distinct()

cat("Shared terms:", nrow(shared_terms), "\n")
cat("Only Group A:", nrow(onlyA_terms), "\n")
cat("Only Group B:", nrow(onlyB_terms), "\n")
print(shared_terms)
print(onlyA_terms)
print(onlyB_terms)


# nucellus cluster 18 ------
DefaultAssay(harm.int) <- "RNA"
# Subcluster analysis cluster 18
cluster18 <- subset(harm.int, idents = 18)
cluster18 <- NormalizeData(cluster18)
cluster18 <- FindVariableFeatures(cluster18)
cluster18 <- ScaleData(cluster18)
cluster18 <- RunPCA(cluster18)
cluster18 <- FindNeighbors(cluster18, dims = 1:20)
cluster18 <- FindClusters(cluster18, resolution = 0.5)
cluster18 <- RunUMAP(cluster18, dims = 1:4)
my_colors <- c ("#E94D26", "#FEB24C", "#8B1A1A", "#FD8D3C")
DimPlot(cluster18, label = TRUE, cols = my_colors, reduction = "umap")
table(cluster18$seurat_clusters)

DefaultAssay(cluster18) <- "RNA"

features <- c("AGO3")
seurat_colors <- c("#E0E0E0","#f0f0f0","#FED976","#FEB24C","#FD8D3C","#FC4E2A","#E31A1C","#BD0026") 
p_feature <- FeaturePlot(cluster18, features = features, order = T, min.cutoff = 'q1', max.cutoff = 'q99') + scale_color_gradientn(colors = seurat_colors)
print(p_feature)

# cluster 18 subcluster markers
cluster18 <- JoinLayers(cluster18)
cluster18.markers <- FindAllMarkers(cluster18, only.pos = TRUE)
top150.markers <- cluster18.markers %>%
  group_by(cluster) %>%
  slice_max(order_by = p_val_adj, n = 150)
head(top150.markers)
print(top150.markers, n = 100)


# GO term analysis cluster 18

DefaultAssay(cluster18) <- "RNA"
Idents(cluster18) <- "seurat_clusters"
cluster18 <- JoinLayers(cluster18, assay = "RNA")
cluster18 <- NormalizeData(cluster18)
cluster18 <- FindVariableFeatures(cluster18)
cat("Layers:", Layers(cluster18[["RNA"]]), "\n")
cat("Data dims:", dim(GetAssayData(cluster18, layer = "data")), "\n")
markers <- FindAllMarkers(cluster18, only.pos = TRUE, min.pct = 0.1, logfc.threshold = 1)
cat("Total markers:", nrow(markers), "\n")

markers_filtered <- markers %>%
  filter(p_val_adj < 0.05, avg_log2FC > 1) %>%
  group_by(cluster) %>%
  slice_max(order_by = avg_log2FC, n = 10000) %>%
  ungroup()

cat("Filtered markers:", nrow(markers_filtered), "\n")
markers_filtered <- markers_filtered %>%
  mutate(gene_clean = gene %>%
           toupper() %>%
           gsub("-", "", .) %>%
           gsub("\\.", "", .))
gene_map <- bitr(
  markers_filtered$gene_clean,
  fromType = "SYMBOL",
  toType = "TAIR",
  OrgDb = org.At.tair.db)
gene_map_clean <- gene_map %>%
  group_by(SYMBOL) %>%
  filter(n() == 1) %>%
  ungroup()
markers_mapped <- markers_filtered %>%
  left_join(gene_map_clean, by = c("gene_clean" = "SYMBOL"))
markers_mapped <- markers_mapped %>%
  mutate(TAIR_final = case_when(
    grepl("^AT[1-5CM]G\\d+", gene) ~ gene,
    !is.na(TAIR) ~ TAIR,
    TRUE ~ NA_character_
  ))
cat("Total genes:", nrow(markers_mapped), "\n")
cat("Mapped genes:", sum(!is.na(markers_mapped$TAIR_final)), "\n")
markers_clean <- markers_mapped %>%
  filter(!is.na(TAIR_final)) %>%
  distinct(cluster, TAIR_final, .keep_all = TRUE)
markers_clean %>% count(cluster)
all_genes <- rownames(cluster18)

all_genes_clean <- toupper(all_genes) %>%
  gsub("-", "", .) %>%
  gsub("\\.", "", .)

gene_map_bg <- bitr(
  all_genes_clean,
  fromType = "SYMBOL",
  toType = "TAIR",
  OrgDb = org.At.tair.db)

gene_map_bg_clean <- gene_map_bg %>%
  group_by(SYMBOL) %>%
  filter(n() == 1) %>%
  ungroup()

universe <- unique(c(
  gene_map_bg_clean$TAIR,
  all_genes[grepl("^AT[1-5CM]G\\d+", all_genes)]
))

cat("Universe size:", length(universe), "\n")
ego_list <- markers_clean %>%
  split(.$cluster) %>%
  lapply(function(df) {
    genes <- unique(df$TAIR_final)
    cat("Cluster", unique(df$cluster), "- genes:", length(genes), "\n")
    if (length(genes) < 10) {
      message("Too few genes, skipping cluster ", unique(df$cluster))
      return(NULL)
    }
    
    enrichGO(
      gene = genes,
      universe = universe,
      OrgDb = org.At.tair.db,
      keyType = "TAIR",
      ont = "BP",
      pAdjustMethod = "BH",
      qvalueCutoff = 0.05,
      readable = TRUE
    )
  })

dotplot_list <- lapply(names(ego_list), function(clust) {
  ego <- ego_list[[clust]]
  
  if (!is.null(ego) && nrow(as.data.frame(ego)) > 0) {
    dotplot(ego, showCategory = 100) +
      ggtitle(paste("Cluster", clust))
  } else {
    message("No significant GO terms for cluster ", clust)
    NULL
  }
})

print(dotplot_list[[3]])

# Subcluster GO term comparison
ego2  <- as.data.frame(ego_list[["0"]])
ego13 <- as.data.frame(ego_list[["2"]])
cat("Cluster 2 terms:", nrow(ego2), "\n")
cat("Cluster 13 terms:", nrow(ego13), "\n")
terms2  <- ego2$ID
terms13 <- ego13$ID

shared_terms_ids <- intersect(terms2, terms13)
only_2_ids       <- setdiff(terms2, terms13)
only_13_ids      <- setdiff(terms13, terms2)

shared_terms <- ego2 %>%
  dplyr::filter(ID %in% shared_terms_ids) %>%
  dplyr::select(ID, Description, p.adjust)

only_2_terms <- ego2 %>%
  dplyr::filter(ID %in% only_2_ids) %>%
  dplyr::select(ID, Description, p.adjust)

only_13_terms <- ego13 %>%
  dplyr::filter(ID %in% only_13_ids) %>%
  dplyr::select(ID, Description, p.adjust)

cat("Shared terms:", nrow(shared_terms), "\n")
cat("Only cluster 1:", nrow(only_2_terms), "\n")
cat("Only cluster 12:", nrow(only_13_terms), "\n")

print(shared_terms)
print(only_2_terms)
print(only_13_terms)


# abscission zone script --------------------

# Subcluster analysis cluster 17
cluster17 <- subset(harm.int, idents = 17)
DefaultAssay(cluster17) <- "RNA"
cluster17 <- NormalizeData(cluster17)
cluster17 <- FindVariableFeatures(cluster17)
cluster17 <- ScaleData(cluster17)
cluster17 <- RunPCA(cluster17)
cluster17 <- FindNeighbors(cluster17, dims = 1:3)
cluster17 <- FindClusters(cluster17, resolution = 0.5)
cluster17 <- RunUMAP(cluster17, dims = 1:3)
my_colors <- c("#80C8E0", "#F78F44", "#4287BE", "#F9C464")
DimPlot(cluster17, label = TRUE, cols = my_colors, reduction = "umap")

cluster17 <- JoinLayers(cluster17)

cluster17.markers <- FindMarkers(cluster17, ident.1 = 3, only.pos = TRUE)
library(dplyr)
top150_cluster17 <- cluster17.markers %>%
  arrange(p_val_adj) %>%
  slice_head(n = 200)
head(top150_cluster17, 200)  


features <- c("HAE")
seurat_colors <- c("#E0E0E0","#f0f0f0","#FED976","#FEB24C","#FD8D3C","#FC4E2A","#E31A1C","#BD0026") 
p_feature <- FeaturePlot(cluster17, features = features, order = T, min.cutoff = 'q1', max.cutoff = 'q99') + scale_color_gradientn(colors = seurat_colors)
print(p_feature)


#comparison Lee et al. 2018 secession and residuum top 500 DEGs sorted after adjusted p-value
#supplementary file of Lee et al. 2018 was sorted after only secession/just residuum cell expressed, then sorted after adjusted p-value and inserted as "study_genes" vectors
#secession cell markers
library(RColorBrewer)

study_genes <- c("AT1G01240","MPK11","AT1G01725","AT1G01770","NAC003","MAN1","GPAT2","AT1G02660","GRXS13","AT1G05340","UGT75B1","AT1G05562","GPAT1","AT1G06760","AT1G07160","ECA4","FATB","ESL1","AT1G09250","AT1G09500","GLP5","AT1G10040","AT1G10050","AT1G10160","NFXL1","ARR4","AT1G10682","AT1G11170","MLO2","AT1G12320","AT1G12760","AFB3","AT1G13340","AT1G13360","AT1G13520","AT1G13880","AT1G14040","AT1G14600","AT1G14780","GLYI4","AT1G15385","AT1G15400","ABCG40","SRG1","CYP72C1","GSTU25","AT1G17340","AT1G18470","AT1G19200","DA1","MP","AT1G20100","SAUL1","AT1G22180","AT1G22190","UGT85A1","AT1G22882","CRF7","MDP40","FH4","AT1G25390","AT1G26920","STZ","GH3.17","AT1G28760", 
                 "AT1G29290","RKF1","AT1G29800","GA2OX2","MUR4","AT1G30700","SGR2","XLG3","SBT3.5","AT1G34220","PHI-1","AT1G35190","TPPD","SHM7","5PTASE11","NTMC2T5.1","AT1G50590","AT1G51790","IOS1","LEA7","NAC019","PES1","Cand1","PUP18","CAT2","MSL3","ZF14","AT1G59660","IRT3","BGLU46","AS2","AT1G66090","TMK1","AT1G66480","AT1G67580","AT1G67850","AT1G68140","ENDO2","AT1G68410","AT1G68620","CLE12","AT1G68820","AT1G69570","AT1G69572","MMP","TPS8","CRK3","AT1G70790","AT1G73240","AT1G73480","AT1G73750","AT1G74010","SS2","AT1G75000","AT1G75340","CKX5","HMG1","TLP1","EXT4","SKP2B","AT1G77220","AT1G77770","AT1G77840","RAP2.4","AT1G78280","ALIS5","NTT1","RAE1","AT1G80940",
                 "PAP1","AT2G01460","AT2G02370","AT2G03200","AT2G05910","ADF5","AT2G17500","AT2G17760","AT2G18690","TPS11","AT2G20320","RD2","AGP2","PSK2","AT2G23450","TET8","LRL1","AT2G25450","AT2G25460","EBF1","AT2G25625","ARSK1","AR781","PLA2A","AT2G27389","AT2G27390","AT2G27500","ARF10","AT2G28400","AT2G28690","AT2G28710","AT2G29065","GSTU4","AT2G30020","MAPKKK14","ASL5","AT2G30550","APG9","AT2G31345","AT2G31945","AT2G32190","AP4.3A","PP2CG1","AT2G33710","AT2G35390","AT2G36220","AT2G37150","ZAT11","AT2G38000","AT2G38240","AT2G38250","PRA1.B4","AGT3","AT2G38410","WRKY33","AT2G38800","ABCG1","AT2G39520","AT2G39530","AT2G39650","BIK1","HSPRO2","AT2G40095","PP2C5",
                 "AT2G41380","AT2G41410","AT2G41835","AT2G42270","ML2","IBH1","MSS3","AT2G44130","AT2G44140","AT2G44500","UCC2","PI4K GAMMA 4","HB-7","AT2G47550","AT2G47560","HIR2","DOX1","AT3G02030","SRG3","AT3G02880","XPO1B","LCAT3","EDA5","NHX2","AT3G05165","AT3G05390","AT3G05670","APC11","RCI2A","AT3G05937","AT3G05940","TLP9","ATG8H","MYB108","IBR3","AT3G07340","AIR12","AT3G07470","OBE1","QRT2","AT3G08650","AT3G09010","GSTU8","NAC052","BZIP28","AT3G10815","NHL1","AT3G11760","NF-YC11","AT3G12510","AT3G12640","NANA","AT3G13275","AT3G13310","BRL3","AT3G13650","RSH2","CAX9","NTMC2T6.2","CYP72A15","AT3G15430","NAC3","MLP-300B","COBL8","AAE7","CTL2","AT3G17770","PAP17",
                 "AT3G18250","AT3G18950","AT3G19240","AHP1","AT3G21530","AT3G21680","AT3G22060","OASA2","AT3G22600","AT3G22910","IAA2","AT3G23170","AT3G23570","AT3G24460","PERK1","AGC2-1","AT3G25610","ABCG21","AT3G26100","TBL22","PMZ","GATL10","AT3G28850","AT3G29000","SDR4","AT3G42180","NIT2","AT3G45040","AT3G46080","AT3G46930","G3Pp1","ABCA2","ABCA7","AT3G48060","SPT1","AT3G49540","ATG13","PSK4","AT3G50910","AT3G51110","AT3G51660","UGT73D1","AtCDC48B","CHX20","AT3G54200","BME3","ABCG16","ABCG17","AT3G55430","AT3G55470","PROT2","AT3G55890","AT3G55900","CAN","AT3G56360","AGK2","CSY2","IQD13","AT3G59695","BGLU16","BAP1","ABCB21","GATL7","XT1",
                 "AT4G02940","Tic20-IV","SEC1B","NYC1","KMS1","LigB","AT4G15120","AT4G15610","AT4G16146","ACX1","ERF6","AT4G17790","AT4G18340","AtS40-3","XTH29","IWS2","AT4G19200","AT4G19670","AT4G20320","AT4G20460","AAT1","AT4G22360","AT4G22470","NYE1","NLP7","UBP16","NBR1","AT4G25170","AT4G25390","SLK1","XTR6","AT4G25940","ABI1","PFK3","WES1","AT4G27280","AT4G27680","ACBP2","AT4G28300","ADS1","AT4G29780","HA2","XTH24","AT4G30390","AT4G30400","AT4G30790","AT4G32250","AT4G32440","AT4G32920","AT4G33150","AT4G33160","AT4G33420","AT4G33920","CER9","UGT73B2","CAD5","MYB32","AT4G35985","J11","BEH2","AT4G36790","AT4G37290","AT4G37520","AT4G37900","ELI3-2","AT4G39780",
                 "AT5G01380","LECRKA4.1","AT5G01715","AT5G01720","SR1","AT5G03110","AT5G03330","AT5G03610","AT5G03810","SDP1","MTM2","ALA1","DREB2A","ASA1","AT5G06370","ML4","GDH2","AT5G08240","AT5G08660","AT5G08780","AT5G10650","AT5G11090","GLR2.5","NADP-ME2","AT5G12010","Rap2.6L","AT5G13370","PPH","TBP1","ELC-Like","AT5G13880","RGLG2","AT5G15640","AT5G16380","AT5G16960","AER","AT5G17350","AT5G17650","AT5G19070","RAN-1","EFR","SUS1","AT5G21940","AT5G22850","AT5G23340","AT5G25770","AT5G25930","DPD1","CNI1","PNC2","LACS7","AT5G35110","AT5G35735","AT5G38900","PMAT1","BIP2","AT5G43570","UPI","MAP18","APG7","AT5G46080","RBOHD","TT10","NSP5","GLR1.3","WRKY48","AT5G49665",
                 "ERD1","ORG1","AT5G53820","PSKR2","AT5G54300","DFL1","RAT5","Fh5","AT5G54660","AT5G55960","NDL1","AT5G56980","AT5G57010","VIN3","SIP1","EXO70B1","AT5G59530","RHL41","AT5G60710","AT5G61390","AT5G61820","ELF5","AT5G63130","AT5G64190","AT5G64230","AGP1","CMPG2","ACX2","TPPJ","AT5G65290","AT5G65300","AT5G65380","GATA5","AT5G66810","CYP81G1","AT5G67350","WIH1")

genes_present <- study_genes[study_genes %in% rownames(harm.int)]
cluster_cells <- WhichCells(harm.int, idents = "17")
expr_data <- GetAssayData(harm.int, layer = "data")[genes_present, cluster_cells, drop = FALSE]
pct_expressed <- rowSums(expr_data > 1) / length(cluster_cells)
expressed_genes <- names(pct_expressed[pct_expressed >= 0.05])
n_detected <- length(intersect(study_genes, expressed_genes))
n_total <- length(study_genes)
detected_pct <- round(100 * n_detected / n_total, 1)

present_genes <- intersect(study_genes, rownames(cluster17))
module_genes_filtered <- list(present_genes)
print(present_genes)
cat("number of genes found in Seurat object:", length(present_genes), "\n")
if (length(present_genes) >= 5) {
  cluster17 <- AddModuleScore(object = cluster17, features = module_genes_filtered, name = "secession_module")
  cat("module score added. Metadata column: secession_module1\n")

# plot as feature plot or violin plot
rdylbu_colors <- rev(brewer.pal(11, "RdYlBu"))
rdylbu_trimmed <- rdylbu_colors[-c(1:3)]
FeaturePlot(cluster17, features = "secession_module1", reduction = "umap", cols = rdylbu_trimmed) + ggtitle("secession score")
VlnPlot(cluster17, features = "secession_module1", group.by = "seurat_clusters", pt.size = 0.1, cols = my_colors) + ggtitle("secession score")

} else {
  cat("Not enough genes found to calculate a reliable module score.\n")
}

# residuum cell markers
study_genes <- c("PPa1", "AT1G01100","GPAT4","AT1G01920","AT1G02190","CER1","emb2386","AT1G04040","RPS15","c-NAD-MDH1","AT1G04480","STO","PSBP-1","AT1G07210","RPS15A","AT1G07940","AT1G08360","MGP2","AT1G09590","AT1G09620","AT1G09690","SEX1","AT1G11860","AT1G12230","AT1G13930","AT1G14820","AT1G15120","AT1G15930","AT1G16000","AT1G16350","RDH2","RABA1b","HLL","ABCG11","ATARCA","AT1G18540","ARR7","DHAR1","APS2","HYD1","DRT112","OPCL1","AT1G20950","AT1G21440","PFL","RPL27AB","AT1G26550","AT1G26880","AT1G27400","AT1G27435","APT1","LTPG1","AT1G29250","AT1G29670","AT1G29880","eEF-1Bb1","PSAK","IMD3","PSAF","PRK","AT1G32580","PRPL11","PGY2","AT1G34030","EMB3003","RPL21C",
                 "AT1G41880","GAPB","RP1","NPQ4","EMB3105","D-CDES","RACK1B_AT","AT1G48750","AT1G48830","NUC-L1","TOM6","AT1G52300","mMDH1","ACP2","TLP18.3","CPN60B","NOP56","AT1G57660","AT1G57720","AT1G57860","AT1G60010","FED A","RPL3B","NRS/ER","AT1G63610","LACS3","AT1G65150","mtACP2","AT1G65295","AT1G65370","GAD2","AT1G66430","MIR163","RBCS1A","AT1G67430","AT1G67700","HPR","XYL1","AP1","RPL34","HVA22C","SPDS2","BCA4","AT1G70600","GH9B1","ATWHY2","P40","LPA3","AT1G73940","AT1G74050","AT1G74060","AT1G74270","GASA6","ALDH10A8","GH9B7","TUB1","AT1G77940","CYP708A3","LUP2","PSBR","AT2G01250","AT2G01720","EMB2816","CR88","AT2G04700","AT2G05220","MOD1","AT2G06950",
                 "AT2G17360","EMB2296","PPa2","AT2G18400","AT2G19680","AT2G19730","AT2G19740","AT2G19940","AT2G20450","PSB29","AT2G20930","TIM","AT2G21250","AT2G21580","RNR1","AT2G22230","MIPS2","PAB4","SNRNP-G","AT2G24280","AT2G25210","AT2G26500","PGY1","AT2G27680","AT2G27710","AT2G27720","EXPA6","AT2G29510","TIM10","AT2G30060","PSBW","GSTF10","AT2G32060","AT2G32580","HSP60-2","AT2G33370","AT2G34357","LHB1B2","AT2G34460","AT2G34480","GDCH","AT2G35760","AT2G36160","AT2G36170","GAMMA-TIP","PIP2B","RD28","AT2G37190","RPS5B","AT2G37600","AKR4C10","LP1","AT2G39020","ATATH8","RPL23AA","RCA","AT2G40205","AT2G40590","AT2G40660","PPC2","AT2G43460","AT2G43780","LARP6b",
                 "COS1","AT2G44060","AT2G45030","AT2G46535","PSI-P","UBQ6","AT2G47115","LACS1","AT2G47610","AT3G01790","AT3G02080","AT3G02560","UGP1","AT3G03920","AT3G03960","emb2171","AT3G04920","AT3G05060","AT3G05560","RPL18","AT3G06680","AT3G06700","PRMT4B","AT3G07110","AT3G08030","AT3G08740","AT3G09200","AT3G09500","AT3G09630","ADK1","AT3G09860","AT3G10090","GLY2","AT3G11120","AT3G11250","AT3G11500","AT3G11510","AT3G11830","RPS5A","AT3G12345","PGK1","AT3G13882","AT3G14380","AT3G14600","AT3G16080","AT3G16370","NSP1","HS1","TOM40","TCP-1","CYP705A19","GRP5","PUR7","AT3G22230","TSO1","AT3G23390","HSP60","AT3G24420","AT3G24830","ATL5","GLYR1","AT3G25640","LTA2",
                 "RPL15","PRXQ","AT3G26070","TIP2","GAPA","AT3G27230","MYB21","RPL12-A","AT3G28500","AT3G28900","AT3G28940","PHS1","AT3G43690","CAM7","AT3G43980","AT3G44010","HDA3","RPL9","AT3G45030","RPS15AD","TIM9","AT3G47070","AT3G47370","AT3G47380","AT3G47650","CDC2","EMB1080","BBC1","NACA2","AT3G49910","NOXY38","PSRP2","UBQ1","STV1","PIP2A","AT3G53430","AT3G53740","AT3G53870","AT3G53890","TBL36","LHCA1","AT3G55170","RPL23AB","CER10","TPI","AT3G55750","ROC2","AT3G56130","AT3G56340","PSRP5","CRD1","CBL","NAP57","ADSS","AT3G58700","AT3G59540","AT3G60245","AT3G60770","RS27A","HDG1","SDR1","LHCA2","ROC4","AT3G62120","UBQ5","AT3G62530","AT3G62870","APG1","CCD1",
                 "RPS13A","CYP86A2", "AT4G00810","AT4G01150","AT4G01310","SAM-2","AT4G02930","AT4G03200","PETC","PP2C52","ATPC1","KNAT1","NDPK1","RPS18C","LHCB5","AT4G10450","NDPK3","PSAL","ENODL19","AT4G13170","GR-RBP2","AT4G14320","HCD1","AT4G15000","AT4G15620","AT4G15640","AT4G15930","AT4G16155","AT4G16720","AT4G17390","AT4G18100","CHLI1","RPL16B","AT4G21620","PRXR1","AT4G24830","ACP4","AT4G25740","AT4G25890","NAP1;1","AT4G26230","AT4G27090","AT4G29410","AT4G30330","AT4G30800","LAP2","RPS6","GPX7","AT4G31985","ASP5","PDE334","AT4G32330","SHM3","AGD2","CER4","AT4G33865","SEP1","AT4G34670","PIP3","ALDH3F1","BGAL3","AT4G36660","AT4G37220","SHM1","AT4G39040",
                 "AT4G39200","CAD9","AT5G02450","AT5G02540","AT5G02610","AT5G02870","AT5G02960","AT5G04800","AT5G07090","AT5G08050","AT5G08180","STRS2","AT5G08690","PPa6","EMB3010","RING1","AT5G10400","AT5G10920","TUB6","AT5G12470","GUN5","EMB3137","AT5G14920","AT5G15200","BCCP2","AT5G16060","AT5G16070","AT5G16130","AT5G17700","PSRP6","AT5G18380","AT5G19500","AT5G19510","AT5G19760","SPS1F","AT5G20290","CPN20","AT5G20890","AT5G20950","AT5G22440","HD2B","HCF136","RPS11-BETA","AT5G23970","VSP2","VSP1","CRY3","AT5G26110","AT5G26800","AT5G27120","AT5G27700","AT5G27770","AT5G27850","AT5G28060","RPS1","AT5G35530","GS2","AT5G36210","AT5G37540","GRF3","RPL5B","UK/UPRT1",
                 "RPS10B","TOM22-V","AT5G44568","AT5G45500","AT5G45775","APE2","AT5G47190","AT5G47550","AT5G47700","AT5G47930","FKBP15-2","AT5G48760","MMT","TIM8","FIB1","PKP-BETA1","AT5G53070","AT5G53880","LHCB3","AT5G55125","MAP65-1","PSL4","AT5G56670","AT5G56710","NAP1;3","ORP4C","GS1","AT5G57700","CER3","PHOT2","AT5G58330","AT5G58420","AT5G59250","LTP4","LTP3","AT5G59850","AT5G60390","AT5G60670","AT5G61170","GCD1","ADK1","FLS2","FNR1","PSBO1")

genes_present <- study_genes[study_genes %in% rownames(harm.int)]
cluster_cells <- WhichCells(harm.int, idents = "17")
expr_data <- GetAssayData(harm.int, layer = "data")[genes_present, cluster_cells, drop = FALSE]
pct_expressed <- rowSums(expr_data > 1) / length(cluster_cells)
expressed_genes <- names(pct_expressed[pct_expressed >= 0.05])
n_detected <- length(intersect(study_genes, expressed_genes))
n_total <- length(study_genes)
detected_pct <- round(100 * n_detected / n_total, 1)

present_genes <- intersect(study_genes, rownames(cluster17))
module_genes_filtered <- list(present_genes)
print(present_genes)
cat("number of genes found in Seurat object:", length(present_genes), "\n")
if (length(present_genes) >= 5) {cluster17 <- AddModuleScore(object = cluster17, features = module_genes_filtered, name = "residuum_module")
cat("module score added. Metadata column: residuum_module1\n")

# plot as feature plot or violin plot  
rdylbu_colors <- rev(brewer.pal(11, "RdYlBu"))
rdylbu_trimmed <- rdylbu_colors[-c(1:3)]
FeaturePlot(cluster17, features = "residuum_module1", reduction = "umap", cols = rdylbu_trimmed) + ggtitle("residuum score")
VlnPlot(cluster17, features = "residuum_module1", group.by = "seurat_clusters", pt.size = 0.1, cols = my_colors) + ggtitle("residuum score")
  
} else {
  cat("Not enough genes found to calculate a reliable module score.\n")
}


# floral abscission differentiation state comparison Wen et al. 2025
# state markers from Wen et al. 2025 were copied and inserted into "study_genes" vectors
library(eulerr)
library(grid)

#state 1 markers
study_genes <- c("AT5G60670","RBCS1B","RBCS2B","PSBX","PSBTN","FED A","AT4G01150","LHCA1","RCA","FBA2","PSBO1","LHCB5","PSAF","TUFA","CAB2","AT2G10940","ROC4","CRB","PSAG","LHCB2.2","CAB1","CAB3","RBCS3B","RBCS1A","CA1","GAPA-2","LHCB3","PSAN","LHCB6","PSBP-1")
#state 2 markers
study_genes <- c("ARFA1E", "TCTP", "AT5G16590", "AT4G20830", "TRX5", "PSK3", "NADP-ME2", "AT1G67920", "AT2G38800", "AT5G11970", "AT1G19020", "SAG24", "AT5G54165", "GPX6", "AT4G36990", "AT4G36648", "AT5G10695", 
                 "RHL41", "AT5G42300", "AT3G45180", "AT2G37110", "MYB74", "AT5G35735", "DI19", "AT5G42050", "NHL3", "AT1G68440", "GSTU19", "ATHCSTM1", "ELF5A-3", "ERD14", "AT5G01750")
#state 3 markers
study_genes <- c("CER1", "LTPG2", "PRP2", "GLC", "LACS2", "APG", "SHN3", "PEL3", "AT5G45670", "LTL1", "AT4G28780", "AT4G18970", "AT5G33370", "AT1G45010", "PRP4", "CAD9", "AT3G23450", "EXL3", "GRP23", "GPAT1", 
                 "KCS1", "HHP1", "CER4", "AT1G55260", "AT1G29670", "ALDH3F1", "LACS1", "KCS20", "PME44", "GPAT4")

genes_in_data <- rownames(harm.int)
present_genes <- intersect(study_genes, genes_in_data)
missing_genes <- setdiff(study_genes, genes_in_data)
length(study_genes)     
length(present_genes)    
length(missing_genes)   

genes_present <- study_genes[study_genes %in% rownames(harm.int)]
cluster_cells <- WhichCells(harm.int, idents = "17")
expr_data <- GetAssayData(harm.int, layer = "data")[genes_present, cluster_cells, drop = FALSE]
pct_expressed <- rowSums(expr_data > 2) / length(cluster_cells)
expressed_genes <- names(pct_expressed[pct_expressed >= 0.10])
n_detected <- length(intersect(study_genes, expressed_genes))
n_total <- length(study_genes)
detected_pct <- round(100 * n_detected / n_total, 1)
sets <- list("floral AZ" = study_genes, "seed AZ" = expressed_genes)
fit <- euler(sets)
plot(fit, fills = list(fill = c("skyblue", "#FDAE61FF"), alpha = 0.5),
     edges = TRUE, labels = list(font = 1.2, cex = 1.2), quantities = list(font = 1.2, cex = 1.2), main = "state (insert state) floral abscission markers in seed AZ", main.cex = 1.2)
grid.text(paste0(n_detected, "/", n_total, "floral AZ genes detected in seed AZ (", detected_pct, "%)"), x = 0.5, y = 0.1, gp = gpar(fontsize = 10, fontface = "italic"))


# floral abscission transcription factor comparison Wen et al. 2025 with "marker" genes of cluster 17 classified/predicted as TFs
#find markers with only slight filtering to reduce noise was performed. Markers were used as an input for TF domain prediction on Araport (Protein domain enrichment, no test correction, max p-value 1)
DefaultAssay(harm.int) <- "integrated"
markers_17 <- FindMarkers(object = harm.int, ident.1 = 17, min.pct = 0.05, logfc.threshold = 1, only.pos = TRUE)
print(markers_17)

markers_17$gene <- rownames(markers_17)
write.csv(markers_17, file = "cluster17_integrated_input_for_araport.csv", row.names = FALSE)

# 67 (predicted) TFs in seed AZ cluster 17
cluster_genes <- c("OBP2","AT1G10585","DDF1","WRKY4", "AT1G19210","DREB26", "AT1G22190","AT1G22810","STZ","ERF11", "RAP2.6","NAC019","ERF8", "WRKY6","AT1G64380","RAV2","HB18",
                   "AT1G72210", "ORA47", "RAP2.4","WRKY40","HB17","RGF3","WRKY15","WRKY33", "CZF1","AT2G44940","WRKY46","WRKY23","ERF71","MBD11","ZF2","MYB15", "EDF3","MYB94",
                   "CEJ1","AT3G50650","BME3","SZF1","MYB11","MYB74", "AT4G17670","WRKY28","CBF1","AT4G28140","RRTF1","NAC076","HSF4","ZAT6", "ATAF2","WRKY75", "NAC084",
                   "AT5G21960","BZIP9","BHLH92", "WRKY8","HAT2","WRKY48","ERF104")

# TFs of floral AZ, Wen et al 2025
study_genes <- c("PRE1", "TCP11", "ERF21", "ZF2", "ZAT6", "AT1G22810", "MYB7", "ZAT11", "HAT22", "RD26", "DEAR3", "WRKY75", "MYB74", "PMZ", "AT1G66500", "AT5G43620", "RHL41", 
                 "AT1G26800", "HSF4", "HSF A4A", "WRKY15", "NAC3", "CEJ1", "AT1G63840", "CRF7", "MYB62", "GBF6", "AT1G19210", "ERF11", "MYB96", "ATML1", "SHN3", "PDF2")

sets <- list("floral AZ TFs" = study_genes, "seed AZ predicted TFs" = cluster_genes)
fit <- euler(sets)
plot(fit, fills = list(fill = c("grey", "#F78F44"), alpha = 0.5), labels = list(font = 2, cex = 1.2), quantities = list(font = 1.2, cex = 1.2), main = "AZ TFs", main.cex = 1.4)

#TFs present in flower and seed specific datasets, overlap of both
intersect(study_genes, cluster_genes)
floral_AZ_only <- setdiff(study_genes, cluster_genes)
floral_AZ_only
seed_AZ_only <- setdiff(cluster_genes, study_genes)
seed_AZ_only
```
